# Comprehensive transcriptomic profiling uncovers an inflammatory maturation program conserved in human and mouse dendritic cells

**DOI:** 10.64898/2026.07.30.741848

**Authors:** Jessica Bourque, Robert Kousnetsov, Daniel Hawiger

**Affiliations:** Department of Molecular Microbiology and Immunology, Saint Louis University School of Medicine, St. Louis, MO, USA; Department of Medicine, Massachusetts General Hospital, Boston, MA, USA

**Author notes:** These authors contributed equally (Jessica Bourque, Robert Kousnetsov).

**Keywords:** Dendritic cells, scRNA-seq, transcriptomic, Seqtometry, maturation, tolerance, inflammation, Type I interferon, human disease

## Abstract

Conventional dendritic cells (cDCs) integrate signals to balance tolerance and immunity, but how steady-state cDC programs are changed during inflammatory maturation remains incompletely resolved. Here, using Seqtometry-based analysis of mouse and human cDCs, we identify distinct steady-state cDC1 gene programs enriched for either tolerance-associated or immune response-associated (pre-immunogenic) transcriptomic features that are present under homeostatic conditions in fully differentiated cDCs with divergent predicted immune functions. Under inflammatory conditions, the tolerance-linked features are reduced, whereas the pre-immunogenic program is extended in response to type I interferon signaling and is selectively impaired by *Ifnar1* deficiency. This core inflammatory program is conserved across mouse and human cDC subsets, and while retaining disease-specific transcriptomic features, it is detected in infection, cancer, and autoimmunity. Together, these findings establish a gene program-based framework for cDC inflammatory maturation that extends beyond a binary immature-versus-mature classification and supports the identification of disease-associated cDC biomarkers.

## INTRODUCTION

Conventional dendritic cells (cDCs) are essential regulators of adaptive immunity, integrating innate recognition with T cell priming.^1^ They arise from bone marrow progenitors in a FMS-like tyrosine kinase 3 ligand (Flt3L)-dependent manner before seeding peripheral and lymphoid tissues, where they differentiate into XCR1⁺ type 1 cDCs (cDC1s) and SIRPα⁺ type 2 cDCs (cDC2s).^2,3^ cDC1s prime both CD4^+^ and CD8^+^ T cells, which are critical in initiating anti-tumor and anti-viral immune responses, whereas cDC2s may prime predominantly CD4+ T cells.^4,5^ Moreover, some cDCs, mainly of the cDC1 subset, also play a contrasting tolerogenic function, primarily by converting antigen-specific CD4^+^ T cells into peripherally induced, Foxp3-expressing regulatory T cells (pTregs) that negatively modulate various immune and autoimmune responses.^6–8^ How cDCs carry out diverse and often opposing immune functions remains a topic of intense investigation. In general, cDCs determine the specific fates of antigen-responding T cells and their functional differentiation by integrating multiple extrinsic signals and undergoing complex changes, including expressing various immunomodulatory proteins on their surface and secreting required cytokines. This process, termed “maturation” or “activation,” differs from cDC development^2,9^ and confers specific immunogenic functions to cDCs. The prototypical maturation process is initiated when cDCs are exposed to canonical pathogen-associated molecular patterns (PAMPs) and specific cytokines. These comprehensive functions of cDCs depend on their short lifetime and a quick turnover that allow specific cDCs to undergo abrupt and profound maturation changes in response to changing immune conditions and then rapidly reset base populations of cDCs upon a return to homeostasis.^8^ Therefore, studying the molecular mechanisms distinguishing the dynamic processes of maturation of cDCs under homeostatic and inflammatory conditions is challenging.

Among the cytokines, type I interferon (IFN-I)^9^ has been shown to be crucial for cDC maturation by upregulating specific maturation pathways in response to PAMPs.^10–14^ IFN-I-induced genes have also been shown to increase the immunogenicity of cDCs following stimulation with polyinosinic:polycytidylic acid (poly I:C), a model toll-like receptor (TLR) ligand.^10^ Further, IFN-I-dependent responses, in concert with signals from commensal bacteria, have been proposed to maintain homeostatic functions of cDCs in the gut-associated lymphoid organs (GALT).^15,16^ While IFN-I is key in remodeling the cDC functional landscape, it remains unresolved whether IFN-I drives a comprehensive, conserved maturation program across subsets and species that integrates both steady-state and inflammatory contexts.

While the developmental pathways are well defined, the molecular programs that regulate cDC maturation under inflammatory conditions remain incompletely understood. Classical models emphasize “immunogenic” versus “tolerogenic” maturation, but these frameworks rely on narrow protein markers and context-dependent functional readouts.^17^ A relevant homeostatic activation is an endogenous process that does not rely on extrinsic proinflammatory signals. The intrinsic signals that have been recently proposed to be involved include cholesterol uptake, cell-shape sensing, and erythropoietin receptor (EPOR) functions.^17–20^ In the steady state, some cDCs acquire mixed gene expression programs resembling both pro-immunogenic and tolerogenic states, a process collectively termed “homeostatic maturation.”^17,21,22^ For example, these cDCs express pro-immunogenic genes, such as *Cd40*, *Ccr7*, *Il12b* while also expressing the tolerogenic gene *Cd274.*^17,22^ In addition to responses mediated by physiological signals, upon uptake of tumor cells, cDCs can undergo very similar changes and become “mature DCs enriched in immunoregulatory molecules”(mregDCs)^23,24^ via a mechanism that may also depend on cholesterol metabolism.^19,25^ How these homeostatic processes relate to subsequent inflammatory maturation of cDCs remains unresolved.

A given biological process is associated with multiple genes whose expression is specifically altered. Therefore, a defined group (set) of genes constitutes a precise molecular signature of a corresponding biological process surpassing the conventional reliance on individual biological markers.^26,27^ We recently developed a gene signature-based approach, termed “Seqtometry,” a single-cell analytical strategy using an advanced scoring method to test multiple genes pertinent to specific biological processes.^26,28^ By bypassing unsupervised clustering and instead focusing on information from specific gene signatures, Seqtometry pinpoints populations among all other cells in a given data set based on multiple specifically-defined characteristics. Subsequently, such populations can be subject to a rigorous comparative analysis within a collection of data sets resulting in the faithful and comprehensive characterization of biological processes under investigation, either in experimental or disease conditions.^26,28^ To directly address the challenge of studying the molecular mechanisms distinguishing the maturation of cDCs under homeostatic and inflammatory conditions, we systematically dissected cDC maturation across mice, humans, and disease contexts. We identified a conserved immune response-associated maturation program that is established under homeostatic conditions in a defined group of cDCs and is orchestrated by IFN-I to specifically expand under various inflammatory conditions. This program lacks tolerance-linked molecular features and is shared across multiple cDC subsets. Overall, our findings define the inflammatory maturation program as a conserved program of dendritic cell biology across multiple inflammatory and disease states including infection, cancer, and autoimmunity. By demonstrating that IFN-I governs this program linking innate sensing to adaptive immune priming, we further establish a molecular framework for understanding cDC responses and highlight inflammatory dendritic cells as potential biomarkers and therapeutic targets.

## RESULTS

### Steady-state cDC1s contain distinct *Tolerogenic* and *Pre-immunogenic* transcriptomic programs

To comprehensively characterize cDC1s in homeostatic (steady-state) conditions, we isolated splenocytes from wild-type (WT) mice corresponding to the cDC1 subset^29^ and performed single-cell RNA sequencing (scRNA-seq) analysis (Figure S1A). Since cDCs can lose expression of subset-specific markers upon activation,^17,30^ this approach allowed for the inclusion of cDC1s that may have undergone activation-associated transcriptomic changes while retaining XCR1 protein, a hallmark marker of murine cDC1s.^31^ We first identified proliferating cDC1s, as reported previously,^30,32^ by expression of *Mki67* (encoding Ki-67), a marker of proliferating or recently divided cells^33^ (Figure 1A). The decreasing transcriptomic diversity of individual cells within a given cell type is a hallmark of an advancing differentiation process, and such differentiation states can be predicted from CytoTRACE scores calculated from scRNA-seq data.^34^ To estimate differentiation status without relying on predefined clusters, we calculated inverted CytoTRACE scores for all cells within our splenic cDC1 dataset (see Methods). Using this CytoTRACE score metric, with the lowest scores corresponding to the least differentiated cells and higher values corresponding to more differentiated cells, we observed a gradient of transcriptomic differentiation across the cDC1 compartment (Figure 1B). As expected, *Mki67*-expressing cells were concentrated among the least differentiated cells, whereas other cDC1s extended toward more differentiated transcriptomic states (Figures 1A and 1B). To further evaluate the organization of this differentiation landscape, we applied Slingshot-based pseudotime inference (see Methods).^35^ In contrast to a previously described single maturation trajectory,^17^ this analysis identified two trajectories emerging from the least differentiated cells and extending into more differentiated regions of cDC1 population (Figure 1C). This provided a framework to examine whether steady-state cDC1s contain separable differentiated gene programs.

**Figure 1.**
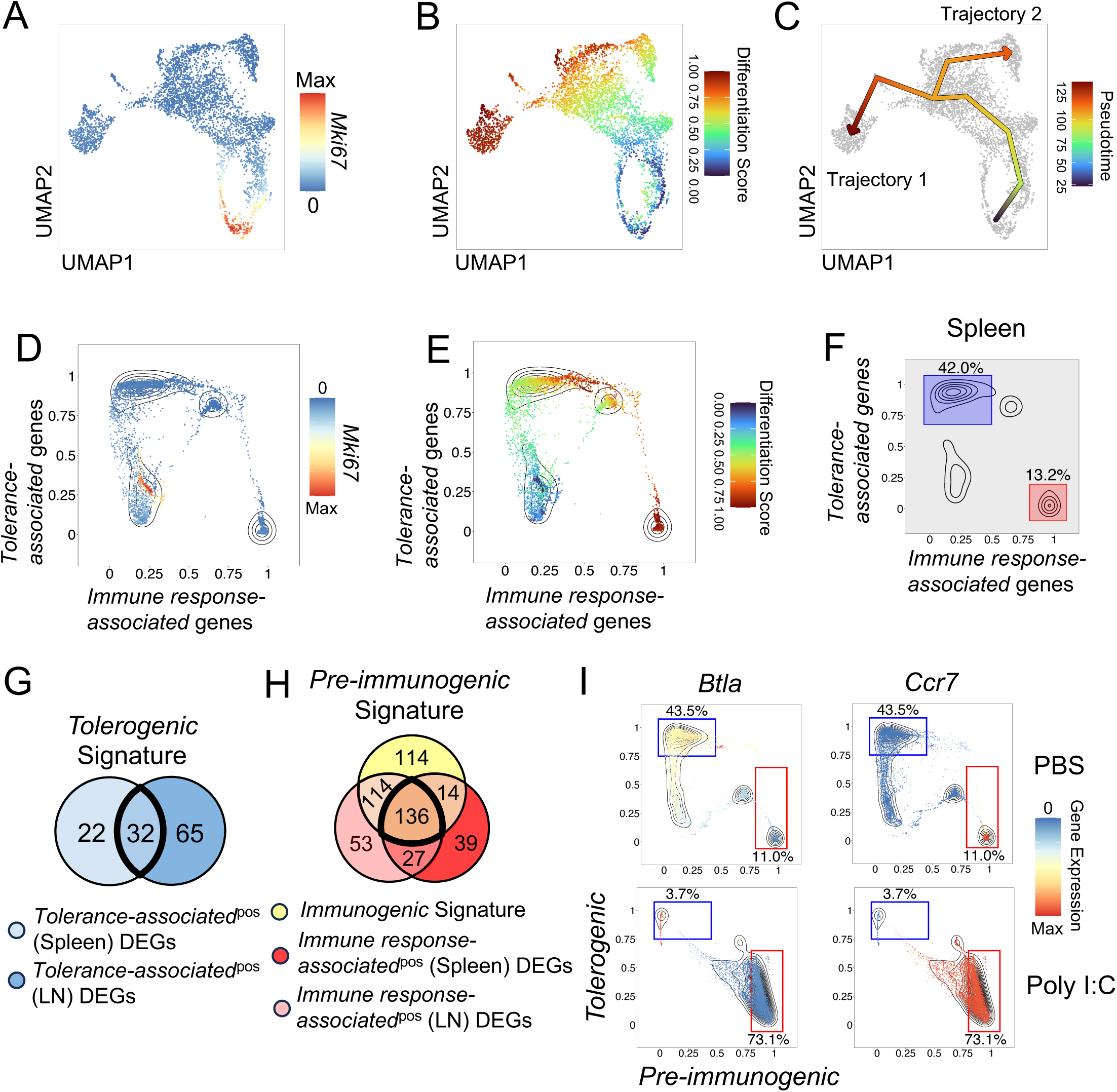
Steady-state cDC1s contain distinct *Tolerogenic* and *Pre-immunogenic* transcriptomic programs. **(A-F)** cDC1s were sorted from spleens or peripheral lymph nodes of wild-type C57BL/6J mice, and scRNA-seq was performed (see Figure S1A for experimental design and gating strategy). **(A)** Uniform manifold approximation and projection (UMAP) embedding of splenic cDC1s. Color overlay indicates the expression of *Mki67*. **(B)** UMAP embedding of splenic cDC1s. Color overlay indicates the differentiation score, calculated as 1-CytoTRACE^2^ (see Methods). **(C)** UMAP embedding of splenic cDC1s overlaid with two main pseudotime trajectories (see Methods). Color of the arrows indicates the position along the pseudotemporal trajectory. **(D-F)** Plot showing distributions of Seqtometry scores for the *Tolerance-associated* gene (*Ahr, Btla, Entpd1, Itgae, Tgfb1*) and *Immune response-associated* gene (*Cd40, Cxcl16, Il12b, Il15, Irf1*) signatures among all splenic cDC1s. **(D)** Color overlay indicates the expression of *Mki67* (as in A). **(E)** Color overlay indicates the differentiation score (as in B). **(F)** Regions indicate *Tolerance-associated*^pos^ (blue) and *Immune response-associated*^pos^ (red) populations used for subsequent analysis. Numbers next to regions indicate corresponding percentages. **(G)** Venn diagrams showing derivation of the *Tolerogenic* signature. Colored regions represent the DEGs (see Methods) in *Tolerance–associated*^pos^ cells from spleens and LNs (gated as in Figure 1F and Figure S1E). Numbers indicate genes per section; bold indicates overlapping genes included in the *Tolerogenic* signature. **(H)** Venn diagram showing derivation of the *Pre-immunogenic* signature. Colored regions represent the *Immunogenic* signature and DEGs (see Methods) in *Immune response–associated*^pos^ cells from spleens and LNs (gated as in Figure 1F and Figure S1E). Numbers indicate genes per section; bold indicates the overlapping genes included in the *Inflammatory* and *Pre-immunogenic* signatures as indicated. **(I)** cDC1s were sorted (see Figure S1A for gating strategy) from spleens of wild-type C57BL/6J mice 12 hours after intraperitoneal injection of PBS or poly I:C followed by performing scRNA-seq. Plots showing distributions of Seqtometry scores for indicated signatures among all splenic cDC1s from each indicated treatment overlaid with *Btla* or *Ccr7* expression, as indicated.

The previously described process of homeostatic maturation involves acquisition of gene programs that partially overlap with immunogenic activation.^2,17,22,36^ To test whether steady-state cDC1s contain discrete immune-response-associated and tolerance-associated transcriptomic features, we selected five genes whose products are well established in cDC-mediated immune responses: *Cd40*, *Cxcl16*, *Il12b*, *Il15*, and *Irf1* (“Immune response-associated genes”).^37–42^ We also selected five genes whose products have established roles in tolerance-associated cDC function, *Ahr*, *Btla*, *Entpd1*, *Itgae*, and *Tgfb1* (“Tolerance-associated genes”).^7,43–47^ We visualized the expression of these genes within individual cells and observed clearly delineated patterns that corresponded to regions of highest gene expression (Figures S1B and S1C, and compare to Figures 1A and B). Moreover, differentiation trajectory 1 that we established above overlapped with cells expressing *Immune response-associated* genes, whereas differentiation trajectory 2 overlapped with cells expressing *Tolerance-associated* genes, suggesting distinctly differentiated states of such cDC1s (Figures 1C, S1B, and S1C).

To further evaluate gene program-level structure rather than individual marker expression alone, we next used Seqtometry to quantify signature enrichment in individual cells. Briefly, predefined or data-derived gene sets (signatures) were scored in each cell, biaxial plots were used to identify cells enriched for specific gene programs, and gated populations were then subjected to differential gene expression (DGE) analysis to define additional comprehensive signatures (Figure S1D). We scored individual cells for enrichment of the *Tolerance-associated* and *Immune response-associated* gene sets. Overlaying *Mki67* expression onto the Seqtometry plot confirmed that proliferating cDC1s were largely excluded from the *Tolerance-associated*^pos^ and *Immune response-associated*^pos^ populations (Figure 1D). In contrast, overlaying the differentiation score further revealed that both *Tolerance-associated* and *Immune response-associated* gene set expression occurred predominantly among more differentiated cDC1s (Figure 1E). Collectively, these results reveal two separate groups of differentiated cells within the cDC1 subset in steady state, each expressing distinct gene sets associated with divergent predicted immune functions.

To define these transcriptomic programs more comprehensively, we used Seqtometry plots to identify *Tolerance-associated*^pos^ and *Immune response-associated*^pos^ single-positive cDC1 populations in spleen and peripheral lymph nodes (Figures 1F and S1E). To derive comprehensive gene signatures from these populations, we performed differential gene expression analysis comparing each gated population against all other cDC1s in the corresponding dataset using the thresholds described in the Methods. We identified 32 genes conserved among *Tolerance-associated*^pos^ populations from spleen and peripheral lymph nodes which we defined as the *Tolerogenic* signature (Figure 1G; genes listed in Table S1).

We next asked whether the genetic program expressed in steady-state *Immune response-associated^pos^* cells was related to genes induced by inflammatory stimulation. Previous studies found that some genes expressed by cDCs under homeostatic conditions overlap with genes induced under proinflammatory conditions, including after administration of strong adjuvants or high molecular weight poly I:C, a TLR3 and MDA5 agonist that mimics viral double-stranded RNA.^2,10,22,48,49^ Therefore, we established an *Immunogenic* signature consisting of genes whose expression was upregulated in the majority of cDC1s after poly I:C treatment (Figure S1F, genes listed in Table S1). Consistent with previously established gene expression patterns,^17,22,36^ we identified 136 genes overlapping between the *Immunogenic* signature and the *Immune response-associated* DEG gene sets from lymph nodes and spleens, which we defined as the *Pre-immunogenic* signature (Figure 1H; genes listed in Table S1). Thus, the *Pre-immunogenic* signature identifies steady-state immune-response-associated genes that overlap with the inflammatory stimulation-induced program. To relate these steady-state signatures to recently described early immature, late immature, early mature, and late mature cDC1 maturation states, ^17^ we overlaid these maturation signatures onto the steady-state PBS cDC1 UMAP (Figure S2A). The UMAP region enriching the *Tolerogenic* signature aligned most closely with the region that enriched the *Late immature* cDC1 signature, whereas the *Pre-immunogenic* signature corresponded more closely to the *Late mature* cDC1 signature in their comparative enrichments (Figure S2A). Thus, the cDC1s with *Tolerogenic* and *Pre-immunogenic* programs show relationships to published maturation-associated states. However, *Tolerogenic* and *Pre-immunogenic* programs occupy distinct regions of the steady-state cDC1 landscape, are associated with separate Slingshot trajectories, and are enriched in more differentiated cDC1s rather than simply reflecting proliferating or immature versus mature cells.

To directly determine the effects of inflammatory stimulation on cDC1s, we applied the *Tolerogenic* and *Pre-immunogenic* signatures to cDC1s from PBS- and poly I:C-treated mice. As discussed above, under steady-state conditions, the *Tolerogenic* and *Pre-immunogenic* signatures identified distinct cDC1 populations. Following poly I:C treatment, the *Tolerogenic*^pos^ population was reduced, whereas the *Pre-immunogenic*^pos^ population increased (Figure 1I). Overlay of *Btla* and *Ccr7* expression showed that *Btla* expression aligned with *Tolerogenic*^pos^ population that was reduced after poly I:C treatment, whereas *Ccr7* expression correlated with *Pre-immunogenic*^pos^ population that increased following poly I:C treatment (Figure 1I). These results link the transcriptomicly defined *Tolerogenic* and *Pre-immunogenic* signatures to gene expression features previously implicated in tolerogenic and mature cDC states, ^7,8,22,50,51^ a relationship further supported by orthogonal BTLA and CCR7 surface staining (Figure S2B). Analogous to our Seqtometry analysis, we observed a loss of BTLA^hi^CCR7^lo^ cDC1s and gain of BTLA^lo^CCR7^hi^ cDC1s 12 hours after exposure to poly I:C (Figure S2C).

Functional enrichment analyses further supported the biological distinction between these programs. Gene ontology (GO) analysis of the *Tolerogenic* signature revealed enrichment for immune-regulatory pathways, including regulation of tolerance induction, negative regulation of T cell-mediated immunity, negative regulation of leukocyte-mediated cytotoxicity, negative regulation of adaptive immune responses, and negative regulation of immune response to tumor cells (Figure S2D). In contrast, GO analysis of the *Pre-immunogenic* signature showed enrichment for antigen processing and presentation through MHC class I/class Ib, T cell chemotaxis and migration, lymphocyte migration into lymphoid organs, T cell-mediated immunity, T cell receptor signaling, and regulation of T helper cell differentiation (Figure S2E). As an independent validation of these functional profiles, we performed Gene Set Enrichment Analysis (GSEA; see Methods)^52,53^ by comparing each individually gated population against all other cells within the corresponding cDC1 dataset, using gene ontology biological process gene sets related to tolerance induction or immune effector response to confirm distinct tolerogenic and immunogenic transcriptomic profiles of the *Tolerogenic*^pos^ and *Pre-immunogenic*^pos^ populations, respectively, in both spleen and peripheral lymph nodes (Figure S2F and S2G). Together, these analyses show that steady-state cDC1s contain distinct differentiated transcriptomic programs with divergent predicted immune functions. One population is enriched for a *Tolerogenic* signature associated with genes known to have tolerogenic functions in cDCs, whereas a separate population is enriched for a *Pre-immunogenic* signature containing immune-response-associated genes that overlap with inflammatory stimulation-induced programs.

### Inflammatory stimulation extends the *Pre-immunogenic* program and induces a conserved *Inflammatory* cDC1 program

Having identified *Tolerogenic* and *Pre-immunogenic* programs in steady-state cDC1s, we next sought to define the transcriptomic program specifically induced by inflammatory stimulation. To do so, we separated the *Immunogenic* signature into two components: genes overlapping with the steady-state *Immune response-associated*^pos^ DEG sets, which defined the *Pre-immunogenic* signature, and the remaining 114 poly I:C-induced genes, which defined the *Inflammatory* signature (Figure 2A). We then combined PBS- and poly I:C-treated cDC1s into a shared UMAP embedding and overlaid the *Inflammatory* signature. The *Inflammatory* signature was strongly induced after poly I:C treatment, indicating that it captures the stimulation-specific component of inflammatory maturation (Figure 2B). Consistent with the expansion of the *Pre-immunogenic*^pos^ population that we observed under inflammatory conditions (Figure 1I), Seqtometry analysis revealed that poly I:C treatment resulted in the approximately five-fold increased *Inflammatory*^pos^ population. In contrast, the *Tolerogenic*^pos^ population was distinctly reduced (Figure 2C). Thus, inflammatory stimulation induces a stimulation-specific Inflammatory program while reducing tolerance-linked cDC1 features.

**Figure 2.**
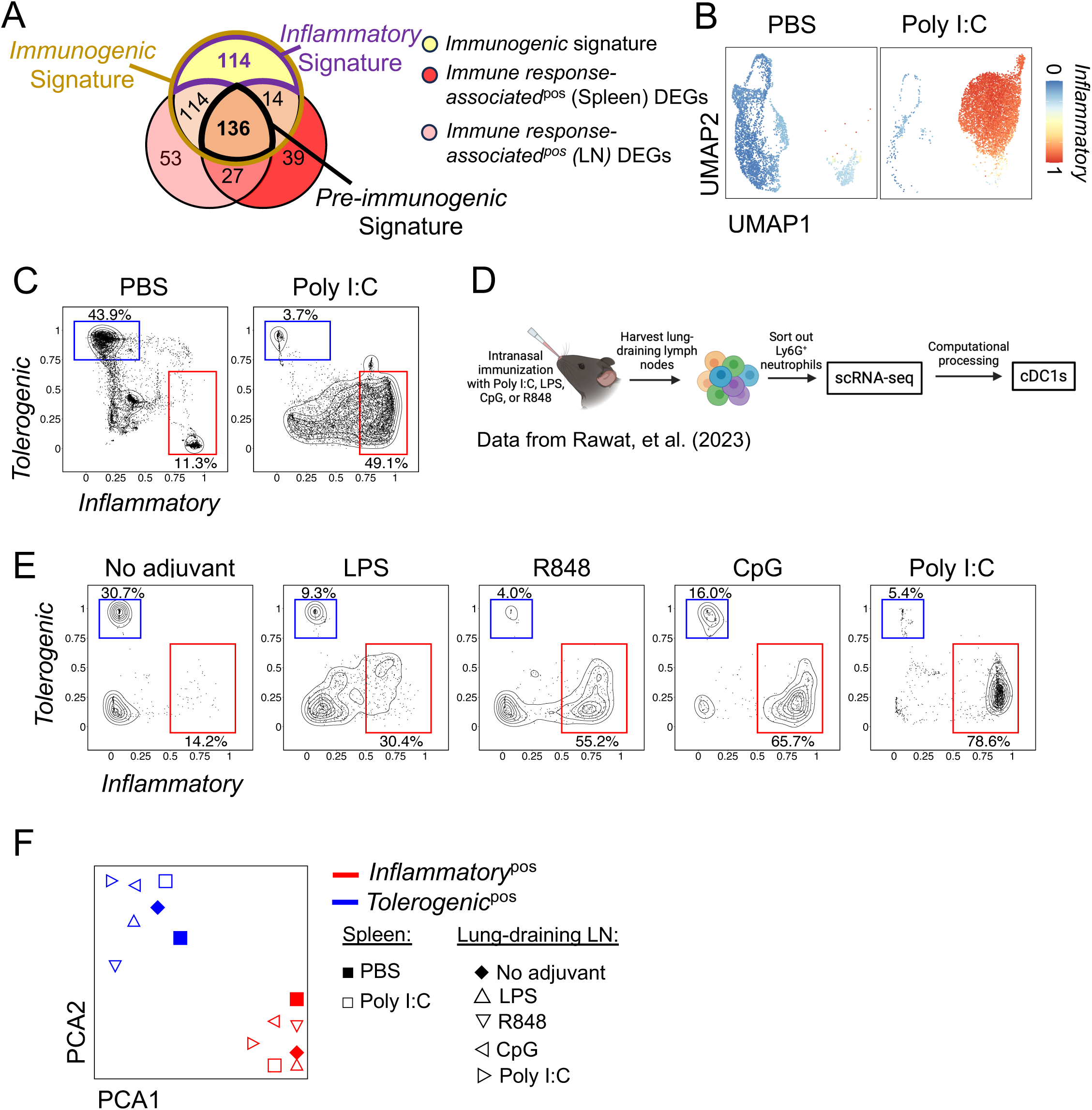
Inflammatory stimulation extends the *Pre-immunogenic* program and induces a conserved *Inflammatory* cDC1 program. **(A)** Venn diagram showing derivation of the *Pre-immunogenic* signature and relation to the *Inflammatory* and *Immunogenic* signatures. Colored regions represent the *Immunogenic* signature and DEGs (see Methods) in *Immune response–associated*^pos^ cells from spleens and LNs (gated as in Figure 1F and Figure S1E). Numbers indicate genes per section; bold indicates separate Immunogenic genes included in the *Inflammatory* and *Pre-immunogenic* signatures as indicated. **(B)** UMAP embedding of splenic cDC1s from mice treated with PBS or poly I:C with color overlay indicating the *Inflammatory* signature. **(C)** Plots showing distributions of scores for *Tolerogenic* and *Inflammatory* signatures among cDC1s from spleens of mice treated with PBS or poly I:C. Regions and corresponding percentages indicate *Tolerogenic*^pos^ (blue) and *Inflammatory*^pos^ (red) populations. **(D)** Schematic of experimental design representing cDC1s from murine lung-draining lymph nodes obtained from Rawat, et al.^59^ **(E)** Plots showing distributions of scores for *Tolerogenic* and *Inflammatory* signatures among murine lung-draining lymph nodes cDC1s (as described in Figure 2D) from the indicated treatment conditions. Regions and corresponding percentages indicate *Tolerogenic*^pos^ (blue) and *Inflammatory*^pos^ (red) populations. **(F)** Principal component analysis (PCA) of *Tolerogenic*^pos^ (blue) and *Inflammatory^pos^* (red) populations from murine splenic and lung-draining cDC1s (gated as in Figures 2C and 2E).

We next determined whether the remaining *Tolerogenic*^pos^ *and Pre-immunogenic*^pos^ cells preserved their respective transcriptomic identities under inflammatory conditions by examining expression of individual genes within the *Tolerogenic* and *Pre-immunogenic* signatures. The expression of individual genes comprising the *Tolerogenic* signature was similar in *Tolerogenic*^pos^ cells present under steady-state conditions and those remaining after poly I:C treatment (Figure 1I and S3A). Similarly, expression of individual *Pre-immunogenic* signature genes was comparable in *Pre-immunogenic*^pos^ cells under steady-state and proinflammatory conditions (Figure 1I and S3B). At the same time, Tolerogenic genes were not induced in the *Pre-immunogenic*^pos^ population under proinflammatory conditions, nor were *Pre-immunogenic* genes induced in the remaining *Tolerogenic*^pos^ population (Figures S3A and S3B). Overall, these analyses further support the separation of *Tolerogenic* and *Pre-immunogenic* gene programs and indicate that inflammatory stimulation reduces the abundance of tolerance-linked cells without converting the remaining populations into a mixed transcriptomic state.

We next compared these gene program-defined populations to previously described cDC maturation signatures. To provide a dataset-internal reference for the least differentiated cDC1s in these overlap analyses, we derived an *Immature* signature by comparing cells with low differentiation scores (1-CytoTRACE2 < 0.2) to all other cDC1s (Figure S3C and Table S1). Notably, the *Immature* signature did not overlap with the *Tolerogenic* signature (Figure S3C). As expected, the *Pre-immunogenic and Inflammatory* signatures partially overlapped with the *Convergent* signature previously established to include genes induced under immunization conditions and homeostatic maturation^22^ as well as some other previously established signatures from CCR7^hi^ migratory cDC populations, consistent with their roles in the initiation of immune responses (Figure S3C).^19,54–58^ Moreover, the *Tolerogenic* signature did not overlap with the multiple other signatures, further underscoring the specificity of our current analysis (Figure S3C). Separately, we also compared our signatures against the recently described maturation stages of cDC1s^17^ (Figure S3D) and LNP-induced maturation^36^ (Figure S3E). We observed a partial overlap between the *Tolerogenic* signature with the *Late immature* signature, consistent with our findings above (Figure S3D). We also found a substantial overlap between the *Pre-immunogenic* signature with *pIC-LNP-induced cDC1 maturation*, *Homeostatic cDC1 maturation*, and *Common signature cDC1 maturation* signatures (Figure S3E). Overall, our framework relates to prior maturation nomenclature while preserving distinctions revealed by gene program-level analysis.

To determine whether the *Inflammatory* program is specific to poly I:C or is more broadly induced by inflammatory stimuli, we applied the *Tolerogenic* and *Inflammatory* signatures to a publicly available scRNA-seq dataset of cDC1s from lung-draining lymph nodes of mice treated intranasally with poly I:C, LPS, R848, or CpG (Figure 2D).^59^ These adjuvants activate distinct innate sensing pathways, including TLR3/MDA5-associated sensing, TLR4 signaling, TLR7/8 signaling, and TLR9 signaling.^60,61^ Across these inflammatory conditions, we detected an increase in *Inflammatory*^pos^ cDC1s accompanied by a reduction in *Tolerogenic*^pos^ cDC1s, although the magnitude of this shift varied by stimulus (Figure 2E). This variability is consistent with the distinct innate sensing pathways engaged by different adjuvants and with the possibility that some stimuli activate cDC1s indirectly through cytokines or signals produced by other cell types. For example, R848-driven cDC1 activation may occur in part through signals derived from directly TLR7/8-responsive cell types, including cDC2s, consistent with prior evidence that direct *cis*-activation of DC subsets shapes their immunogenic capacity.^62^ Similarly, LPS induces TNF-α production by directly responding cDC1s that can indirectly act on neighboring cDC1s that do not themselves sense LPS.^50^

We next performed a transcriptome-level comparison of *Tolerogenic*^pos^ and *Inflammatory*^pos^ populations identified across the adjuvant datasets and our PBS and poly I:C splenic cDC1 dataset. Principal component analysis (PCA) separated *Tolerogenic*^pos^ and *Inflammatory*^pos^ populations into distinct groups (Figure 2F). *Inflammatory*^pos^ populations from distinct inflammatory stimuli clustered together relative to *Tolerogenic*^pos^ populations (Figure 2F). Together, these results demonstrate that inflammatory maturation extends a *Pre-immunogenic* program already present in steady-state cDC1s while inducing an additional *Inflammatory* gene program and reducing tolerance-linked molecular features. At the same time, stimulus-dependent variation suggests that this conserved inflammatory program can be further shaped by the specific innate sensing pathway and cellular context of activation.

### IFNAR signaling governs the core inflammatory cDC1 program

Type I interferon (IFN-I) transduces signals via the interferon-α/β receptor (IFNAR), which regulates complex responses in multiple cell types, including cDCs, across health and disease contexts.^63^ To determine whether the mouse-derived *Inflammatory* program could be induced by IFN-I in human cDCs, we translated the *Inflammatory* signature into human orthologs (see Methods and Table S2) and analyzed a published scRNA-seq dataset of peripheral blood cDCs from healthy donors cultured with IFN-β for different lengths of time (Figure 3A).^64^ Before IFN-β exposure, few human cDCs enriched the *Inflammatory* signature (Figure 3B). However, within 4 hours of IFN-β treatment, cDCs showed robust upregulation of the *Inflammatory* signature that persisted through the remainder of the 36-hour culture period (Figure 3B). These findings indicate that the Inflammatory program identified in mouse cDC1s is conserved in human cDCs and can be induced by IFN-I.

**Figure 3.**
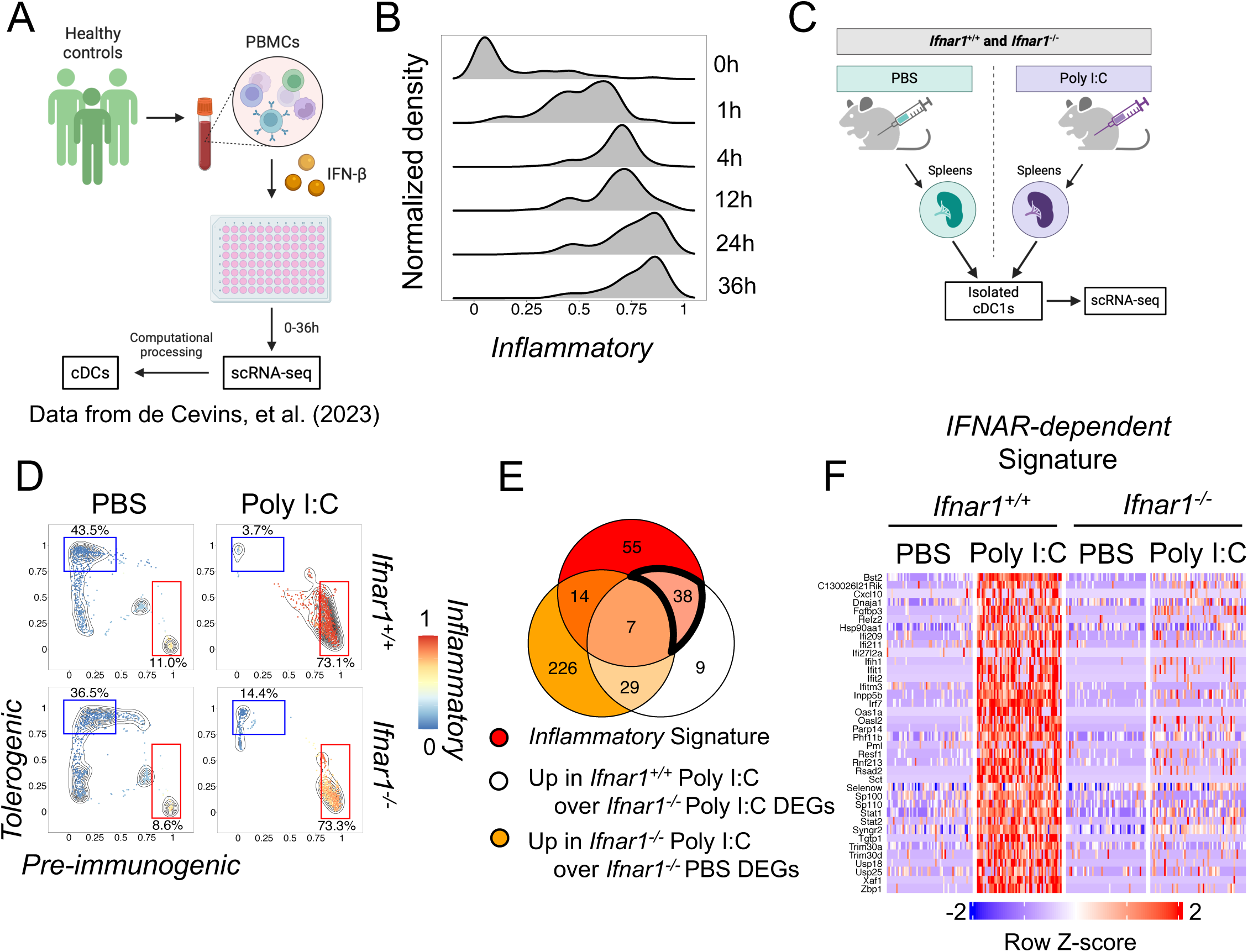
IFNAR signaling governs a conserved core inflammatory cDC program. **(A)** Schematic of experimental design representing data from in vitro–cultured human cDCs obtained from de Cevins, et al.^64^ **(B)** Histogram showing score distribution for the *Inflammatory* signature (translated to human orthologs) among human cDCs from A. **(C)** Schematic of experimental design. cDC1s were sorted from pooled spleens of *Ifnar1*^+/+^ or *Ifnar1*^-/-^ mice 12 hours after intraperitoneal injection of PBS or poly I:C followed by scRNA-seq. **(D)** Plots showing distributions of scores for indicated signatures among splenic cDC1s from C and for each treatment overlaid with the *Inflammatory* signature. Regions and corresponding percentages indicate *Tolerogenic*^pos^ (blue) and *Pre-immunogenic*^pos^ (red) populations. **(E)** Venn diagram showing derivation of *IFNAR-dependent* signature indicated within a bold intersection. Colored regions represent the Inflammatory signature described above and the individual gene sets corresponding to DEGs among all splenic cDC1s from *Ifnar1*^+/+^ or *Ifnar1*^-/-^mice treated with PBS or poly I:C, as indicated and analyzed by scRNA-seq (see Methods). Numbers of genes in specific sections are indicated. **(F)** Heatmap of genes of the *IFNAR-dependent* signature for splenic cDC1s from *Ifnar1*^+/+^ or *Ifnar1*^-/-^ mice treated with PBS or poly I:C (gated as in Figure 3D).

IFN-I-dependent responses after experimental immunization, including poly I:C treatment, are well established.^10,11,65–67^ To directly test the contribution of IFNAR signaling to the *Inflammatory* program *in vivo*, we used an established *Ifnar1*-deficient mouse model.^68^ We performed scRNA-seq on cDC1s sorted from spleens of *Ifnar1*^+/+^ and *Ifnar1*^-/-^ mice treated with PBS or poly I:C (Figure 3C and Methods). Under steady-state conditions, *Ifnar1*^-/-^ cDC1s contained *Tolerogenic*^pos^ and *Pre-immunogenic*^pos^ populations in proportions similar to *Ifnar1*^+/+^ cDC1s (Figure 3D). This suggests that the *Tolerogenic* and *Pre-immunogenic* programs present under homeostatic conditions do not require IFNAR signaling. In contrast, induction of the *Inflammatory* signature after poly I:C treatment was altered in *Ifnar1*^-/-^ cDC1s (Figure 3D), indicating that IFNAR signaling is required for full induction of the *Inflammatory* program.

To identify the IFNAR-dependent component of the *Inflammatory* program, we performed differential gene expression analysis using two comparisons. First, we identified genes increased in cDC1s from poly I:C-treated *Ifnar1*^+/+^ mice compared with cDC1s from poly I:C-treated *Ifnar1*^-/-^mice. Second, we identified genes induced in cDC1s from poly I:C-treated *Ifnar1*^-/-^ mice compared with PBS-treated *Ifnar1*^-/-^ cDC1s. Genes increased after poly I:C treatment in *Ifnar1*^+/+^ cDC1s but not induced in *Ifnar1*^-/-^ cDC1s were defined as *IFNAR-dependent*. This analysis identified 47 IFNAR-dependent genes, 38 of which overlapped with the *Inflammatory* signature, defining an *IFNAR-dependent* signature (Figures 3E and 3F; genes listed in Table S1). GO analysis confirmed that this *IFNAR-dependent* signature was enriched for type I interferon signaling and immune response regulation (Figure S4A).

We next returned to the human IFN-β time-course dataset to determine whether this refined *IFNAR-dependent* signature was also induced in human cDCs. We translated both the *IFNAR-dependent* and *Inflammatory* signatures into human orthologs and scored peripheral blood cDCs cultured with IFN-β.^64^ As expected, few cells enriched either signature before IFN-β treatment. Within 4 hours of IFN-β exposure, human cDCs showed clear upregulation of both the *IFNAR-dependent* and *Inflammatory* signatures, and this enrichment persisted throughout the culture period (Figure S4B). Together, these findings show that IFN-I is sufficient to induce the *Inflammatory* program in human cDCs and is required for full induction of this program in mouse cDC1s after poly I:C treatment. By resolving the *IFNAR-dependent* component of the broader *Inflammatory* signature, these analyses identify a conserved IFN-I-responsive core of inflammatory cDC maturation that is shared across species and induced in response to inflammatory stimulation.

### The inflammatory cDC program is enriched across human cDC subsets in SLE

Having defined an IFNAR-dependent inflammatory program in experimentally stimulated mouse cDC1s and validated its inducibility in human cDCs, we next asked whether this program could identify inflammatory cDC populations in human disease. Therefore, we extended our analysis to human disease, beginning with systemic lupus erythematosus (SLE), a complex autoimmune disease involving multiple immune cell populations and characterized by increased IFN-I signaling, altered lymphocyte activation, and impaired apoptotic cell clearance.^69,70^ A recent study profiled more than 1.2 million peripheral blood mononuclear cells (PBMCs) from SLE patients and healthy controls and identified a myeloid-specific IFN-I-stimulated gene signature (*Myeloid ISG*) in innate immune cells that was associated with disease activity.^71^ Given the central role of cDCs in bridging innate and adaptive immunity, we asked whether the *Inflammatory* and *IFNAR-dependent* signatures identified above enriched a corresponding inflammatory cDC population in SLE patients. Using the authors’ cDC annotations, we scored peripheral blood cDCs from individual patients and healthy controls with human ortholog equivalents of the mouse-derived signatures established above (Figure 4A). *Inflammatory*^pos^*IFNAR-dependent*^pos^ cDCs were markedly increased in SLE patients compared with healthy controls (Figure 4B). As expected, these *Inflammatory*^pos^*IFNAR-dependent*^pos^ cDCs also expressed the *Myeloid ISG* signature,^71^ whereas cDCs enriched for all three signatures were largely absent from healthy controls (Figures 4B and 4C). The frequency of these inflammatory cDCs also corresponded to disease status: patients with active flare had, on average, approximately 70% *Inflammatory*^pos^*IFNAR-dependent*^pos^ cDCs among total cDCs, whereas this population constituted approximately 45% of cDCs in patients with managed disease (Figure 4B). Inflammatory cDCs also enriched a *Glycolysis* signature (Figure 4B and 4C), consistent with metabolic remodeling associated with activated human immune cells.^26^ Thus, the *IFNAR-dependent* inflammatory program identifies a cDC population that is expanded in SLE and tracks with disease activity.

**Figure 4.**
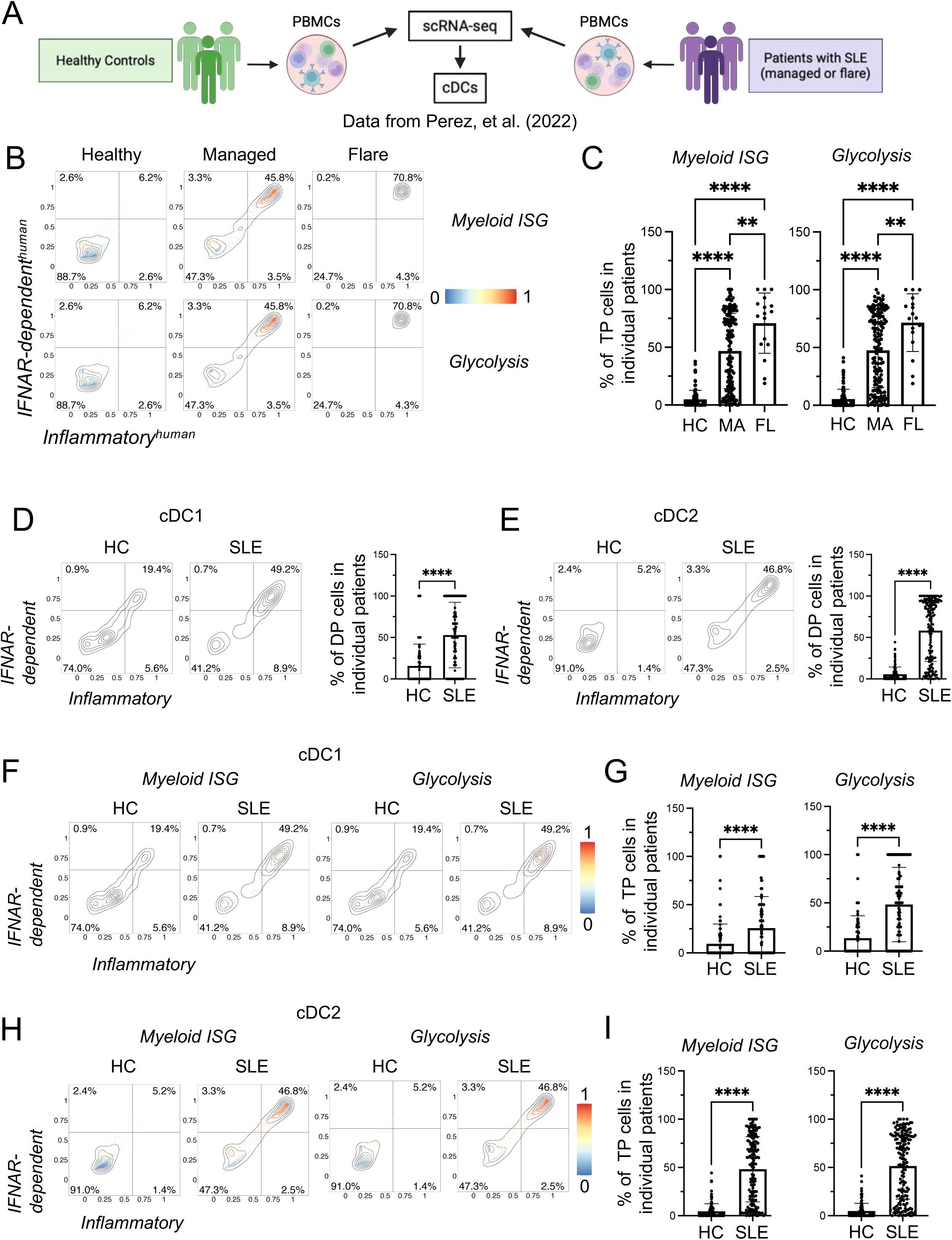
The *IFNAR-dependent* inflammatory cDC program is enriched across human cDC subsets in SLE. **(A)** Schematic of experimental design representing data from peripheral blood mononuclear cells (PBMCs) from systemic lupus erythematosus (SLE) patients and healthy controls obtained from Perez, et al.^71^ **(B)** Plots showing distributions of scores for *Inflammatory* and *IFNAR-dependent* signatures (both signatures translated to human orthologs) among all circulating cDCs from the specified groups overlaid with the indicated signatures. Percentages are indicated for corresponding quadrants. **(C)** Bar graphs showing mean ± SD of triple-positive (TP; *Inflammatory*^pos^, *IFNAR-dependent*^pos^, and positive for the indicated signature) cDCs among individual SLE patients. HC: healthy controls (n=99); MA: managed disease (n=146); FL: flare (n=17). ** P < 0.01, **** P < 0.0001 determined by Dunnett’s T3 multiple comparisons test. **(D-I)** Plots (D, E, F, H) show distributions of scores for indicated signatures overlaid among cDC1s or cDC2s. Bar graphs (D, G for cDC1s; E, I for cDC2s) show mean ± SD of double-positive (DP; *Inflammatory*^pos^, *IFNAR-dependent*^pos^) or triple-positive (TP; *Inflammatory*^pos^, *IFNAR-dependent*^pos^, and also positive for the indicated signature) cDCs. HC: healthy controls; SLE: systemic lupus erythematosus. **** P < 0.0001 determined by Welch’s t-test.

Given the proposed roles of cDC1 and cDC2 subsets in promoting inflammatory responses,^6,9,72–74^ we next stratified the *Inflammatory*^pos^*IFNAR-dependent*^pos^ population among individual cDC subsets using cDC1- and cDC2-specific signatures (Table 2). In most SLE patients, *Inflammatory*^pos^*IFNAR-dependent*^pos^ cells were present at similar frequencies among cDC1s and cDC2s, constituting approximately 46–49% of each subset (Figure 4D and 4E). A subset of healthy controls showed increased frequencies of *Inflammatory*^pos^*IFNAR-dependent*^pos^ cDC1s (Figure 4D), potentially reflecting non-specific inflammatory variation affecting cDC1s in select individuals. In contrast, *Inflammatory*^pos^*IFNAR-dependent*^pos^ cells were negligible among cDC2s from healthy controls (Figure 4E). Consistent with a shared inflammatory state across cDC subsets in SLE, *Inflammatory*^pos^*IFNAR-dependent*^pos^ cDC1s and cDC2s both enriched the *Myeloid ISG* and *Glycolysis* signatures (Figure 4F–I). Therefore, the *IFNAR-dependent* inflammatory program is not restricted to mouse cDC1s or experimentally stimulated human cDCs *in vitro* but is enriched in human cDCs during systemic autoimmunity. In SLE, this program is associated with disease activity, IFN-I-stimulated gene expression, and metabolic features of immune activation. These findings indicate that the *IFNAR-dependent* inflammatory program identified experimentally is also detectable in human disease and can be used to resolve among all cDCs the inflammatory populations associated with disease activity. Together, these analyses underscore a conservation of the *IFNAR-dependent* inflammatory program in both cDC1 and cDC2 subsets during systemic autoimmunity.

### Conservation of the inflammatory molecular program in cDCs across multiple human disease states

IFN-I-dependent mechanisms are involved in various diseases.^64,69,75,76^ Given the relative ease of collecting total cDCs from PBMCs, multiple scRNA-seq datasets containing such cDCs from human patients are available. Therefore, we extended our analysis to include peripheral blood scRNA-seq datasets from other disease processes, including viral infection (COVID-19^76^), autoinflammation (STING-associated vasculopathy with onset in infancy, SAVI^64^), along with corresponding healthy controls. To extend this analysis beyond circulating cDCs, we also examined cDCs from solid-tissue datasets, including neoplastic tissues and matched healthy controls. Specifically, we included scRNA-seq datasets from primary pancreatic ductal adenocarcinoma (PDAC) and control pancreases.^77^ We also included datasets from primary breast carcinomas from treatment-naive patients (i.e., estrogen receptor (ER^+^) and progesterone receptor (PR^+^) positive, human epidermal growth factor receptor 2 amplified (Her2^+^), and triple-negative (TNBC), breast cancers and matched normal breast tissue controls.^78^ As a negative control, we used a pulmonary fibrosis (PF) dataset in which cDC involvement was not reported as a central feature of the disease process.^79^ In all samples, we identified cDCs using the original authors’ annotations and examined them by using the human orthologs of the key *Inflammatory* and *IFNAR-dependent* signatures we established above as specific for the mature inflammatory population (Figure 5A-F). Because scoring with gene signatures is relative and only valid within individually analyzed groups that include cells from patients and the specific disease dataset-matched healthy controls, we represented independent scoring results for patients and matched healthy controls within each disease-specific group on biaxial plots. Strikingly, for SLE, PDAC, breast cancer, SAVI, and COVID-19, we observed statistically significant increased enrichments with *Inflammatory* and *IFNAR-dependent* signatures among cDCs from patients with the indicated disease compared to their matched healthy controls (Figure 5A-E). In contrast, no statistically significant enrichments were found among patients with PF and their matched healthy controls (Figure 5F). These results demonstrate that the inflammatory cDC program is not restricted to a single disease. Instead, the *Inflammatory and IFNAR-dependent* signatures identify corresponding cDC populations across multiple human inflammatory disease contexts, including circulating and tissue-resident cDCs.

**Figure 5.**
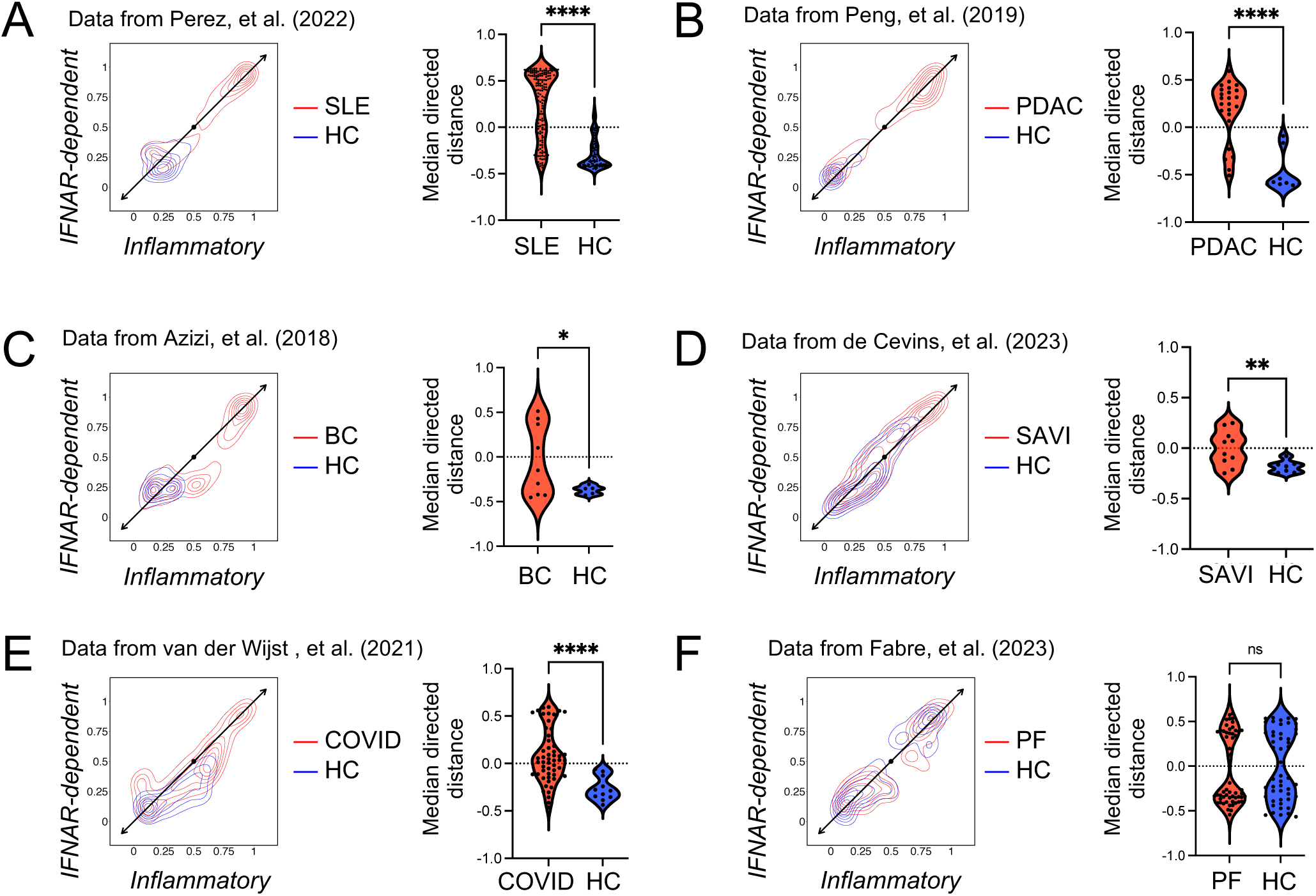
The inflammatory cDC program is conserved across multiple human disease states. **(A–F)** Data were obtained from van der Wijst, et al.,^76^ de Cevins, et al.,^64^ Perez, et al.,^71^ Peng, et al.,^77^ Azizi, et al.,^78^ and Fabre, et al.^79^ HC: corresponding data set-matched healthy controls, SLE: systemic lupus erythematosus^71^ (A), PDAC: pancreatic ductal adenocarcinoma^77^ (B), BC: breast cancer^78^ (C), SAVI: STING-associated vasculopathy with onset in infancy^64^ (D), COVID: coronavirus disease 2019^76^ (E), PF: Pulmonary fibrosis^79^ (F). Plots show distributions of *Inflammatory* and *IFNAR-dependent* signature scores (both signatures translated to human orthologs) from all individuals within indicated groups. The indicated diagonal axis corresponds to positive and negative joint enrichment of *Inflammatory* and *IFNAR-dependent* signatures relative to the central point. Violin plots show distribution of median directed distances (from each plot’s central point) along the indicated diagonal axis for individual subjects from indicated groups. (SLE: n=99 HC, 163 patients; PDAC: n=9 HC, 24 patients; BC: n=5 HC, 9 patients; SAVI: n=7 HC, 10 patients; COVID: n=11 HC, 53 patients, PF: n=50 HC, n=56 patients). n.s.: not significant, * P < 0.05, ** P < 0.01, **** P < 0.0001 determined by Welch’s t-test.

### Disease-specific gene signatures reveal contextual diversification of inflammatory cDCs

We next asked whether inflammatory cDCs from distinct disease contexts share only a common core program or also acquire disease-specific transcriptomic features. To test this, we compared *Inflammatory*^pos^*IFNAR-dependent*^pos^ cDCs with corresponding *Inflammatory*^neg^*IFNAR-dependent*^neg^ cDCs within each disease dataset and performed individual differential gene expression analyses to define disease-specific inflammatory cDC signatures (Figure 6A, Table S2, and Methods). We then used these signatures to determine whether inflammatory cDCs from each disease could be distinguished from inflammatory cDCs in other disease contexts. Remarkably, each disease-specific signature preferentially identified inflammatory cDCs from the corresponding disease dataset but not the others (Figure 6B-F). This is consistent with the idea that although inflammatory cDCs share a conserved core program, they also acquire context-specific transcriptomic features that distinguish the disease environment in which they arise.

**Figure 6.**
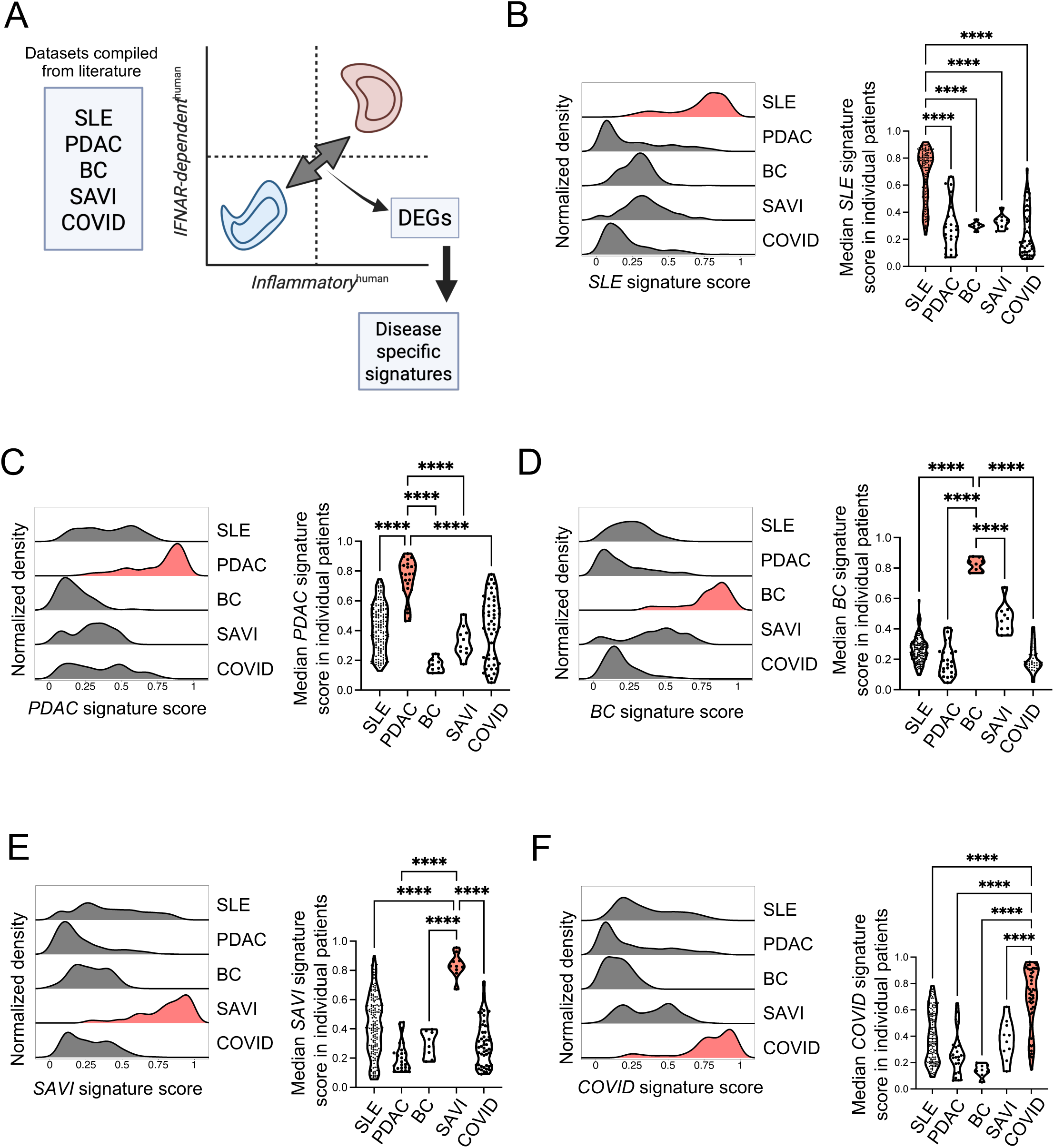
Disease-specific gene signatures reveal contextual diversification of inflammatory cDCs. **(A)** Schematic diagram showing derivation of disease-specific signatures. Data were obtained from van der Wijst, et al.,^76^ de Cevins, et al.,^64^ Perez, et al.,^71^ Peng, et al.,^77^ and Azizi, et al.,^78^ HC: corresponding data set-matched healthy controls, SLE: systemic lupus erythematosus,^71^ PDAC: pancreatic ductal adenocarcinoma,^77^ BC: breast cancer,^78^ SAVI: STING-associated vasculopathy with onset in infancy,^64^ COVID: coronavirus disease 2019.^76^ **(B–F)** Histograms showing score distributions of the indicated disease-specific signatures among cDC populations that enriched *Inflammatory* and *IFNAR-dependent* signatures within each patient group as shown in Figure 5. Violin plots show analogous distributions of median scores from individual patients. Red highlights populations matching the disease-specific signature. **** P < 0.0001 determined by Dunnett’s T3 multiple comparisons test. HC: healthy control, SLE: systemic lupus erythematosus, PDAC: pancreatic ductal adenocarcinoma, BC: breast cancer, SAVI: STING-associated vasculopathy with onset in infancy, COVID: coronavirus disease 2019

To define the biological processes captured by these disease-specific inflammatory cDC signatures, we performed GO enrichment analysis (Figure S5A-E). The SLE-specific signature was enriched for processes related to unfolded protein response, oxidative stress, vascular permeability, TGF-β production, monocyte differentiation, cell–cell signaling, and regulation of T cell-mediated cytotoxicity, consistent with systemic inflammatory, vascular, and lymphocyte-associated pathology in SLE (Figure S5A). The PDAC-specific signature was enriched for epithelial, mesenchymal, endothelial, morphogenetic, and tissue-remodeling processes, consistent with the stromal and desmoplastic features of PDAC (Figure S5B). The breast cancer-specific signature showed enrichment for cell–cell adhesion, cadherin-mediated interactions, TGF-β/BMP/Wnt-associated signaling, angiogenesis, senescence, leukocyte proliferation, and immune receptor signaling, suggesting that inflammatory cDCs in breast tumors acquire transcriptomic features linked to tumor-stromal remodeling and immune–tissue interactions (Figure S5C). The SAVI-specific signature was enriched for IL-18 signaling, NK cell activation, type I IFN signaling, IL-6 signaling, T cell activation, B cell activation, NF-κB transcriptomic activity, and antiviral defense pathways, consistent with STING-driven autoinflammation and type I interferonopathy (Figure S5D). Finally, the COVID-specific signature was enriched for amino acid metabolism, cholesterol metabolism, cysteine/glutathione and redox pathways, complement, B cell receptor signaling, unfolded protein response, and oxygen-response pathways, consistent with viral infection-associated metabolic stress, oxidative stress, complement activation, and humoral immune features (Figure S5E). Together, these analyses show that inflammatory cDCs share a conserved *IFNAR-dependent* core program but also undergo disease-specific diversification. Thus, inflammatory cDCs retain a conserved IFNAR-dependent maturation core while acquiring disease-specific transcriptomic features that reflect the inflammatory, metabolic, and tissue environments in which they arise.

## DISCUSSION

Our results reveal among heterogenous human and mouse cDCs a unified inflammatory maturation program dependent on IFN-I-mediated signaling and conserved across many conditions and disease states. The initiation of immune responses has been proposed to depend on acquisition of immunogenic properties by fully differentiated cDCs.^1,21,22^ More recent studies have further distinguished homeostatic and immunogenic modes of cDC maturation, emphasizing that these processes share broad transcriptomic features while differing in restricted molecular modules associated with inflammatory or tolerogenic function.^2,17,21,22,36^ Our studies clarify this paradigm by showing that inflammatory conditions specifically extend an immune response-associated program that is present under homeostatic conditions in a defined group of cDC1s. We refer to this steady-state program as *Pre-immunogenic* because it contains immune-response-associated genes that overlap with the inflammatory stimulation-induced program, while remaining distinct from the fully induced *Inflammatory* program. In contrast, a unique transcriptomic profile including multiple genes whose products have been established in immunological tolerance^7,43–47^ is separated within distinct cDC1s, consistent with the well-established functions of tolerogenic cDC1s in spleen and systemic peripheral lymph nodes.^6,8^

Importantly, our results do not simply assign these populations to an immature-versus-mature axis. The *Tolerogenic* program showed relationships to previously described late immature cDC1 states, whereas the *Pre-immunogenic* program aligned more closely with late mature or homeostatic/immunogenic maturation-associated states.^17,36^ However, the *Tolerogenic* signature did not overlap with our dataset-derived *Immature* signature and was instead associated with a distinct trajectory, differentiation-associated transcriptomic features, and immune-regulatory functional annotations. Thus, the *Tolerogenic* and *Pre-immunogenic* programs show partial relationships to published maturation nomenclature but are defined here by broader tolerance-associated and immune-response-associated gene programs rather than by maturation status alone. This distinction is important because collapsing these programs into previously defined immature, homeostatic mature, or immunogenic mature categories would obscure the separation of tolerance-linked and immune-response-associated molecular features revealed by our analysis.

Functional enrichment analyses further support this framework. The *Tolerogenic* signature was enriched for immune-regulatory processes, including regulation of tolerance induction, negative regulation of T cell-mediated immunity, negative regulation of leukocyte-mediated cytotoxicity, and negative regulation of adaptive immune responses. In contrast, the *Pre-immunogenic* signature was enriched for antigen processing and presentation, T cell chemotaxis and migration, lymphocyte migration into lymphoid organs, T cell-mediated immunity, T cell receptor signaling, and regulation of T helper cell differentiation. These findings suggest that steady-state cDC1s contain distinct differentiated programs with divergent predicted immune functions. One population is enriched for molecular features associated with immunological tolerance, whereas another expresses an immune-response-associated program that can be extended under inflammatory conditions.

Such *Tolerogenic* cDC1s are reduced at the onset of inflammatory conditions in agreement with an emerging model that postulates a specific ablation of tolerogenic cDCs in response to proinflammatory signals while promoting activation of other cDCs with immunogenic functions.^8,9,50^ A specific reduction of *Tolerogenic* cDC1s may also be a manifestation of their death or specific conversion into immunogenic cDCs.^6,8,9,50,80^ Such an immunogenic conversion, if true, would likely involve cDCs with transitional molecular features, which are not identified in our studies, although they may occur transiently at earlier timepoints. Similarly, future studies might uncover induced tolerogenic features among some inflammatory cDCs akin to homeostatic mechanisms discussed above.^17–20,81,82^ Overall, our current results support a separation of molecular tolerogenic and immunogenic programs within individual cDCs, while leaving open the possibility that additional kinetic analyses may reveal transient transitional states between these programs.

These results are consistent with some previously proposed paradigms of homeostatic maturation and diverse T cell responses that may be induced under steady state conditions.^2,17,21,22,36,80,82–86^ Moreover, the *Pre-immunogenic* and mature cDC1 populations we identified may relate to human CCR7^+^ cDC1s recently shown to control the development and maintenance of tertiary lymphoid structures in human cancer.^87^ Crucially, however, our results provide specific insight into the precisely defined inflammatory maturation process as it builds on a molecular framework pre-existing under homeostatic conditions, excluding most tolerogenic features characterizing cDCs in the steady state. Our use of 12-hour poly I:C treatment should therefore be interpreted as a defined TLR3/MDA5-associated inflammatory condition rather than capturing the peak or complete duration of inflammatory cDC activation. DC activation is highly kinetic, and inflammatory and homeostatic maturation programs can share broad transcriptomic features and vary in magnitude depending on stimulus, formulation, tissue, and time after activation.^17,22,36^ Nevertheless, reduced strength of inflammatory modules at later time points or partial overlap between inflammatory and homeostatic maturation signatures as described by others ^36^ does not by itself demonstrate reversion to a *bona fide* homeostatic mature state. In our data, 12-hour poly I:C-treated cDC1s retained clear induction of the *Inflammatory* and *IFNAR-dependent* signatures that are conserved across species and diseases, supporting its use as an inflammatory comparator in this framework.

Our studies further clarify that interferon-α/β receptor (IFNAR) signaling orchestrates the core molecular program within the inflammatory population and that this program is absent from the persisting tolerogenic transcriptomic program. The role of IFN-I in cDC activation and immunogenic maturation is well established.^10–14,22,36,88^ Prior transcriptomic and functional studies have also identified IFN-I-associated features in thymic cDC1s, including ISG expression^22^ and IFN-dependent activation and maturation of thymic cDC1.^88^ However, because the thymus is a specialized immune organ dedicated to central tolerance and contains distinct developmental, antigenic, and cytokine environments, we considered these thymic IFN-I-associated programs conceptually relevant but not appropriate as direct comparators for the splenic and disease-associated inflammatory cDC programs analyzed here. Thus, the novelty of our work is not the identification of IFN-I as a DC-activating signal *per se*, but rather the placement of IFNAR-dependent genes at the core of a conserved inflammatory maturation framework that is separable from steady-state *Tolerogenic* and *Pre-immunogenic* programs. We further show that IFN-I-dependent signaling is dispensable for the homeostatic transcriptomic identities of splenic cDC1s in the steady state. This differs from some reports on the role of IFN-I for cDCs in the gut,^16^ possibly underscoring specificities of anatomical barrier tissues and local cytokine environments.

The inflammatory program was induced across multiple inflammatory conditions, including poly I:C, LPS, R848, and CpG exposure. However, the magnitude of this response varied by stimulus, consistent with the distinct innate sensing pathways engaged by different adjuvants and with the possibility that some stimuli activate cDC1s indirectly through cytokines or signals produced by other cell types. For example, R848-driven cDC1 activation may occur in part through signals derived from directly TLR7/8-responsive cell types, including cDC2s, whereas LPS can induce inflammatory mediators from directly responsive cDCs that act on neighboring cDC1s.^50,62,89^ Thus, our findings identify a conserved inflammatory program induced across inflammatory contexts, but do not imply that all stimuli generate identical cDC states or identical immunogenic capacities. Rather, the conserved inflammatory core can be further shaped by stimulus-specific sensing pathways, direct versus indirect activation, tissue context, and cellular microenvironment, as further revealed by the disease-specific signatures that we established.

The IFN-I-dependent inflammatory gene expression profile is induced in both cDC1s and cDC2s, consistent with the predominant inflammatory functions of cDC2s.^6,9,72–74^ Therefore, the results underscore the conservation of the molecular inflammatory program, further indicating an important complementarity of such responses across different cDC subsets. Remarkably, we revealed that the *Inflammatory* signature is shared in circulating and tissue-resident cDCs across multiple human disease states, including infection, autoimmunity, autoinflammation, and cancer, further indicating its broad conservation. By demonstrating that IFN-I-dependent signaling is a key molecular feature of the conserved maturation program of specific cDCs in these patients, our results help to clarify the known roles of IFN-I and its signaling in multiple diseases.^64,69,71,75,76,90^

The enrichment of the *IFNAR-dependent* inflammatory program in SLE further supports its disease relevance. *Inflammatory* cDCs were increased in SLE patients compared with healthy controls, enriched a myeloid IFN-stimulated gene signature, and were more frequent in patients with active flare than in patients with managed disease.^71^ These cells also enriched a *Glycolysis* signature, consistent with metabolic remodeling associated with activated human immune cells.^26^ However, we do not interpret glycolytic enrichment as a novel feature of inflammatory DC biology. Instead, it provides a functional correlate of the transcriptomicly defined inflammatory state. Importantly, the experimentally defined mouse cDC1 inflammatory program identifies a corresponding IFNAR-dependent cDC population in human systemic autoimmunity and this population is detectable across both major circulating cDC subsets.

Crucially, we now uncovered disease-specific gene expression profiles present in corresponding inflammatory populations. These findings support a model in which inflammatory cDCs contain two layers of transcriptomic identity: a conserved *IFNAR-dependent* inflammatory maturation core and a disease- or tissue-specific transcriptomic layer imposed by the local pathological environment. The biological processes enriched in disease-specific signatures suggest that inflammatory cDCs may encode features of the disease context in which they arise, including systemic inflammatory and vascular features in SLE, stromal and tissue-remodeling programs in pancreatic and breast cancer, STING-associated autoinflammatory pathways in SAVI, and metabolic, redox, complement, and antiviral features in COVID-19. Although further research is necessary, such unique gene signatures may be informative regarding the specific biological processes related to individual diseases and anatomical microenvironments, possibly becoming a foundation for future precision medicine bioindicators.

Our findings also raise practical possibilities for future translational studies. The conserved *Inflammatory* and *IFNAR-dependent* signatures may help identify inflammatory cDC populations in patient datasets and may serve as starting points for a development of biomarkers and future therapeutically relevant targets in diseases where cDC activation or IFN-I responsiveness correlates with disease activity. In addition, disease-specific inflammatory cDC signatures may help distinguish shared inflammatory activation from context-dependent cDC adaptation. Future work will be needed to determine whether these transcriptomic states can be isolated using surface or intracellular markers. BTLA and CCR7 provide useful protein-level anchors for comparing tolerance-associated and canonical maturation-associated features, whereas additional candidates, including intracellular molecules such as A20/Tnfaip3, may help enrich *Pre-immunogenic* or *Inflammatory* cDC states for functional testing. Such approaches may eventually enable direct assessment of whether these populations differ in antigen presentation, T cell priming, tolerance induction, or therapeutic responsiveness.

Overall, our studies clarify the key roles of IFN-I in the inflammatory maturation program of specific cDCs and reveal closely choreographed dynamic processes occurring under inflammatory conditions to determine the outcomes of cDC maturation. In conclusion, our findings show that IFN-I signaling orchestrates a conserved inflammatory maturation program in cDCs, spanning mouse and human systems, multiple cDC subsets, and diverse disease states. At the same time, inflammatory cDCs acquire disease-specific transcriptomic features that reflect the inflammatory, metabolic, and tissue environments in which they arise. The transcriptomic characteristics identified here therefore provide a robust framework for future studies aimed at defining cDC function, developing disease bioindicators, and evaluating diagnostic or therapeutic applications.

## ACKNOWLEDGMENTS

The authors would like to thank Saint Louis University Flow Cytometry Core for expert help with flow cytometry sorting.

## AUTHOR CONTRIBUTIONS

Conceptualization, development of the methodology, and formal analysis, J.B., R.K. and D.H.; Animal experiments and analysis, J.B. and D.H.; Software and bioinformatics analysis, R.K. and D.H.; Additional analysis and visualization, J.B., R.K. and D.H.; Writing - original draft, J.B. and D.H.; Writing - review & editing, J.B., R.K. and D.H.; Supervision, D.H.; Funding Acquisition, D.H.

## MATERIAL AND METHODS

## REAGENT AND RESOURCE TABLE

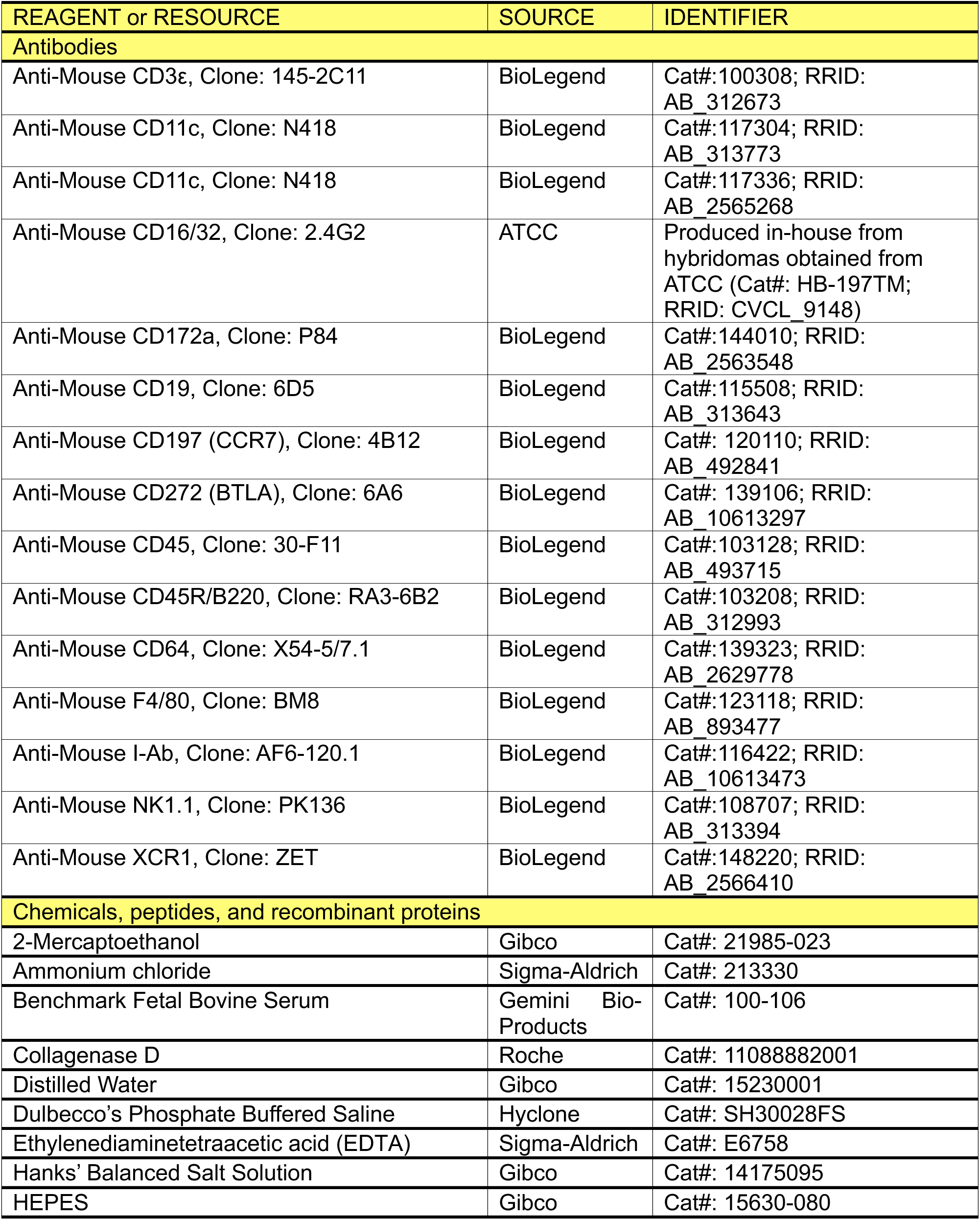
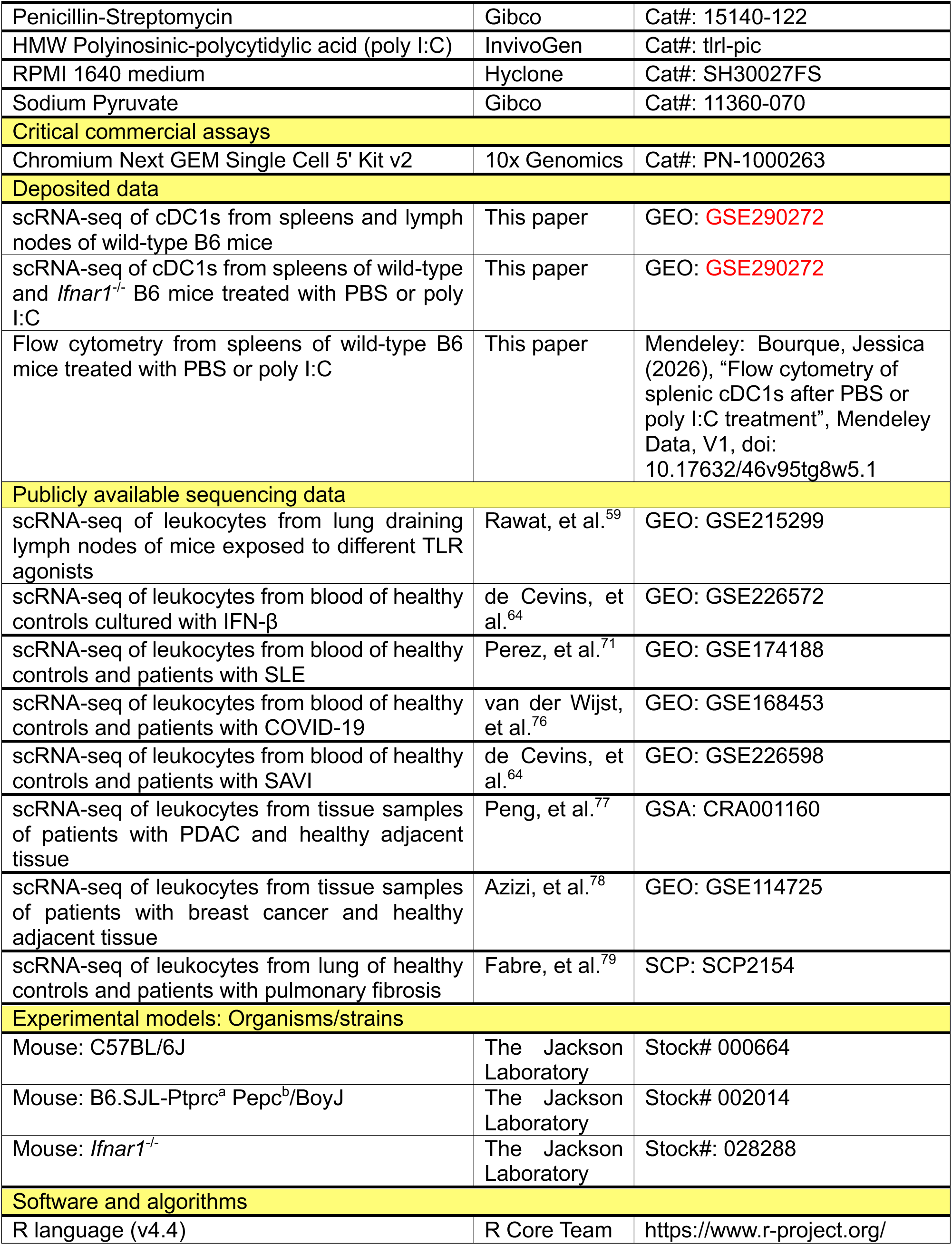
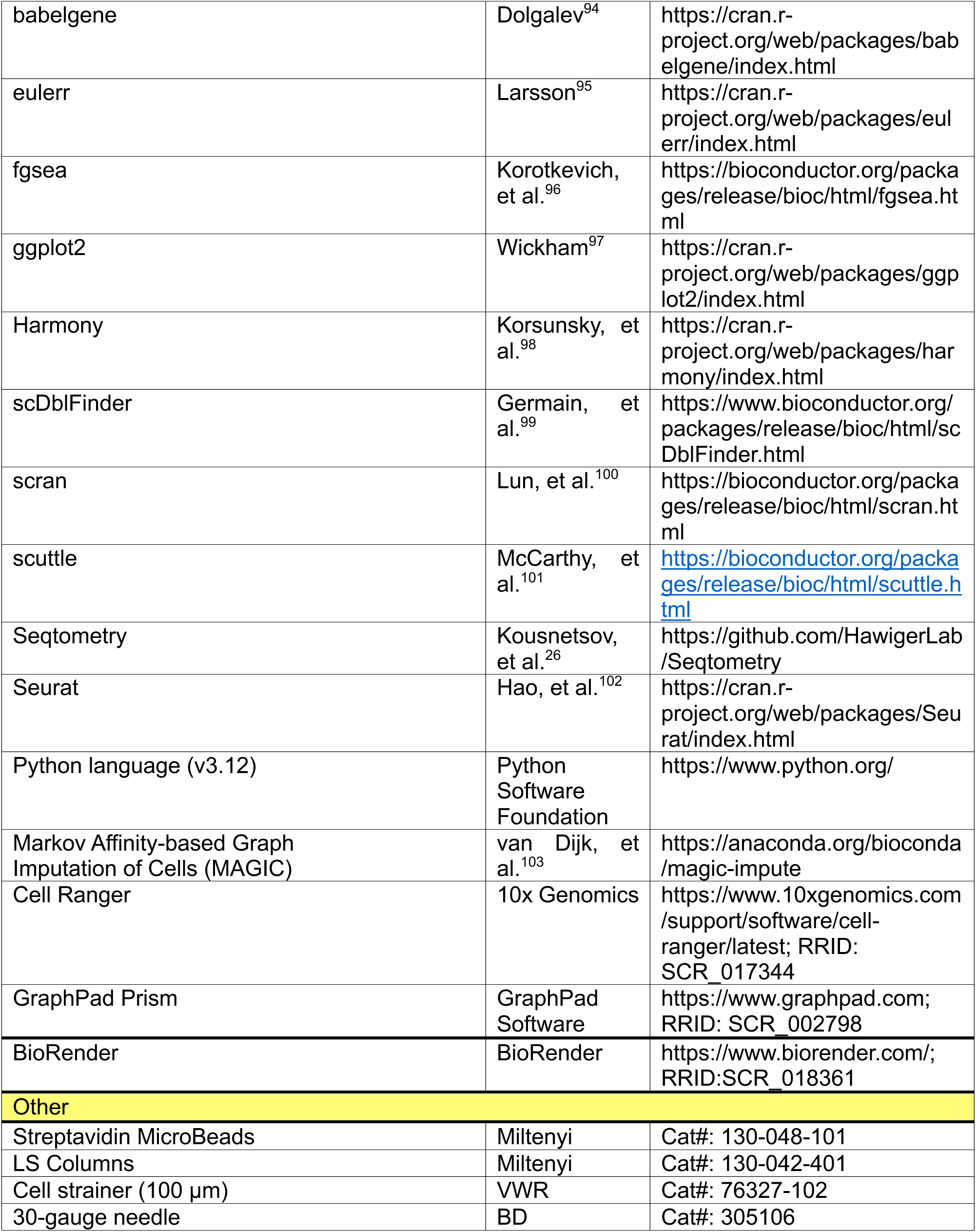

## EXPERIMENTAL MODELS

### Mice

All mice were maintained in our facility under controlled pathogen-free conditions and used in accordance with the guidelines of the Saint Louis University Institutional Animal Care and Use Committee under the ethical approval reference number (Animal Protocol) #2355. All mice including *Ifnar1*^-/-^ ^68^ were from Jackson Laboratory and bred in our colony. 6-to 9-week-old age-matched male and female mice were used in all experiments. Mice were randomly assigned to experimental groups.

## METHODS DETAILS

### In vivo TLR stimulation

High molecular weight polyinosinic-polycytidylic acid (poly I:C) (InvivoGen) was prepared as per the manufacturer’s protocol, diluted in PBS, and intraperitoneally (i.p.) injected at a dose of 50 μg per mouse 12 hours before the start of the experiments.

### cDC preparation and fluorescence-activated cell sorting of cDC1s for single-cell RNA sequencing

Mice were euthanized by CO_2_ inhalation per the recommendations of the Panel on Euthanasia of the American Veterinary Medical Association and Institutional guidelines. cDCs from spleens or peripheral lymph nodes (axial, brachial, and inguinal) of C57BL/6J mice (treated with PBS or Poly I:C 12 hours prior to the experiment) and spleens of *Ifnar1*^-/-^ mice (treated with PBS or Poly I:C 12 hours before harvesting) were mechanically fragmented with forceps and 30-gauge needles (BD), and treated with 2.5 mg/mL Collagenase D (Roche) in RPMI 1640 media (Hyclone) supplemented with 5% fetal bovine serum (FBS) (Gemini Bio), penicillin/streptomycin (100 U/mL), HEPES (10 mM), sodium pyruvate (1 mM), and 2-Mercaptoethanol (55 μM) (all Gibco) at 37°C for 30 min, followed by adding EDTA (10 mM) for 5 minutes at 37°C. After incubation, cells were passed through 100 µm strainers (VWR) and washed using Hanks’ Balanced Salt Solution (Gibco) supplemented with 2% FBS and 1 mM EDTA to obtain single-cell suspensions. Following lysis of red blood cells (RBCs) with ammonium chloride solution (0.16 M) for 5 minutes at room temperature, single-cell suspensions were washed with Dulbecco’s phosphate-buffered saline (PBS) (Hyclone) supplemented with 5% FBS. Cells were then stained with Fc block (anti-CD16/32 clone 2.4G2 – purified in-house) for 15 minutes at room temperature. For surface staining, cells were subsequently incubated with fluorochrome-conjugated antibodies diluted in PBS supplemented with 2% FBS for 30 minutes on ice. After incubation, cells were washed twice and strained through 35 µm strainers prior to acquisition. To stain the surface expression of CCR7, cells were incubated with anti-CCR7 fluorochrome-conjugated diluted in PBS supplemented with 2% FBS for 30 minutes at 37°C prior to incubating with other surface marker antibodies. All samples were acquired on a BDFortessa (BD).

For fluorescence-activated cell sorting, after RBC lysis and washing, pooled cells (from 7-10 mice for each condition) from each organ were incubated with fluorochrome-conjugated antibodies as well as anti-CD11c biotinylated antibody for 30 minutes on ice. Cells were washed and incubated with streptavidin microbeads (Miltenyi) for 15 minutes at 4°C. CD11c^+^ cells were positively selected using LS columns (Miltenyi). The enriched cells were sorted on a FACSAriaIII (BD) or Aria Fusion (BD) and gated as singlet, CD45^+^, Lineage (CD3ε, CD19, B220, NK1.1)^neg^, F4/80^neg^, CD64^neg^, I-A^b+^, CD11c^+^, XCR1^+^, CD172a^lo^ (see Figure S1A for gating strategy). Following sorting, the cells were spun down and resuspended in RPMI medium supplemented with 10% FBS as 1,000,000 cells/mL.

### Chromium single cell 5’ library construction

Single-cell suspensions of sorted cDC1s as above were loaded on a Chromium Single Cell Controller instrument (10x Genomics) to generate single-cell gel beads in emulsion (GEMs). Single-cell RNA sequencing libraries were prepared using the Chromium Single Cell 5′ Library & Gel Bead Kit (v2 Chemistry Dual Index), as per the manufacturer’s instructions. The libraries were then sequenced to a depth of 50 million reads per sample on an Illumina Nova-Seq 6000 by the Washington University in St. Louis Genome Technology Access Center at McDonnell Genome Institute.

## DATA ANALYSIS

### Processing and analysis of scRNA-seq data

Reads from scRNA-seq data were aligned to the mm10 mouse reference transcriptome using 10x Genomics’ Cell Ranger 7.0.1.^104^ All datasets were filtered to exclude doublets and low-quality cells using scDblFinder^99^ and scuttle^101^ libraries, respectively. For publicly available scRNA-seq data obtained from the respective sources listed in the Key Resources Table, the processed data was restricted to dendritic cells and their subsets based on the authors’ clustering and annotations. Expression data from each dataset was then normalized using a LogCP10K transform. Following normalization, data were further analyzed either with a conventional clustering pipeline (using the Seurat package^102^) or Seqtometry.^26,28^

Within Seurat, the top 2000 highly variable features were used to compute the leading 50 principal components. Using the leading 10 principal components, a shared nearest neighbor (SNN) graph was computed, and a two-dimensional embedding was performed using the Uniform Manifold Approximation Projection (UMAP) algorithm.^105^ The SNN graph was then used to compute clusters with the Leiden algorithm,^106^ where the resolution parameter was set to 0.4. To visualize expression of individual genes, the Nebulosa library^107^ was used to compute weighted expression densities of top 5% expressing cells that were then overlaid atop UMAP embeddings.

To analyze data using Seqtometry, normalized data was first imputed using the MAGIC algorithm.^103^ Imputed gene expression matrices were scored using the Seqtometry algorithm^26^ with the scoring results from individual transcriptomic signatures, then displayed using biaxial plots or histograms as described below. Additionally, individual signature scores were overlaid on some plots. To visualize expression of individual genes, their imputed expression values were overlaid on some plots. To quantify the joint enrichments of cells from individual subjects presented in biaxial plots, each cell’s scores were orthogonally projected onto the diagonal joint enrichment axis. Then, each subject’s overall enrichment was summarized as the median directed distance from the central point (0.5, 0.5) for all cells belonging to that subject.

### Signature generation

Signatures were generated by performing differential expression tests between cells within an indicated region versus all other cells outside of a region or between all cells from two different datasets. Specifically, a Wilcoxon rank sum test was performed using the Scran library^100^, where genes included in a signature had a log-fold change greater than or equal to 1.8 and an adjusted p-value less than 0.05. Additional refined signatures were then derived from the intersection of such initial signatures and represented as Venn diagrams. To make use of signatures derived from murine data in the context of human data, murine signatures were converted into their human equivalents by using human orthologs for murine genes, where orthologs were retrieved using the babelgene library^94^. All signatures, including those retrieved from the literature or the molecular signatures database (MSigDB)^53^ are compiled in Table 1 (mouse) and Table 2 (human).

**Table 1.**
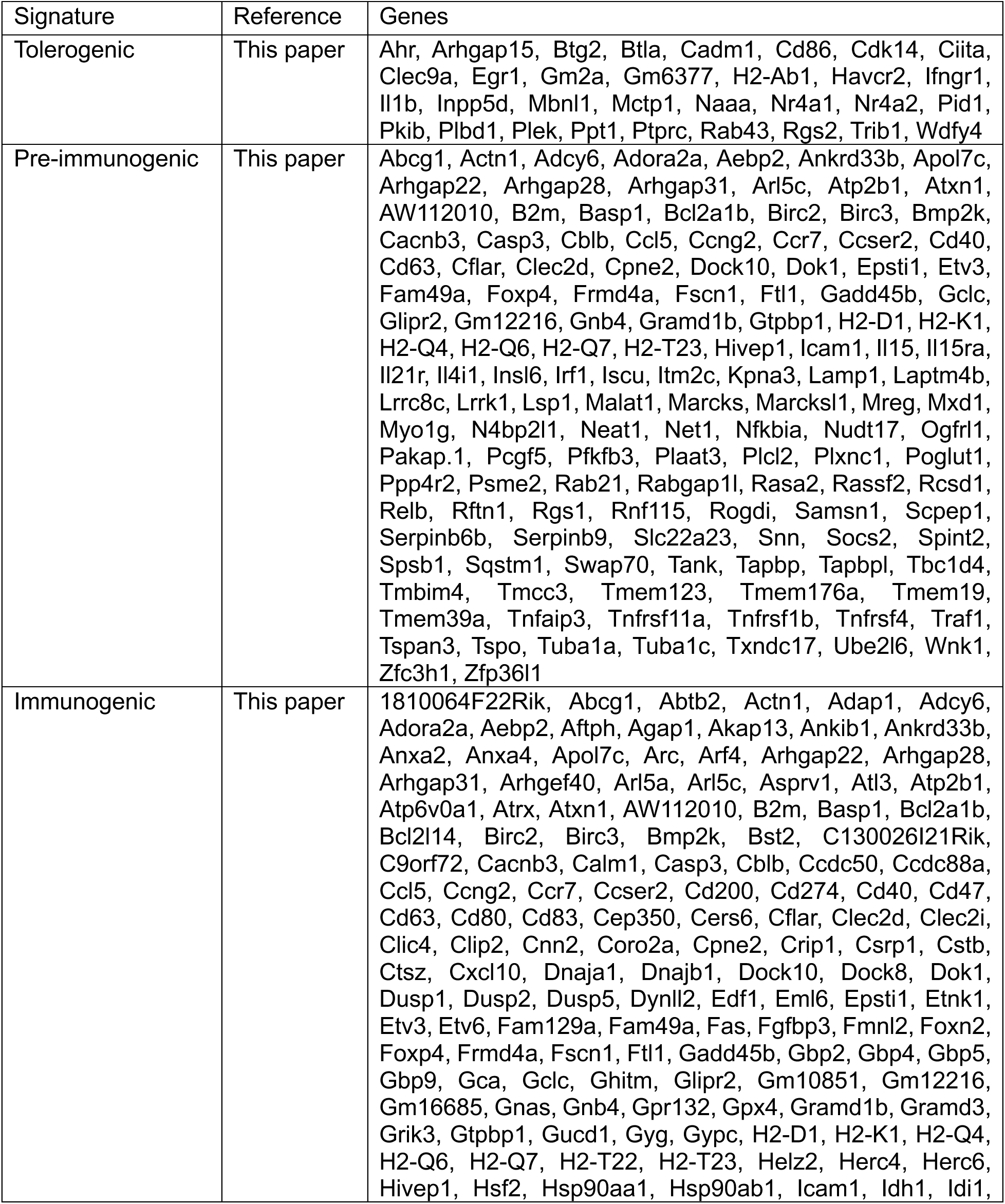
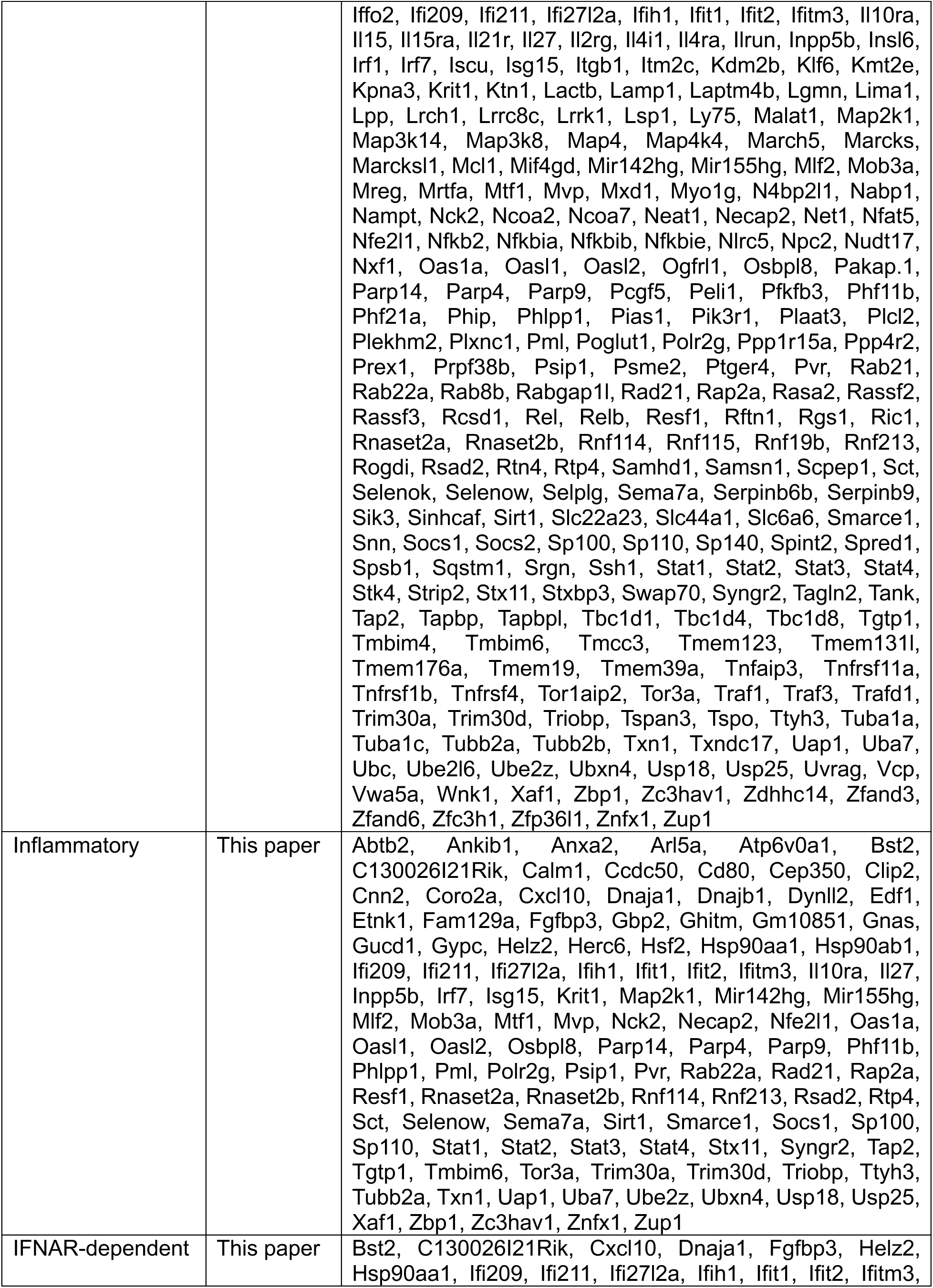
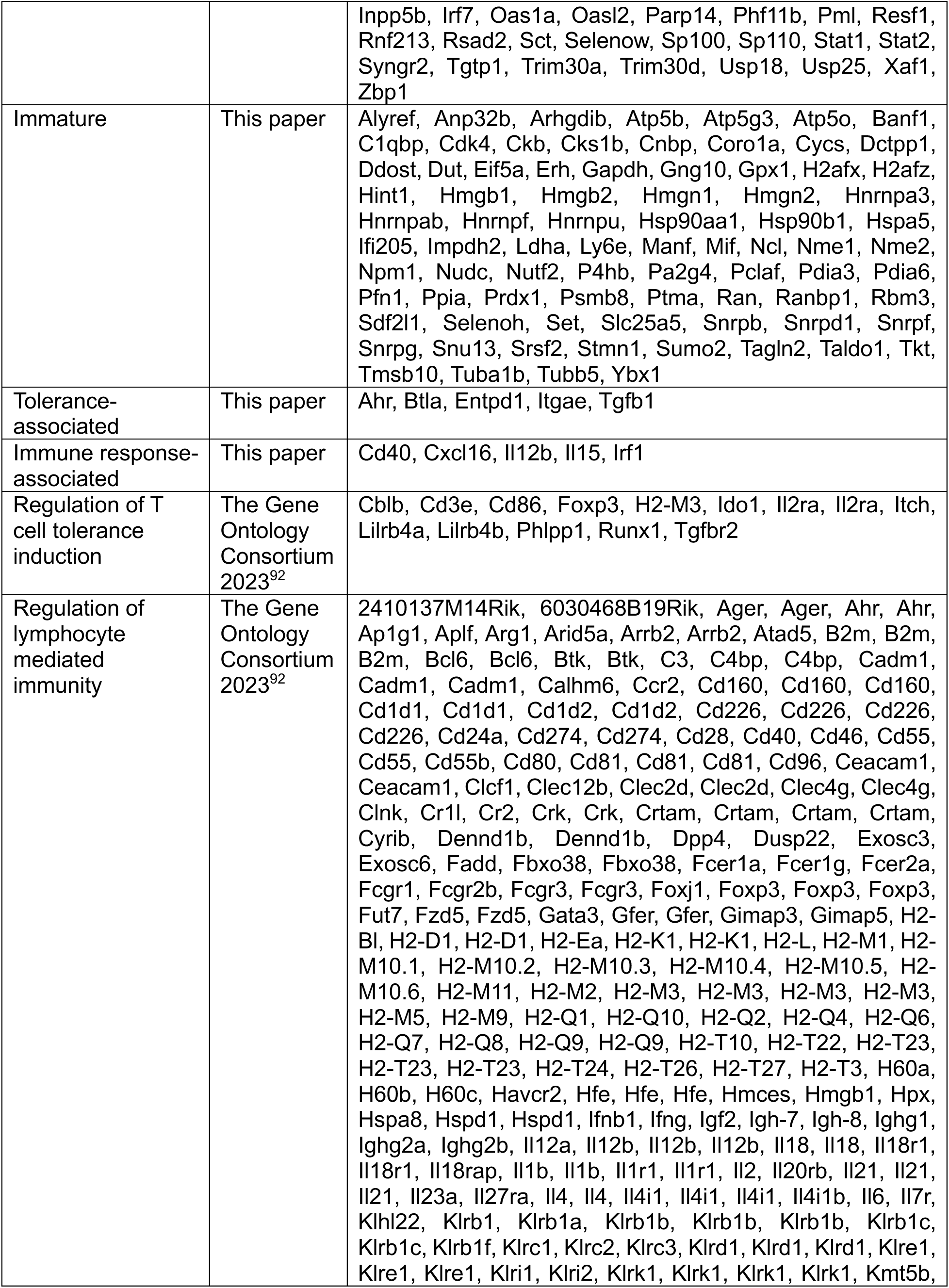
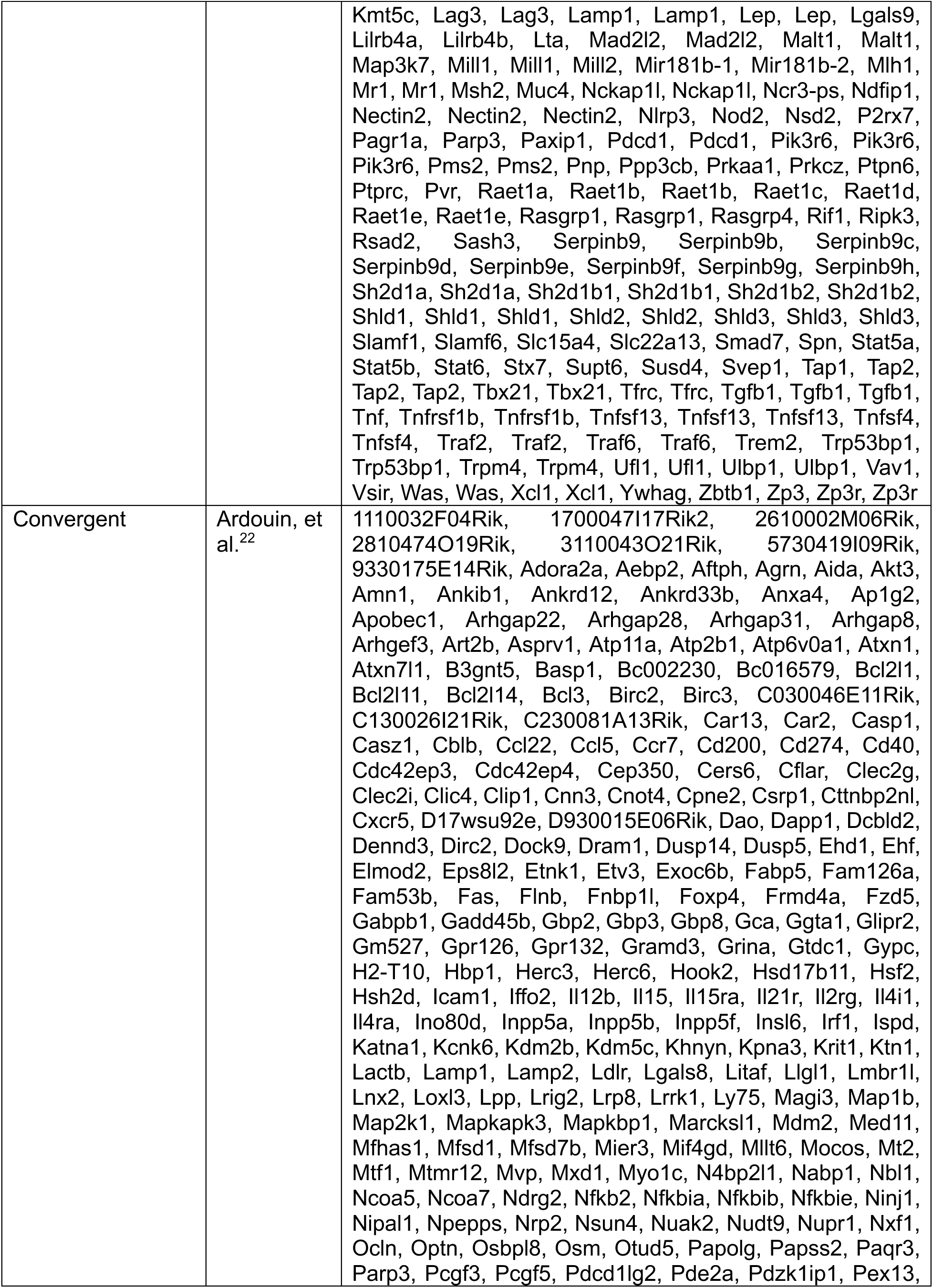
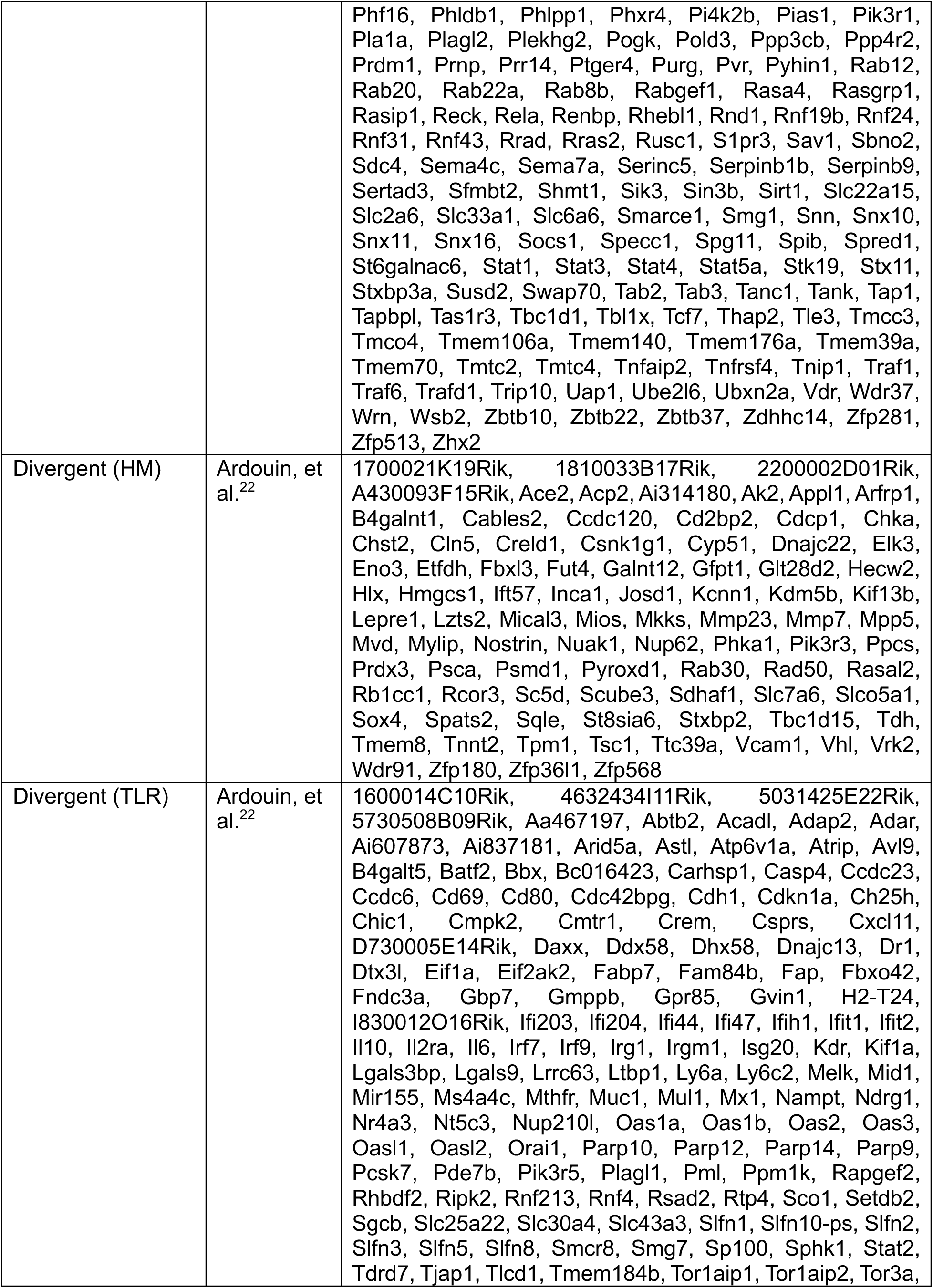
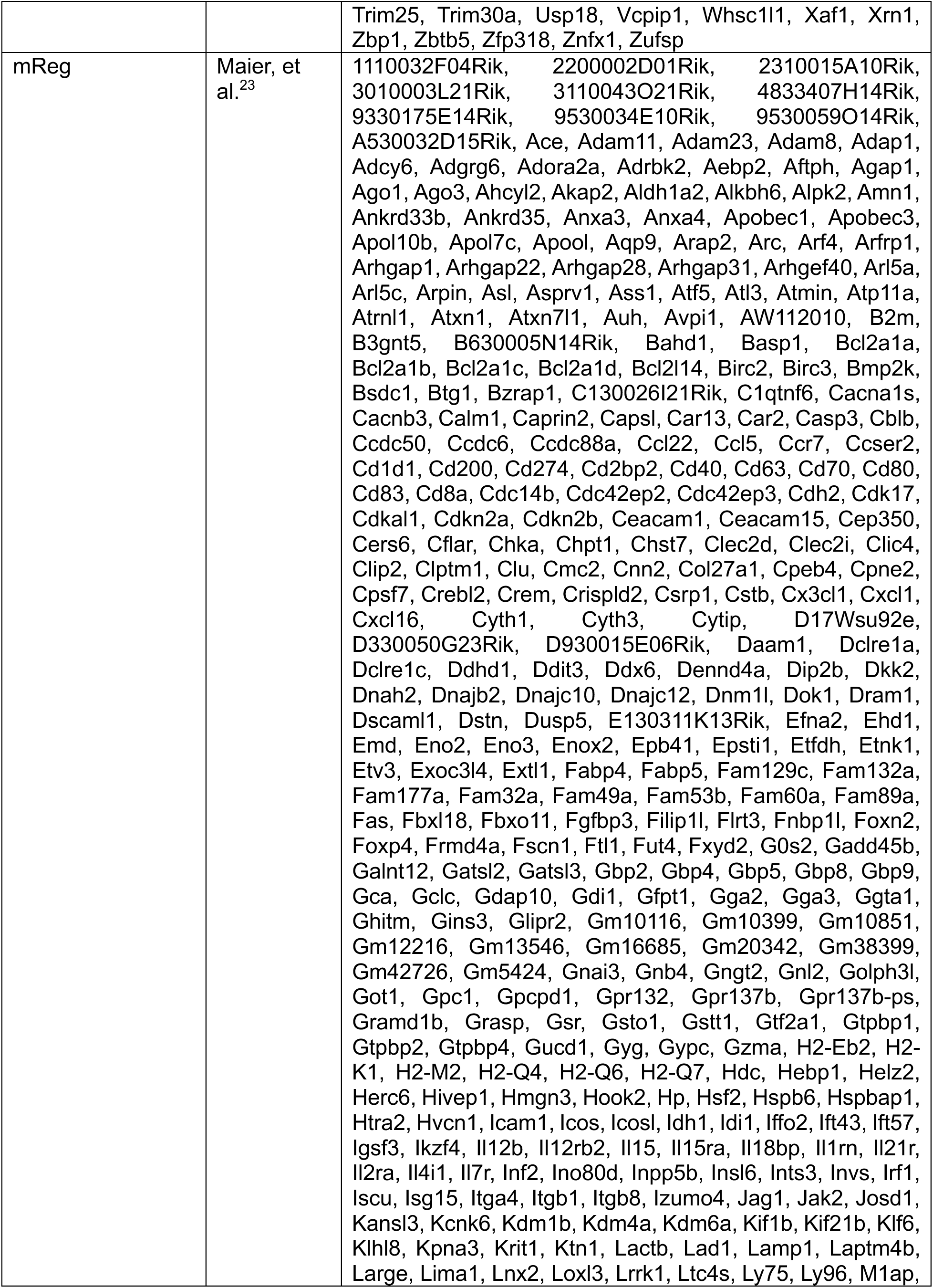
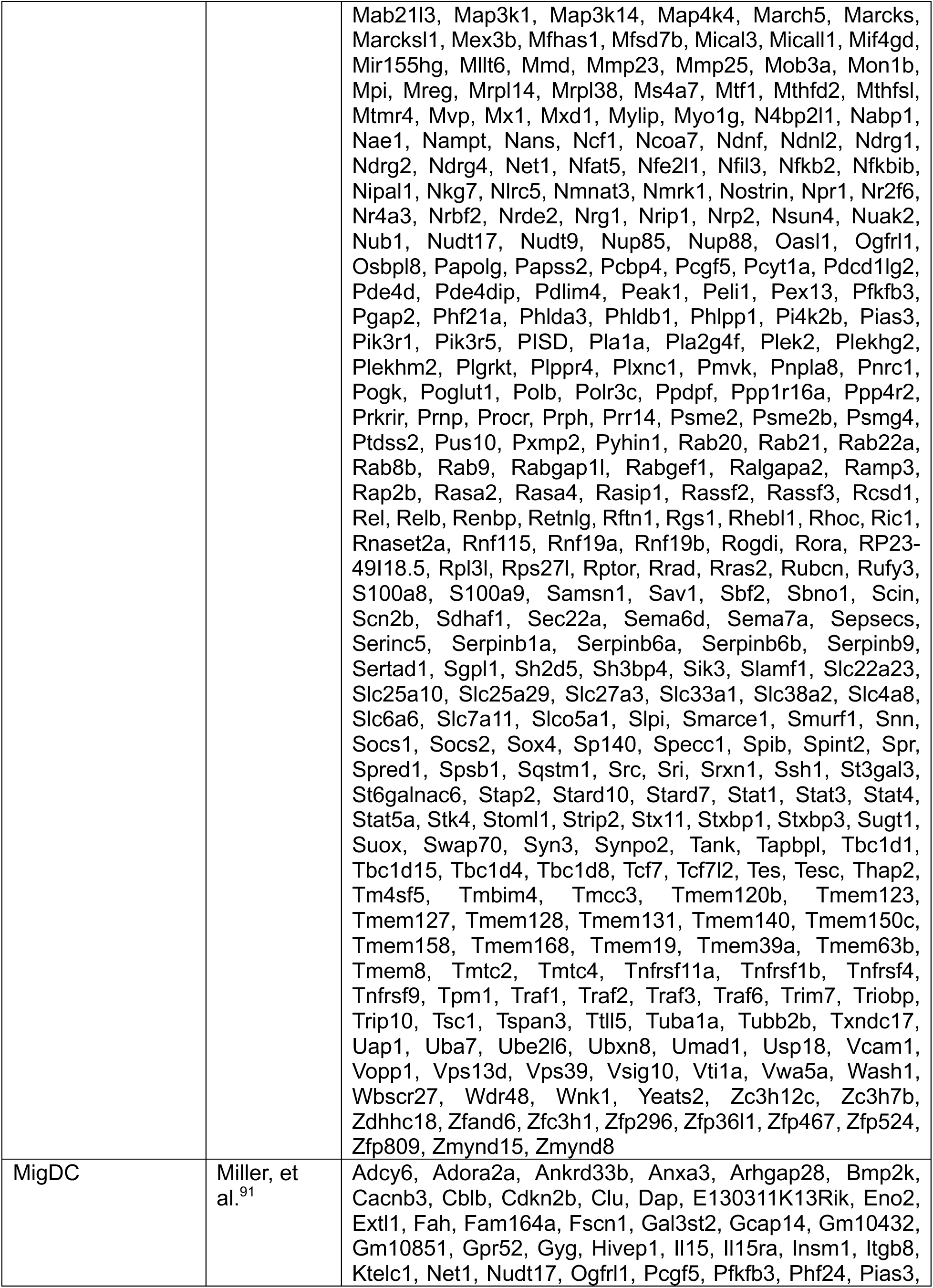
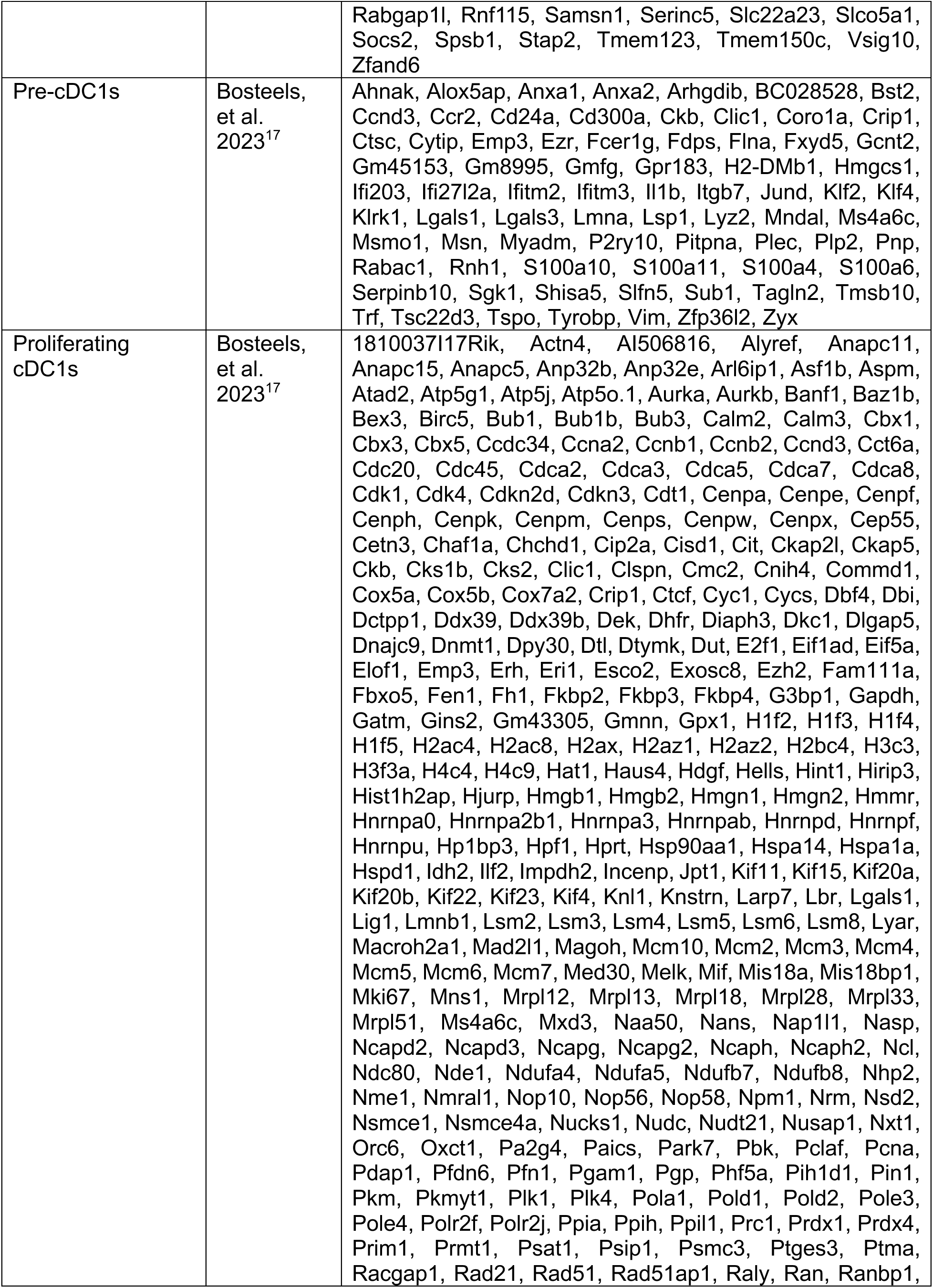
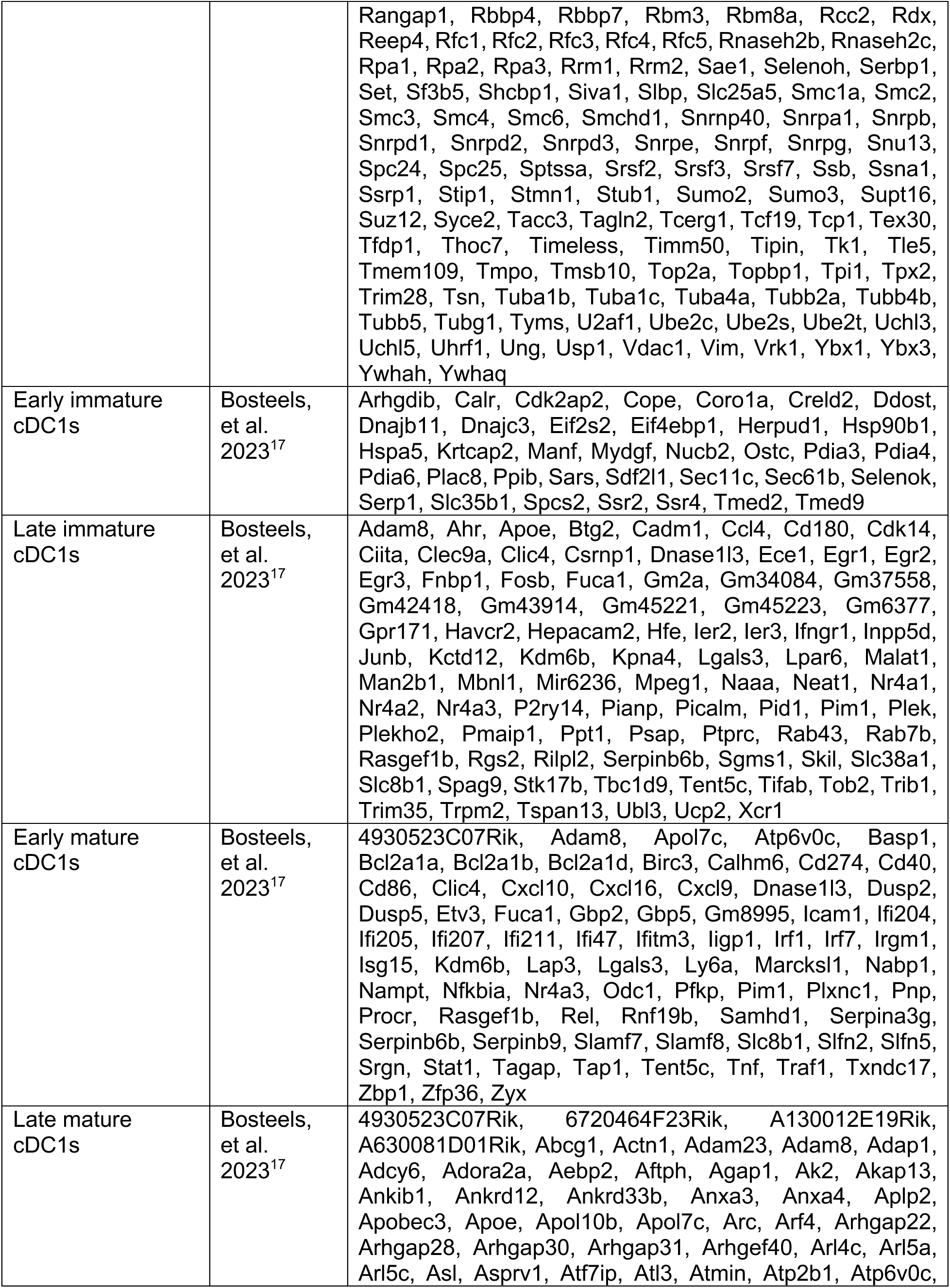
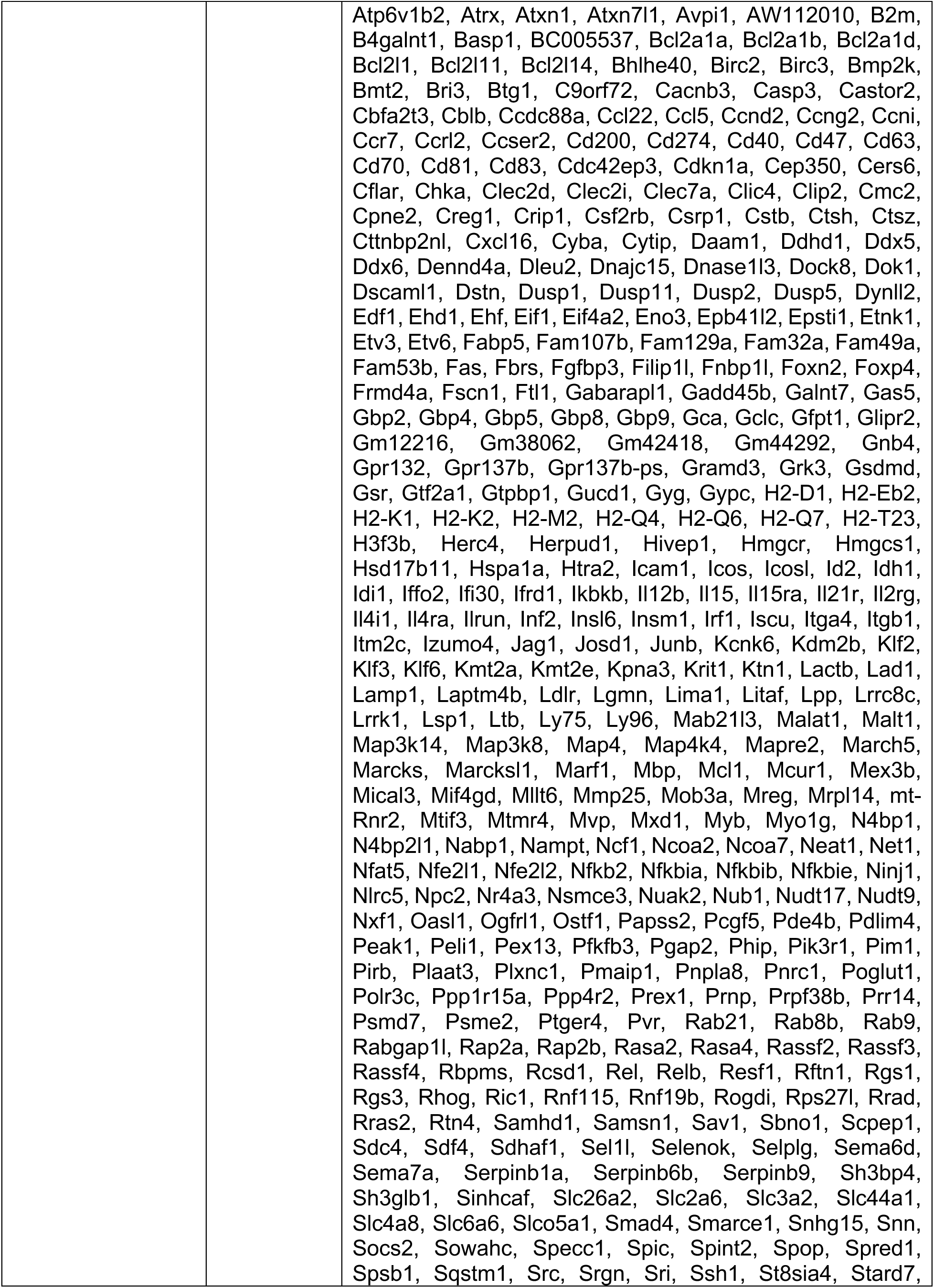
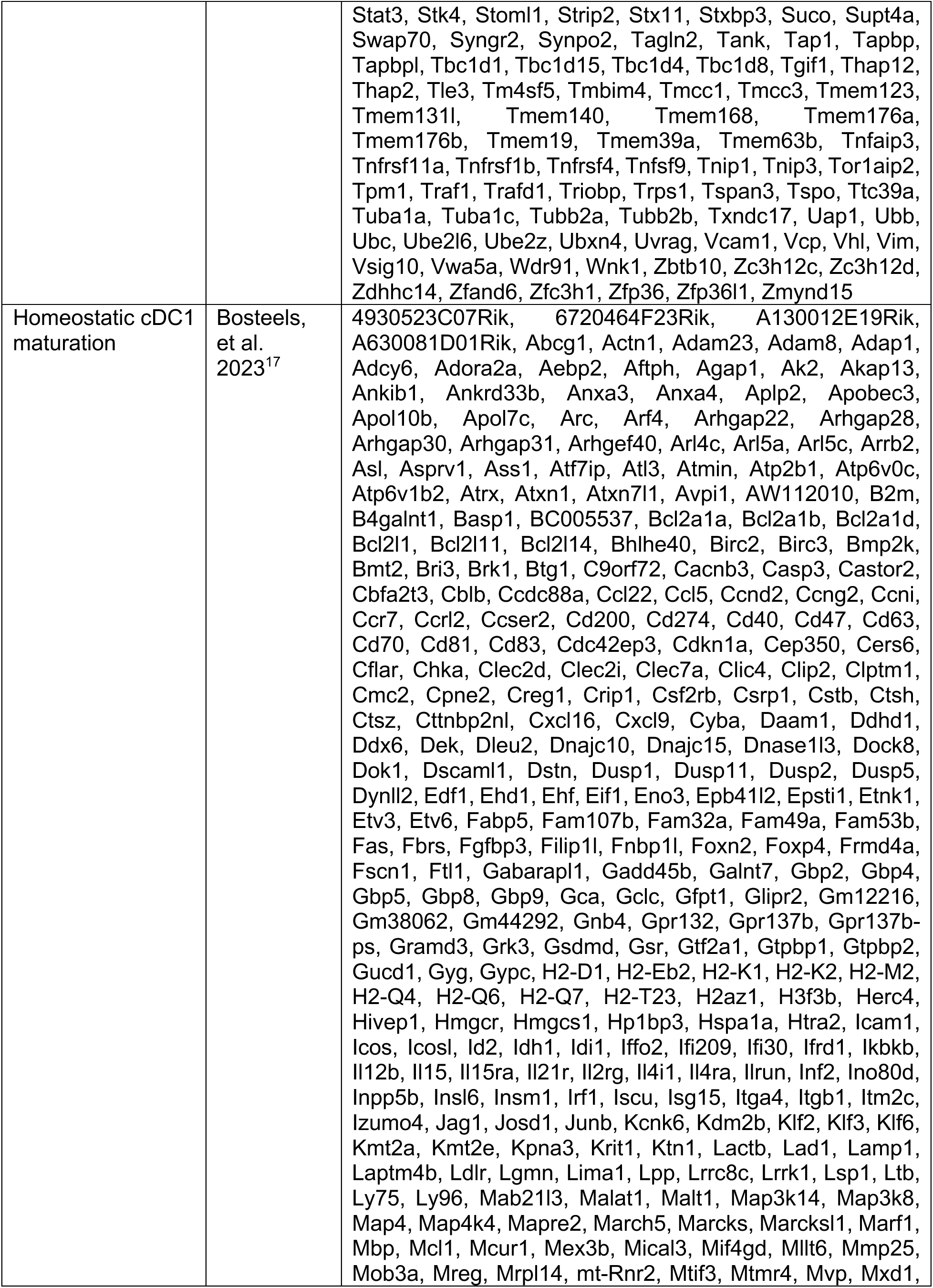
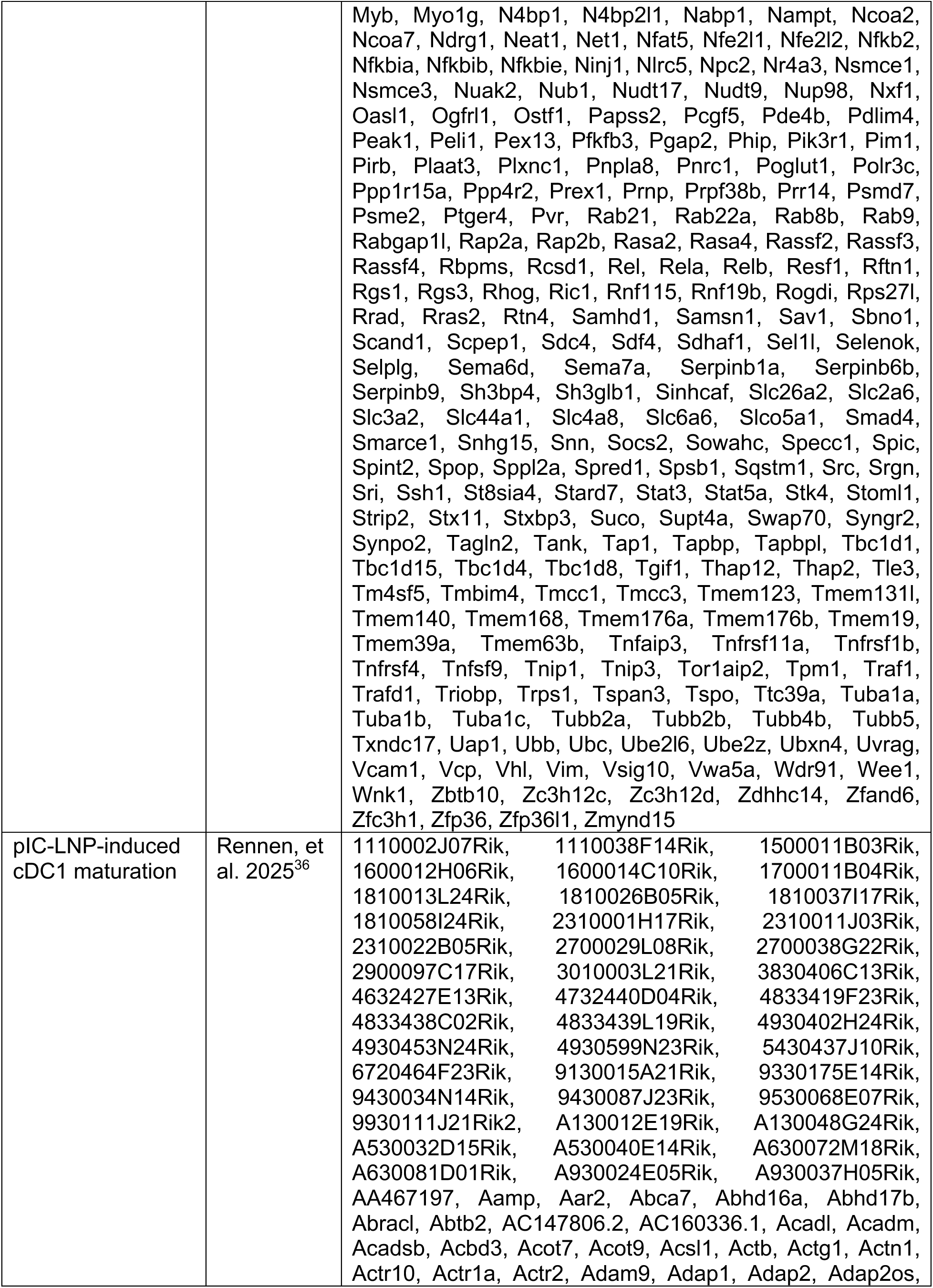
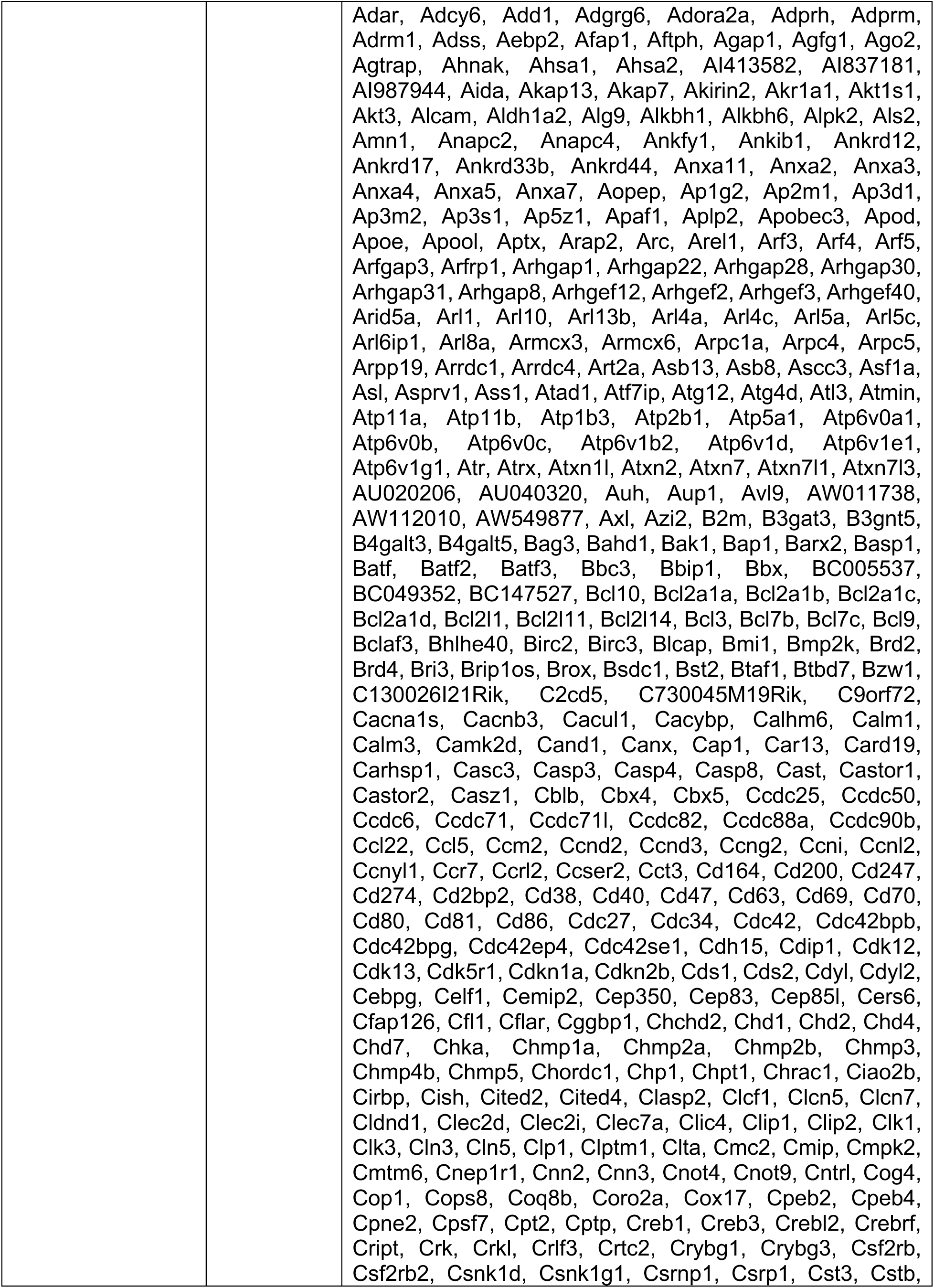
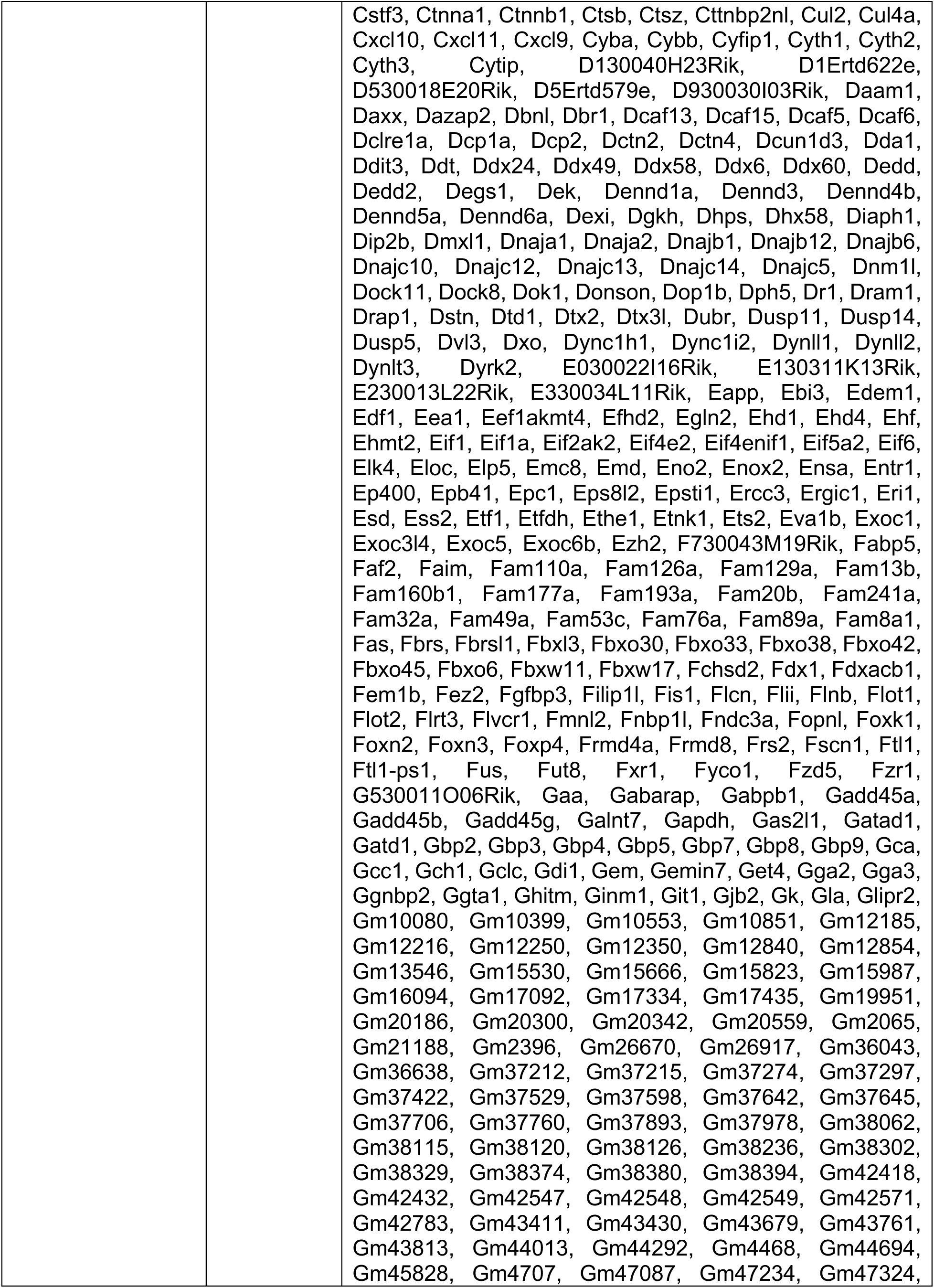
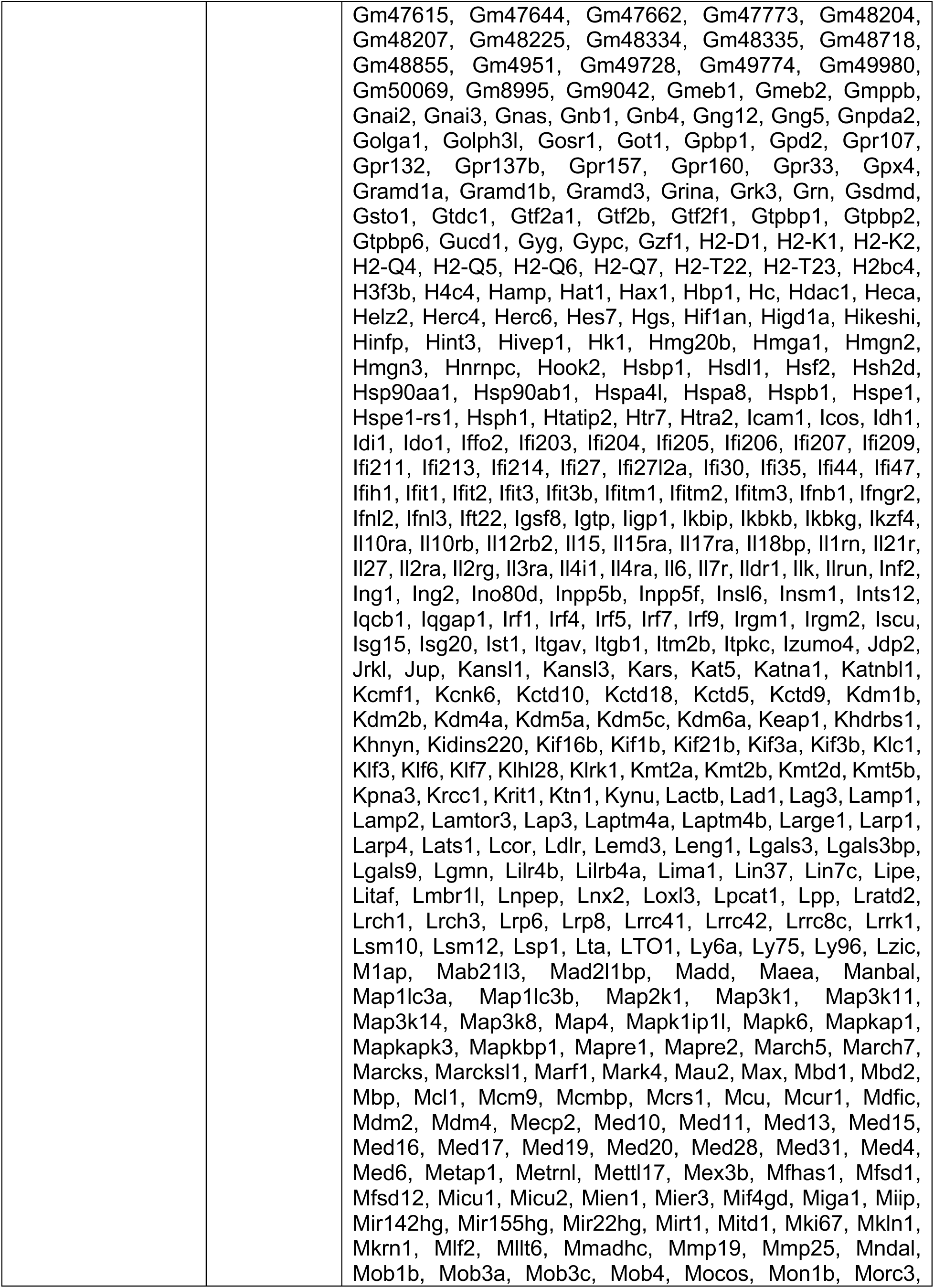
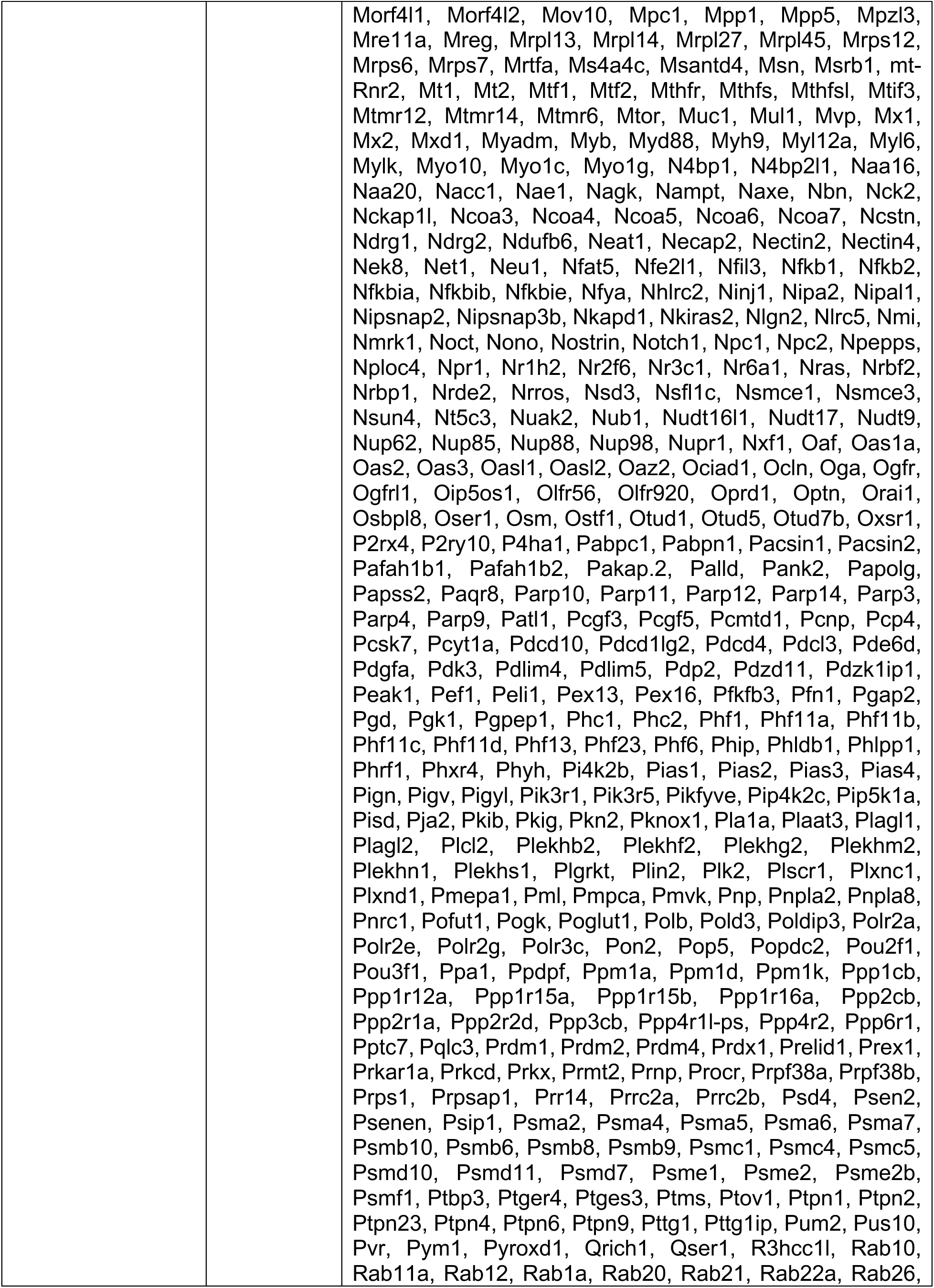
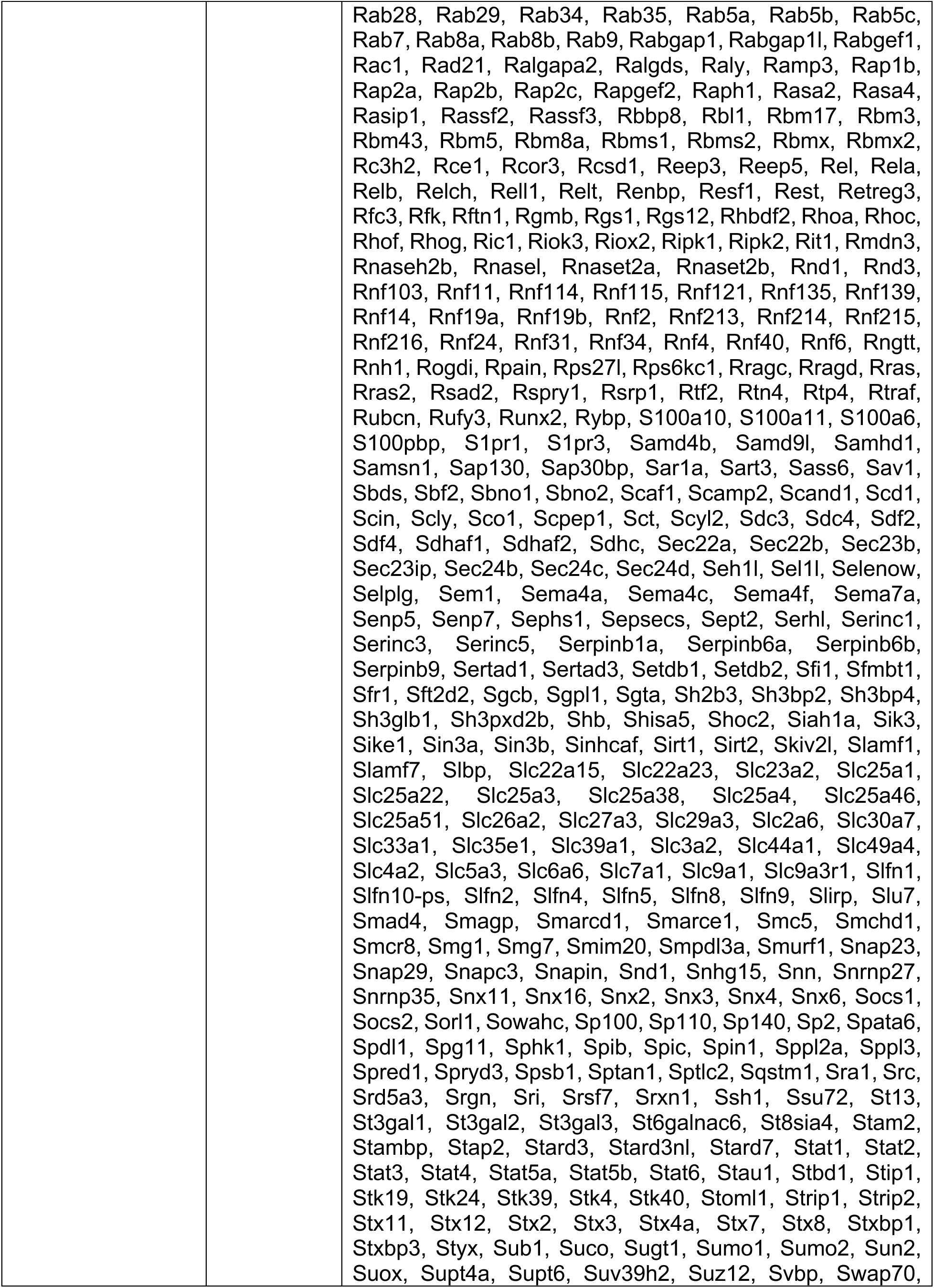
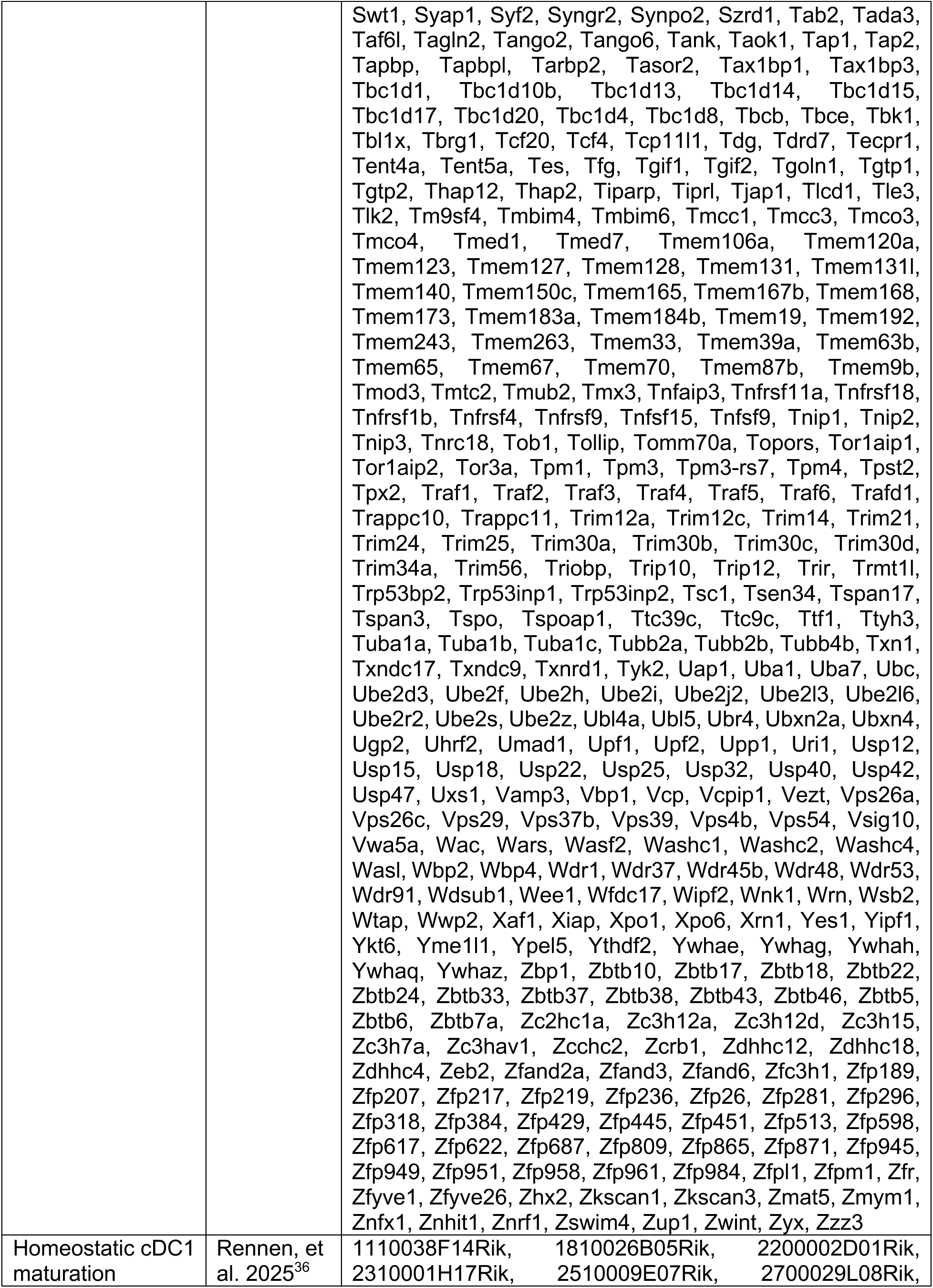
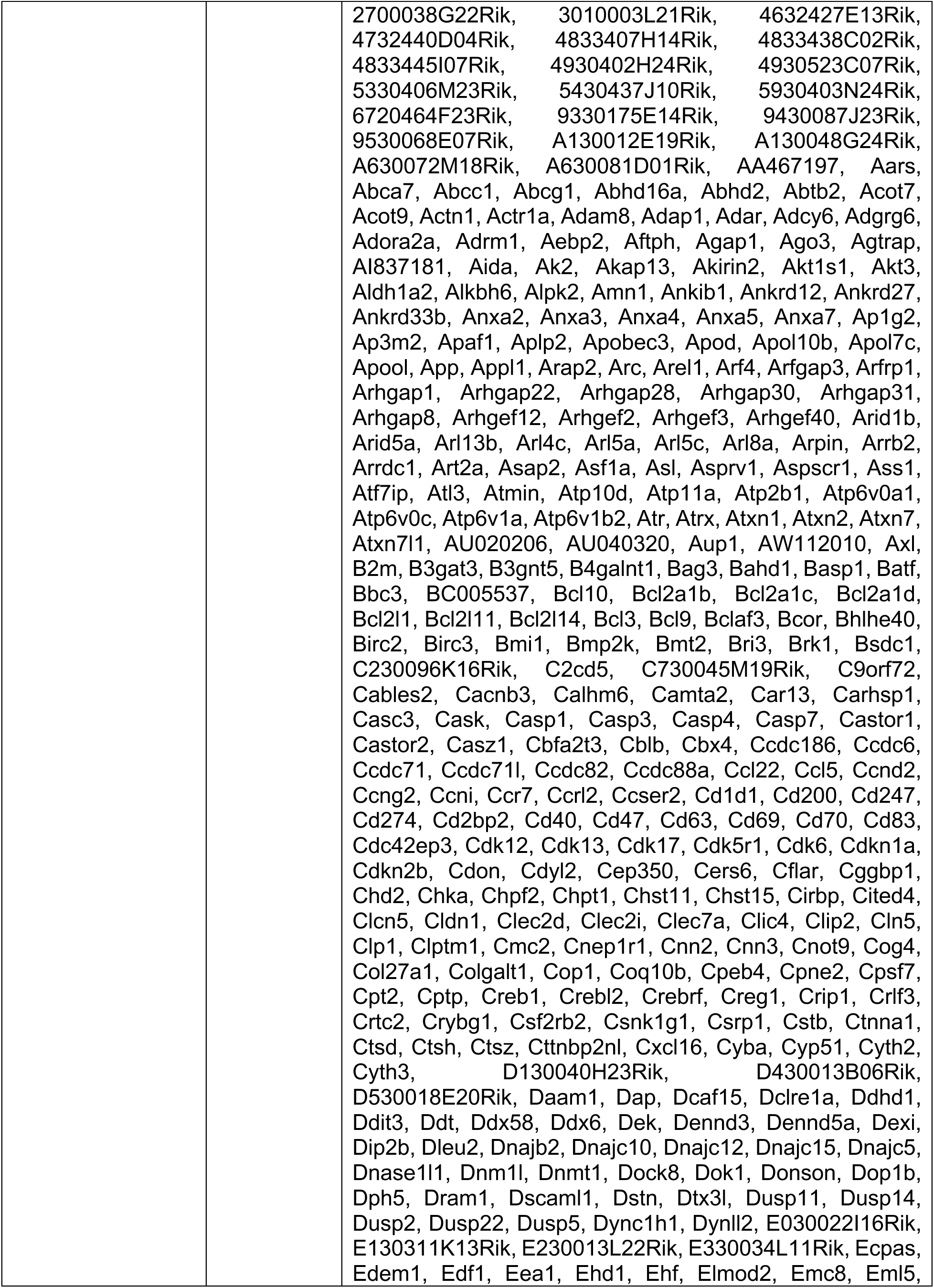
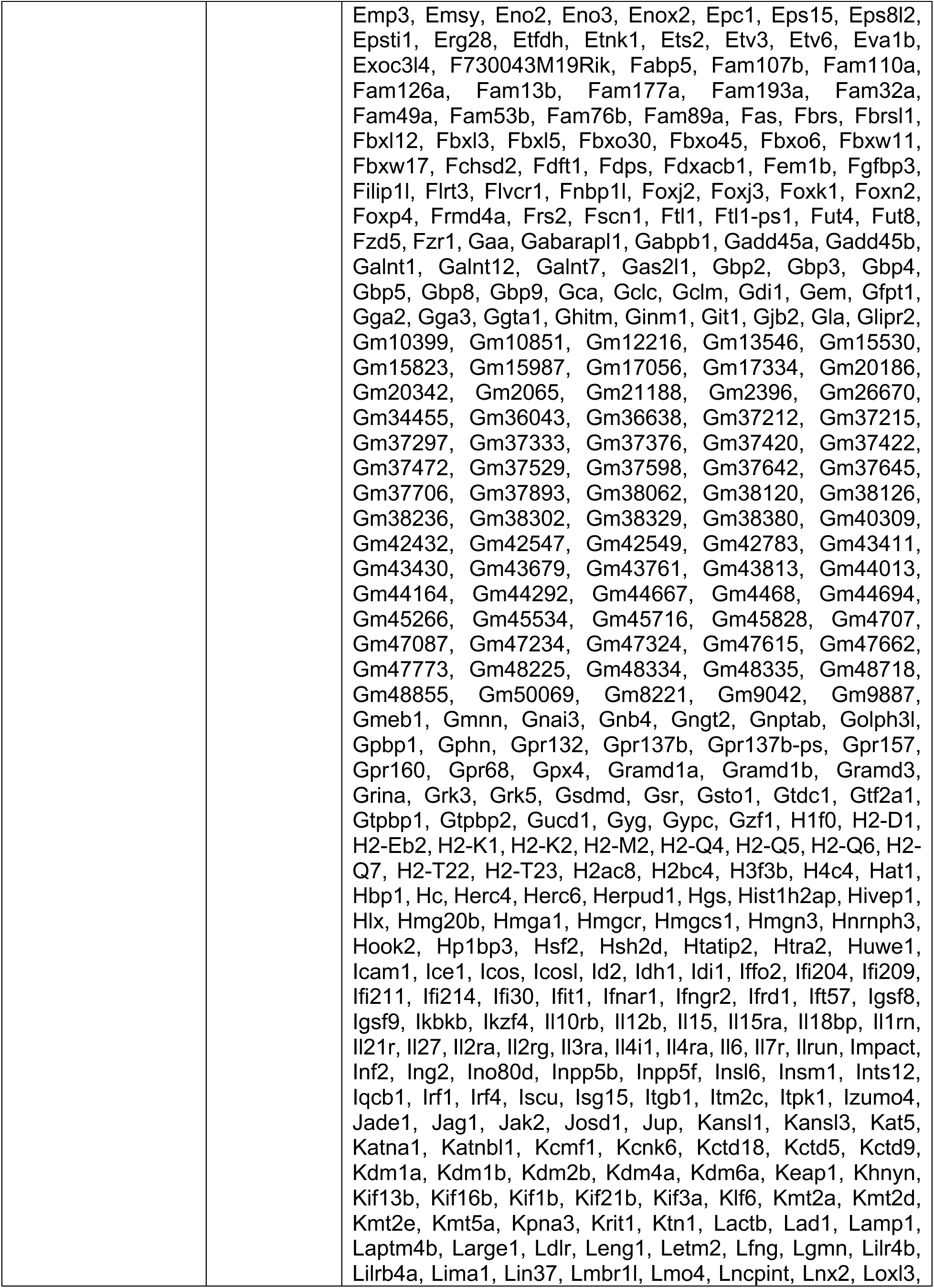
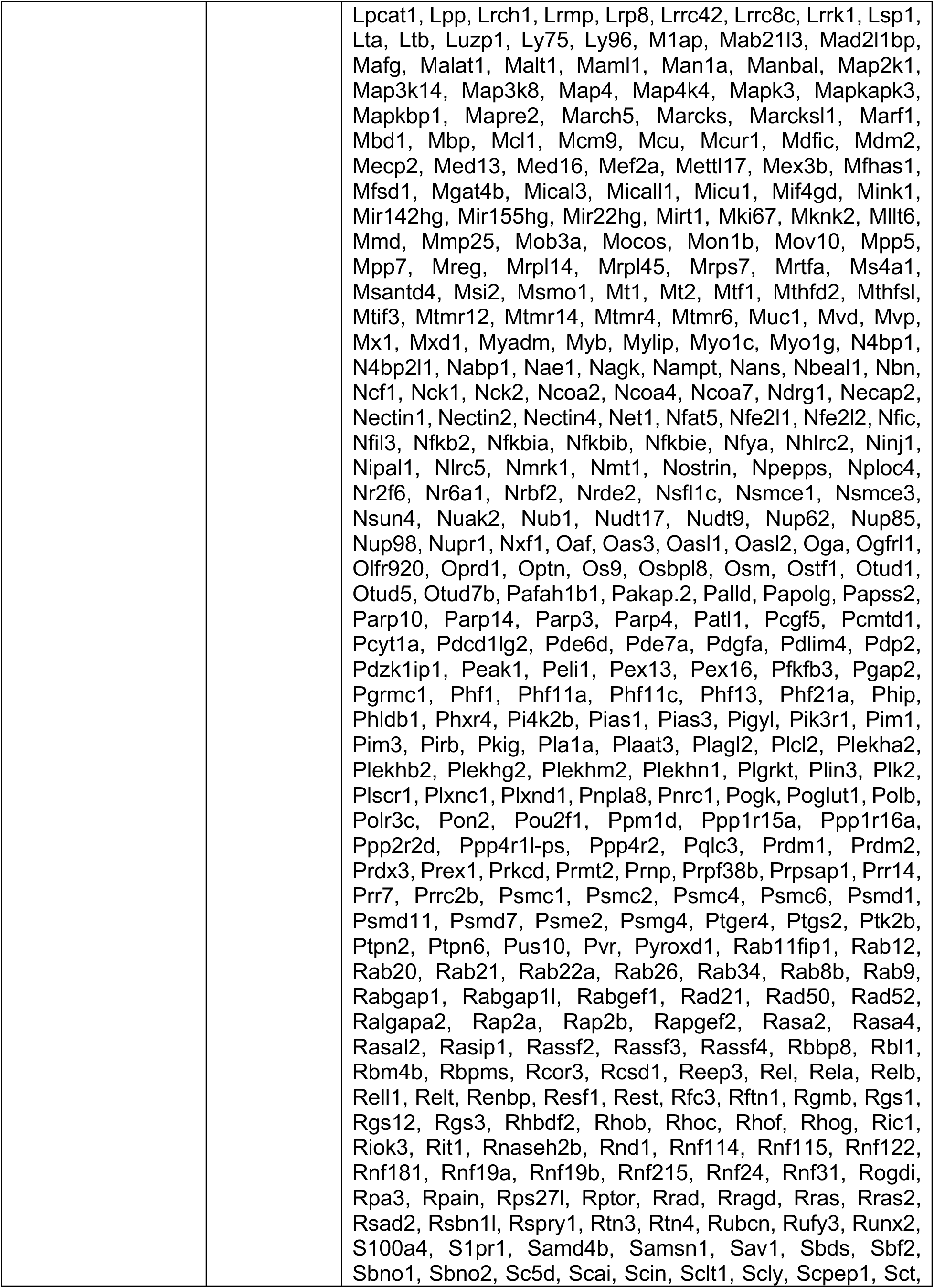
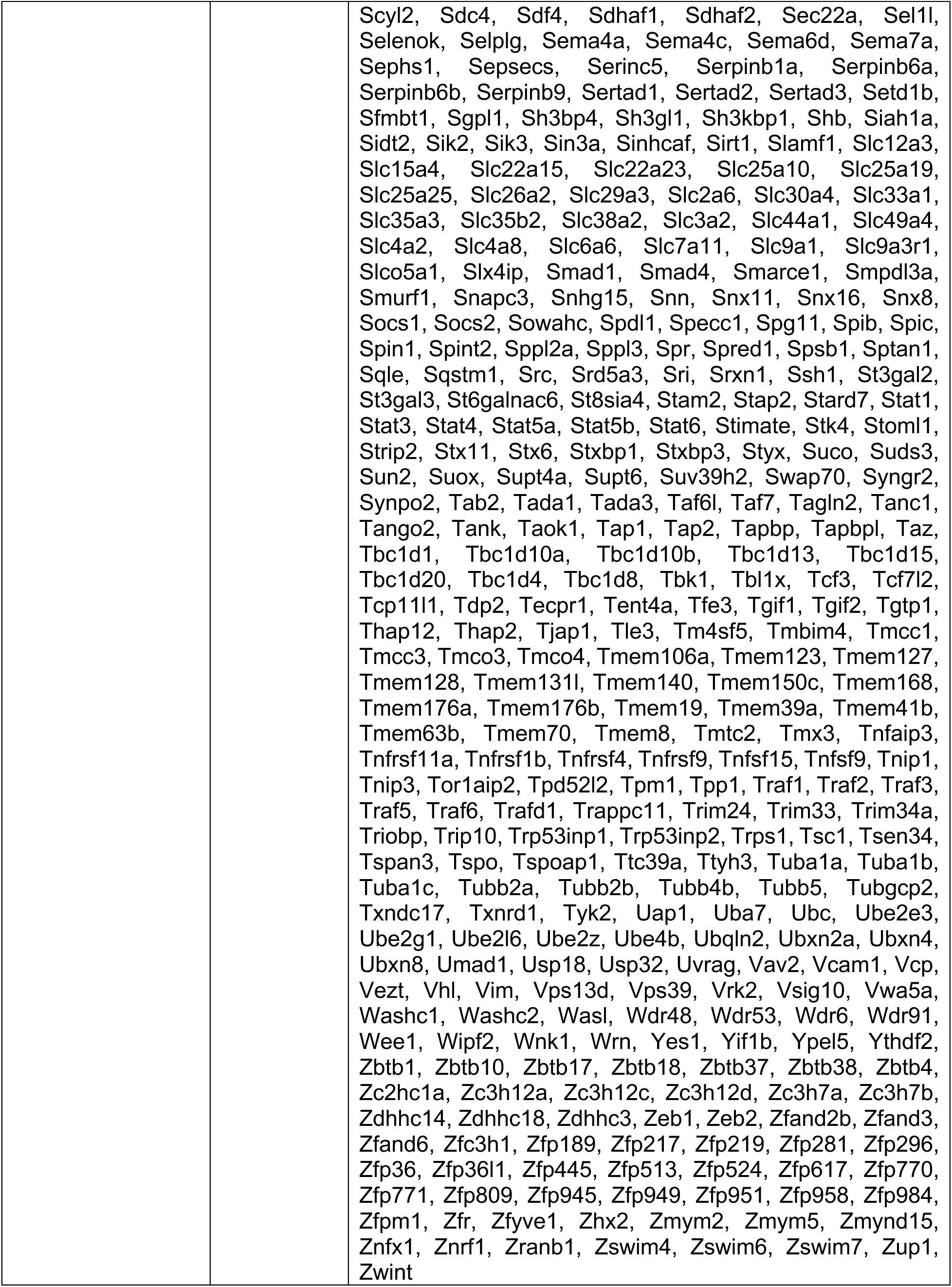
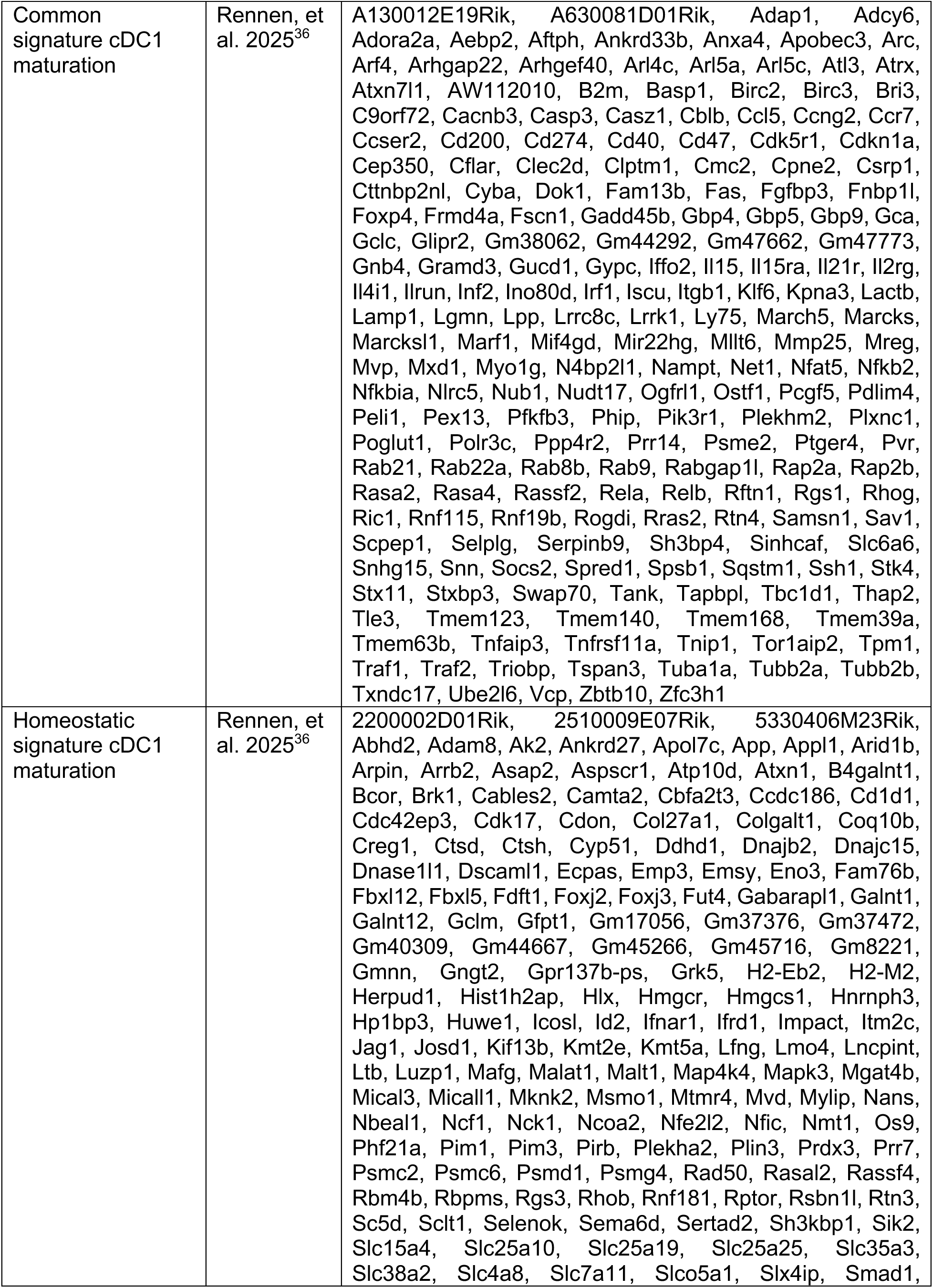
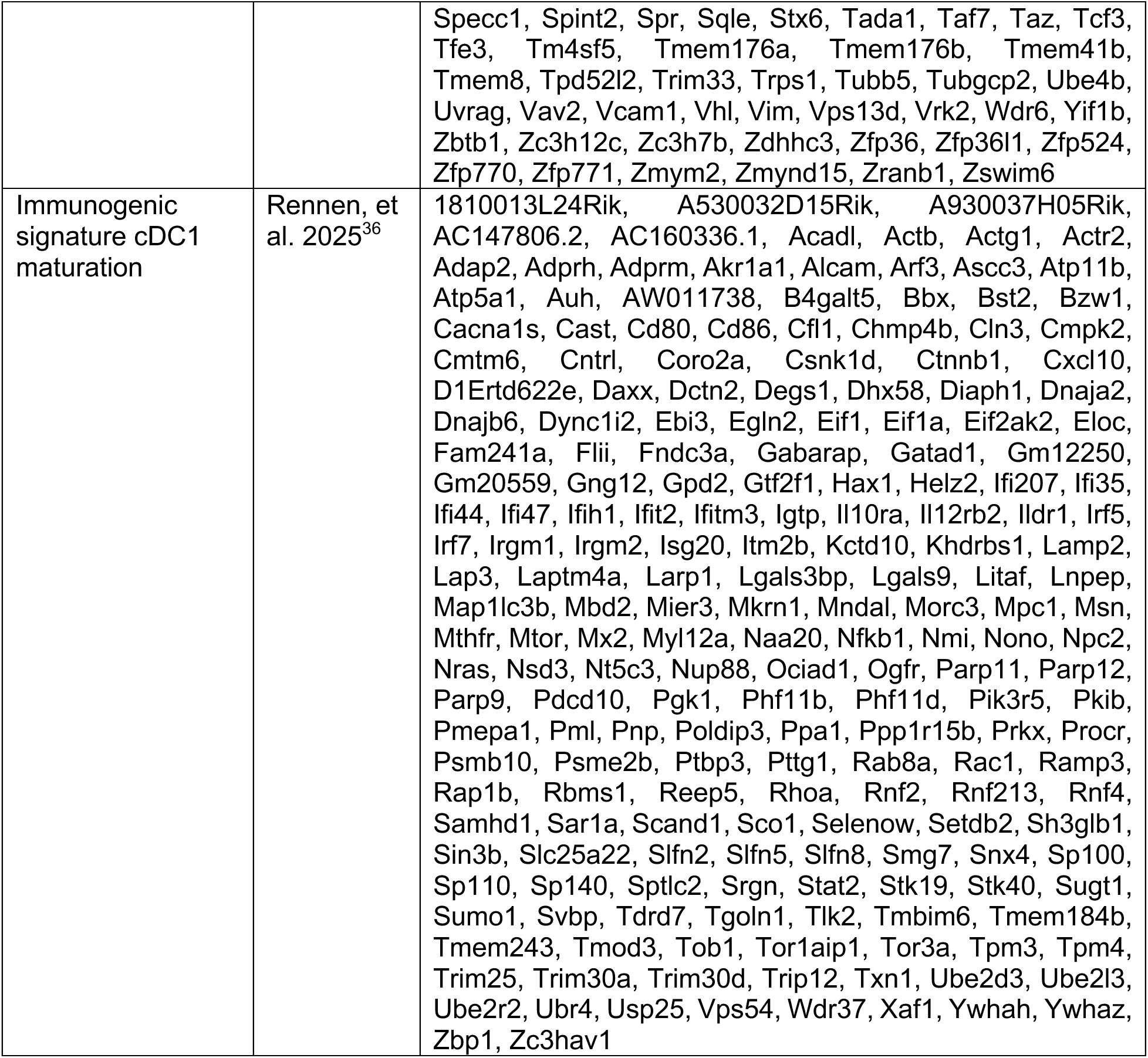
Mouse gene signatures used in this paper. HM: homeostatic maturation; TLR: Toll-like receptor; mReg: mature DCs enriched in immunoregulatory molecules; migDC: migratory DC.

**Table 2.**
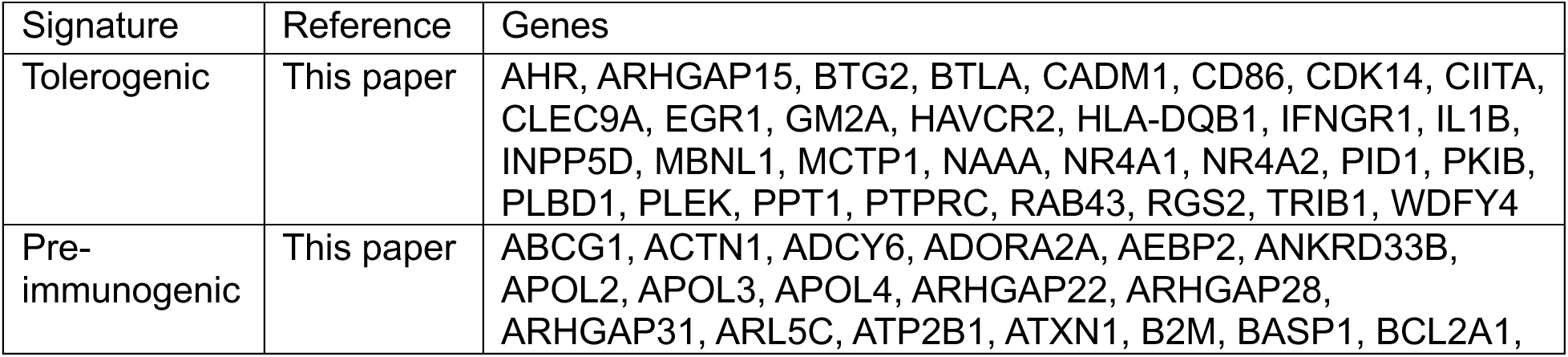
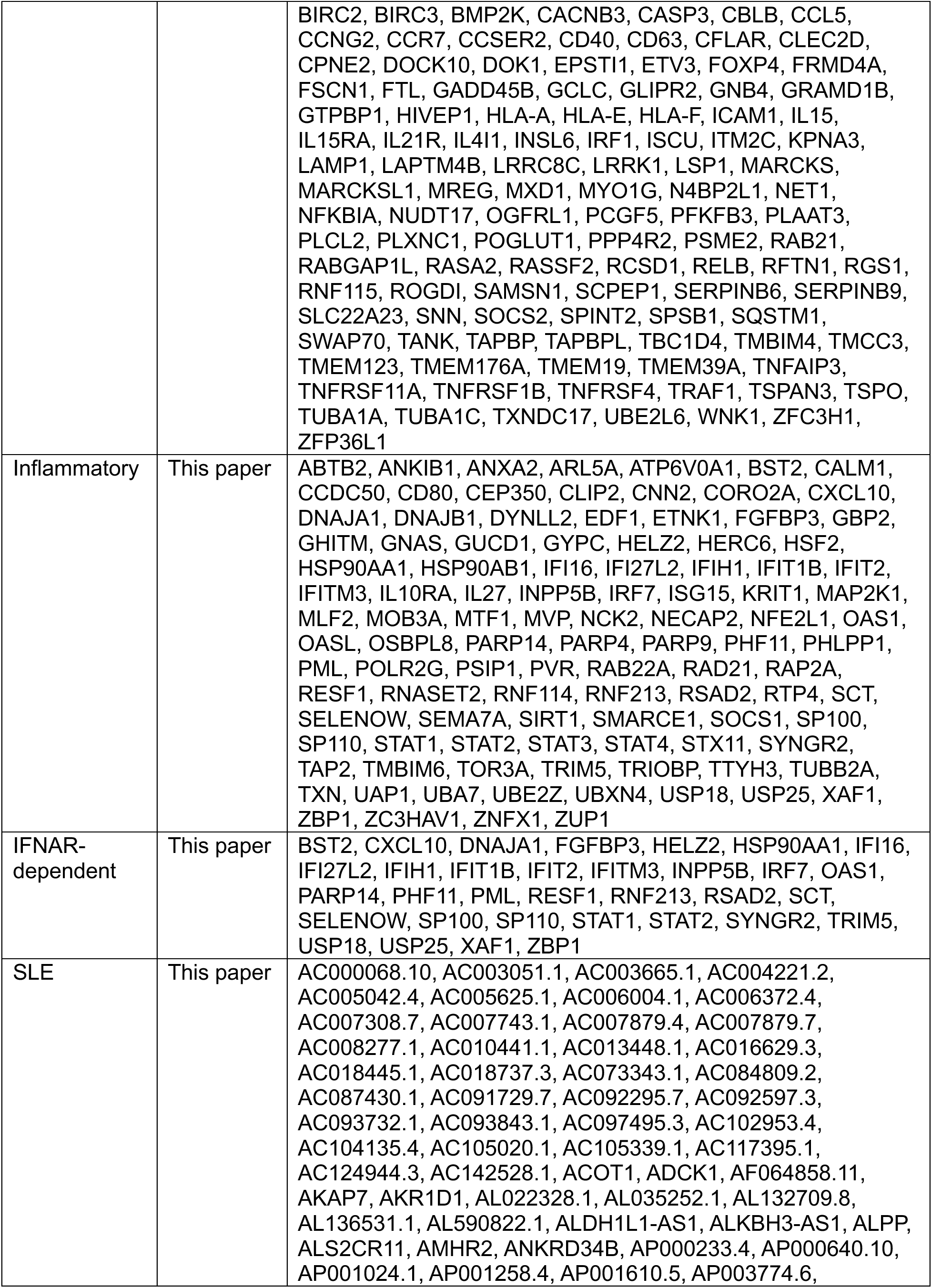
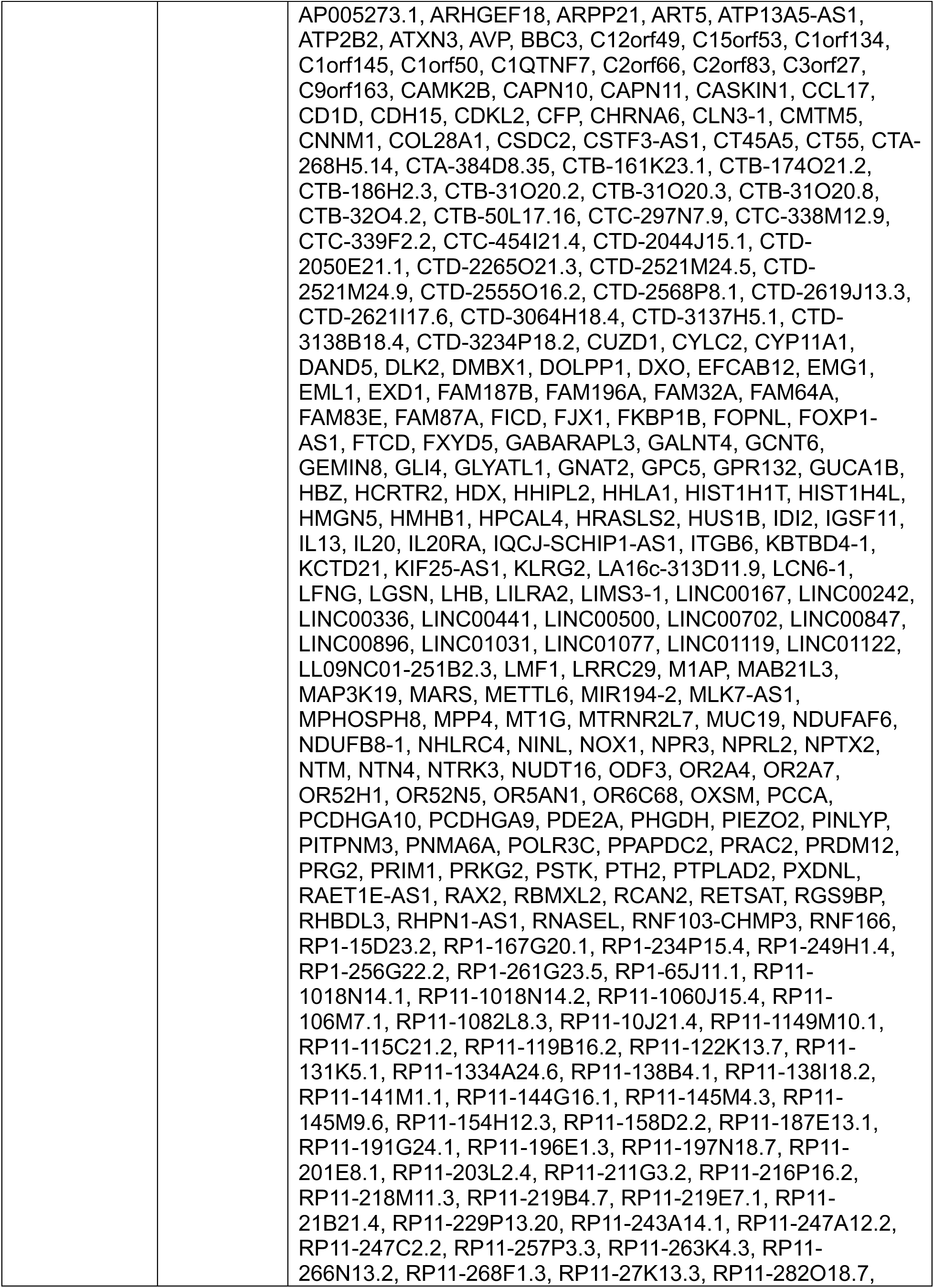
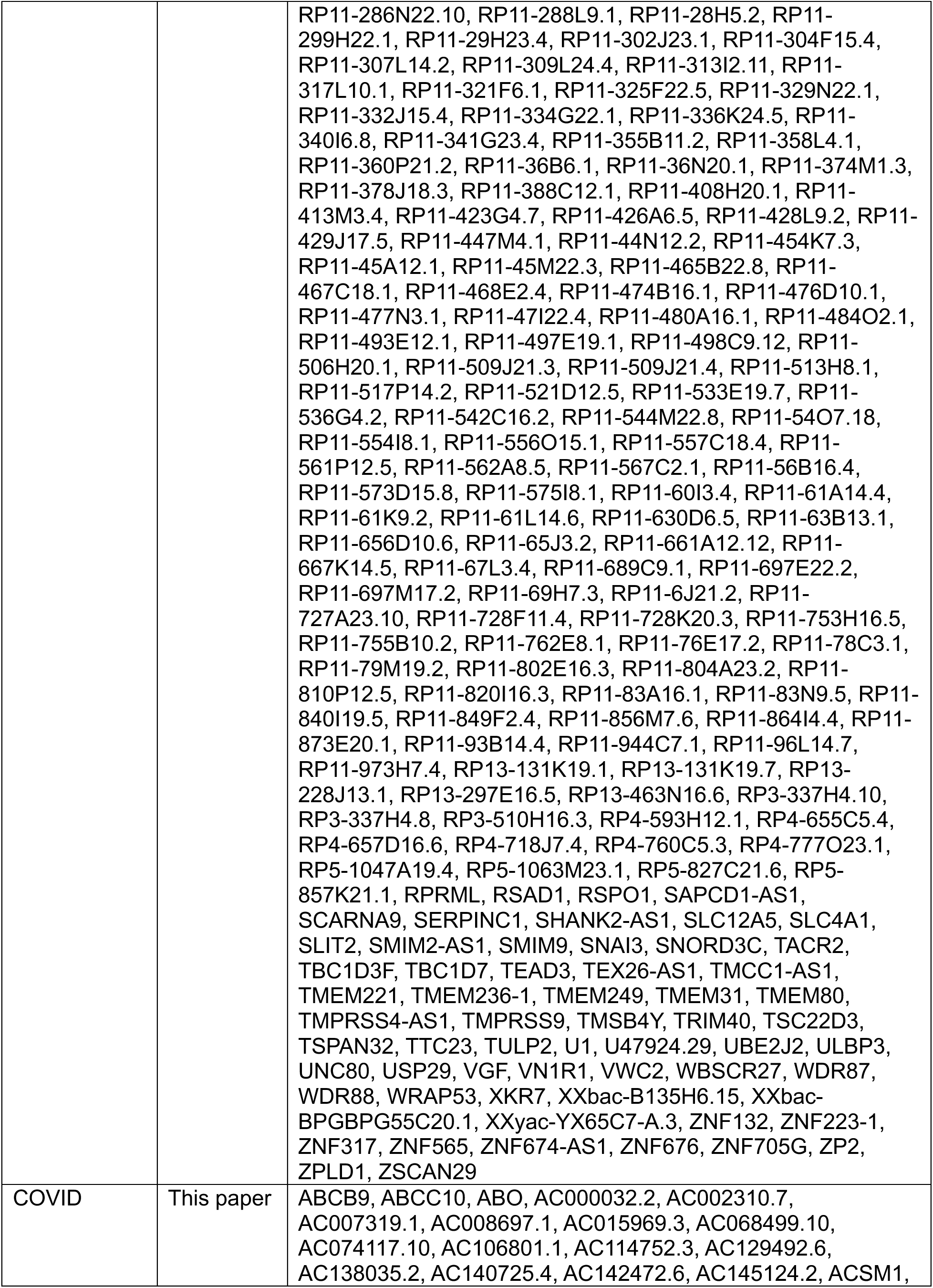
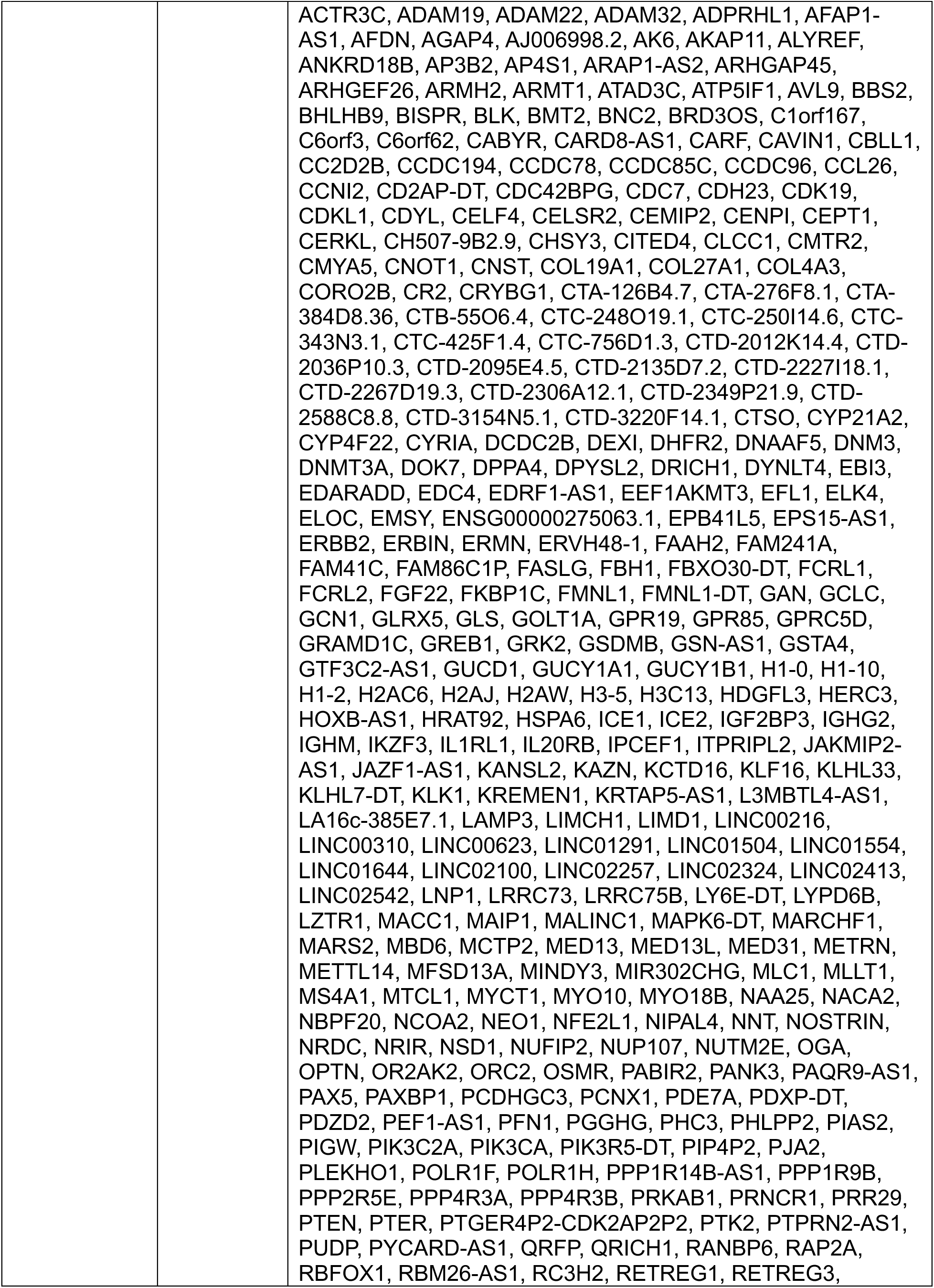
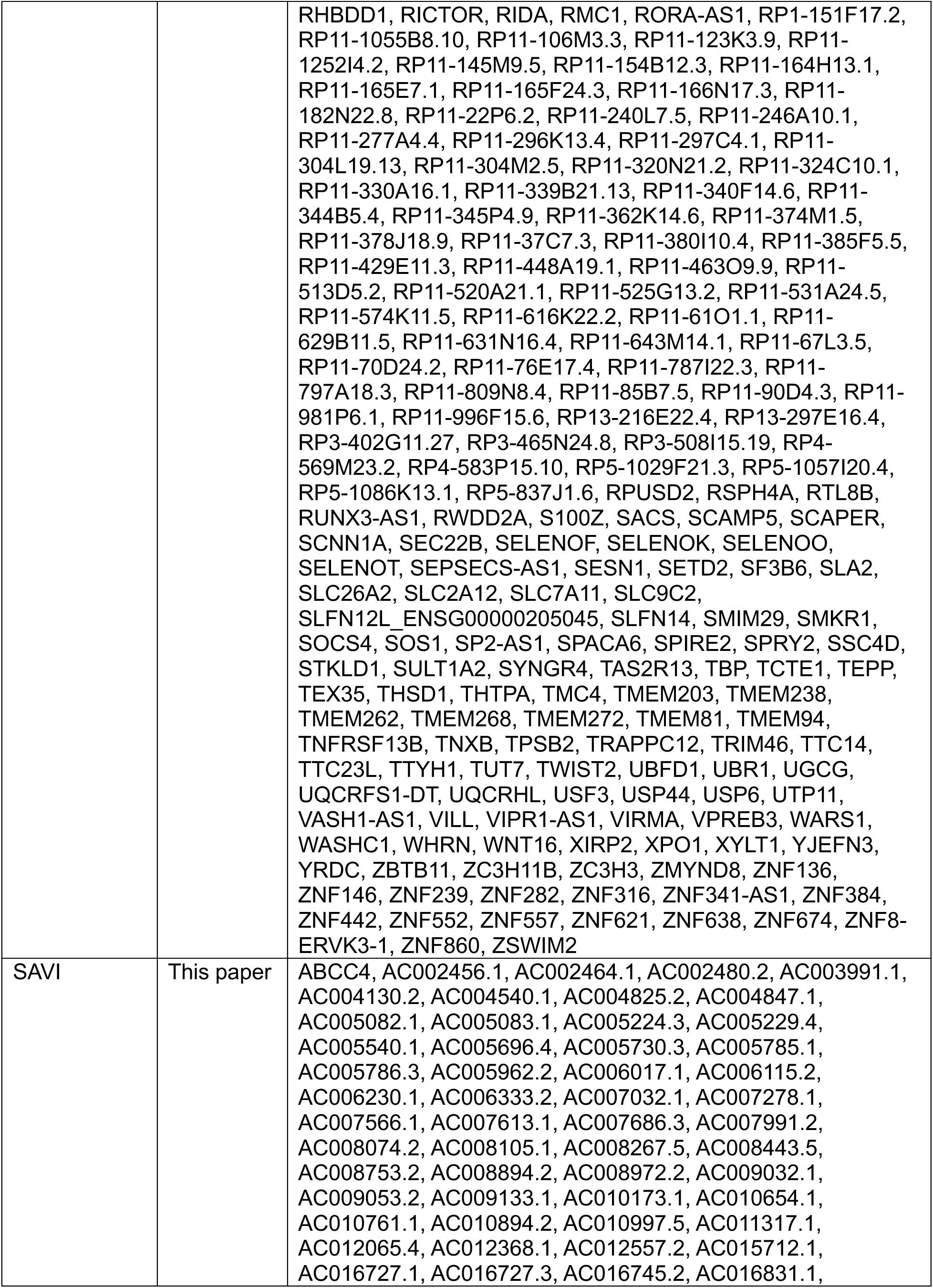
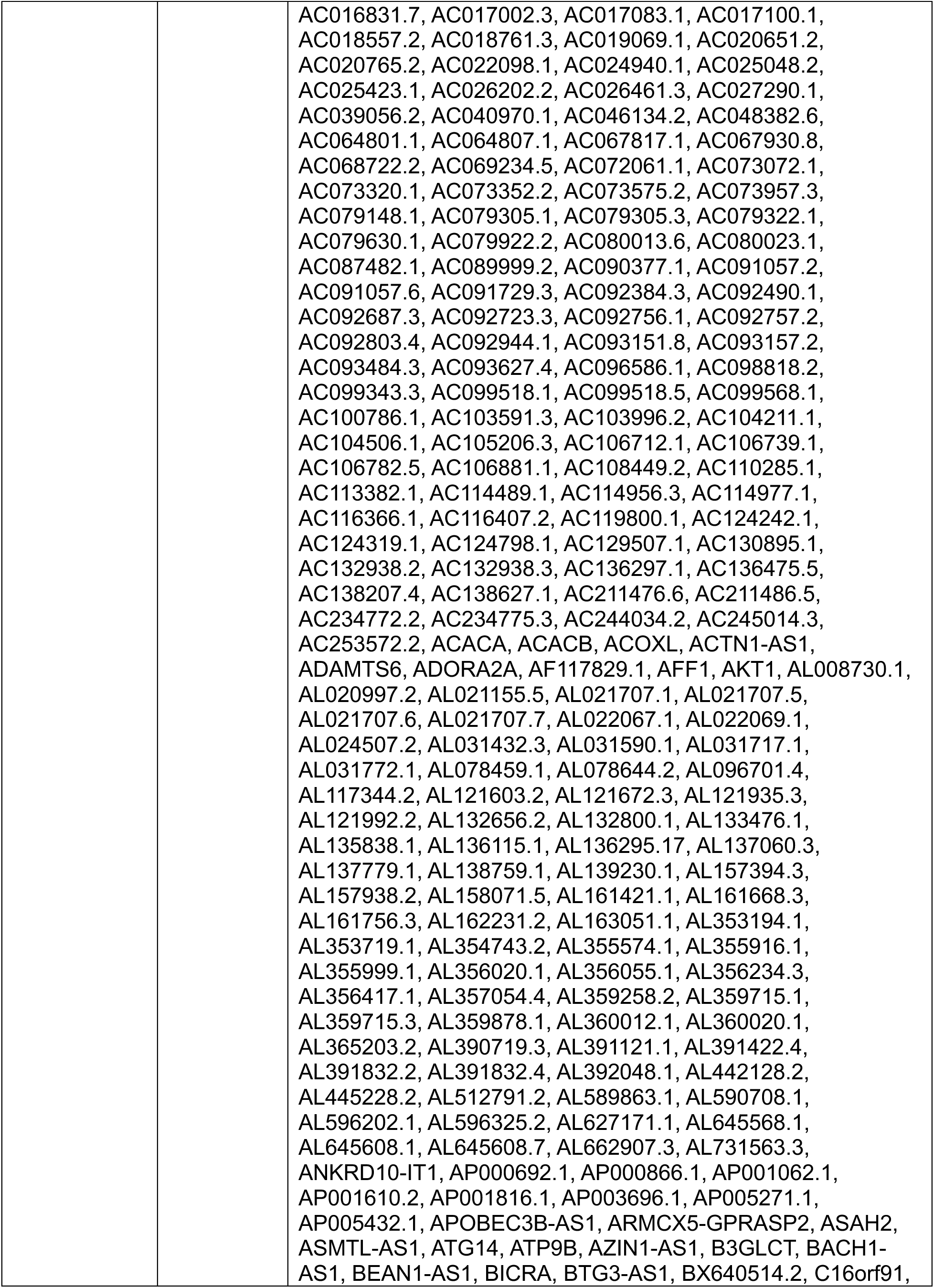
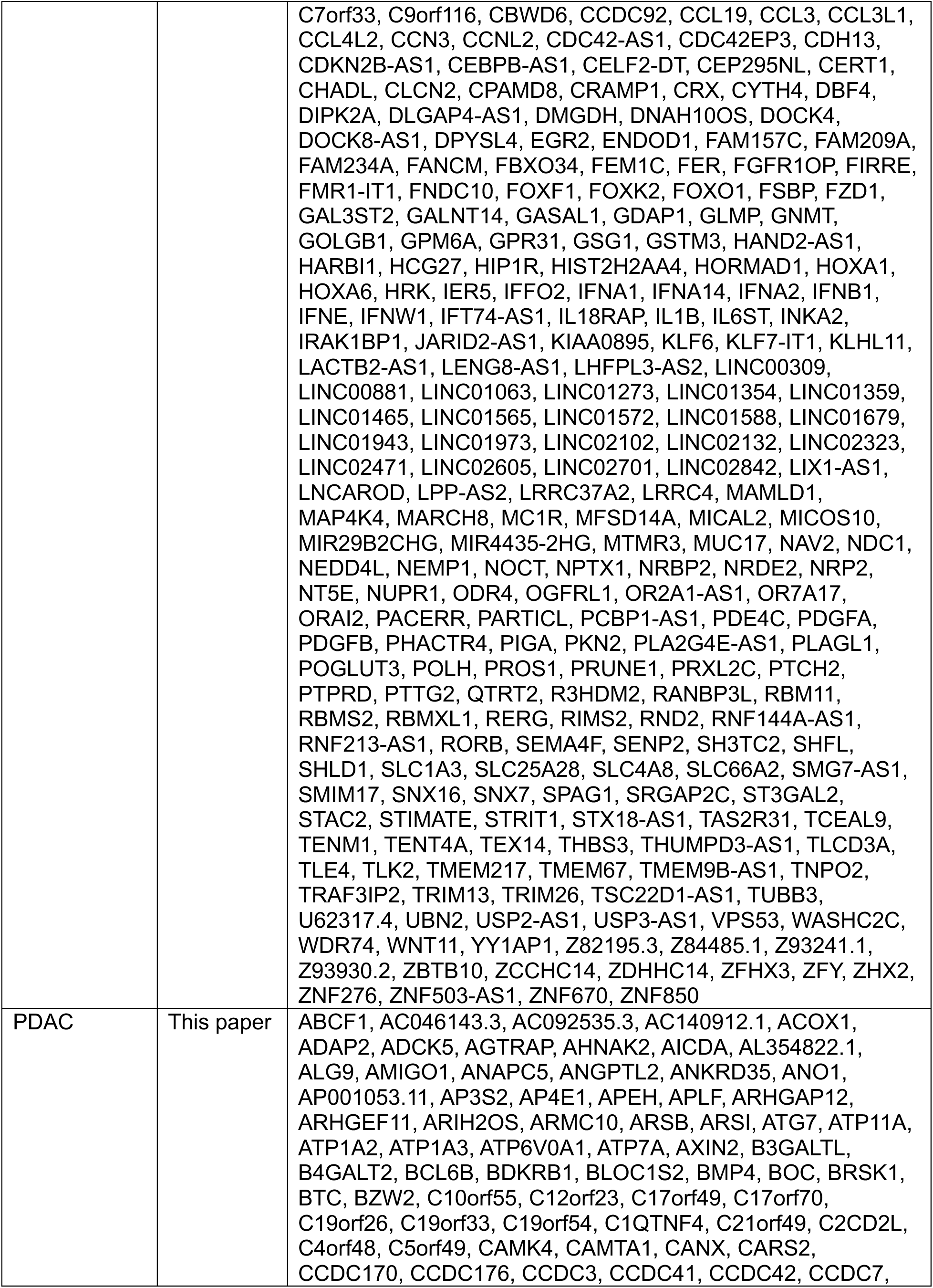
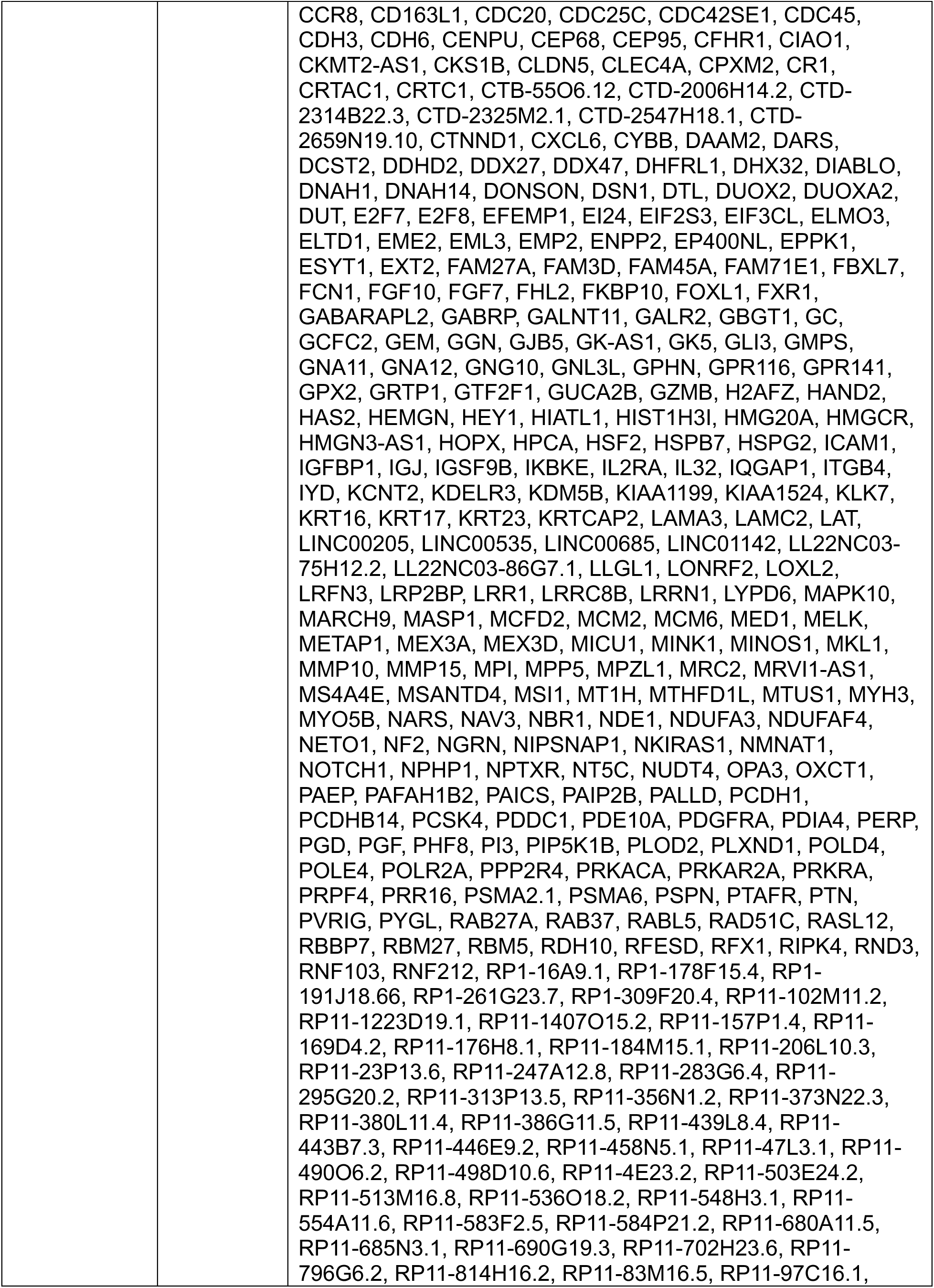
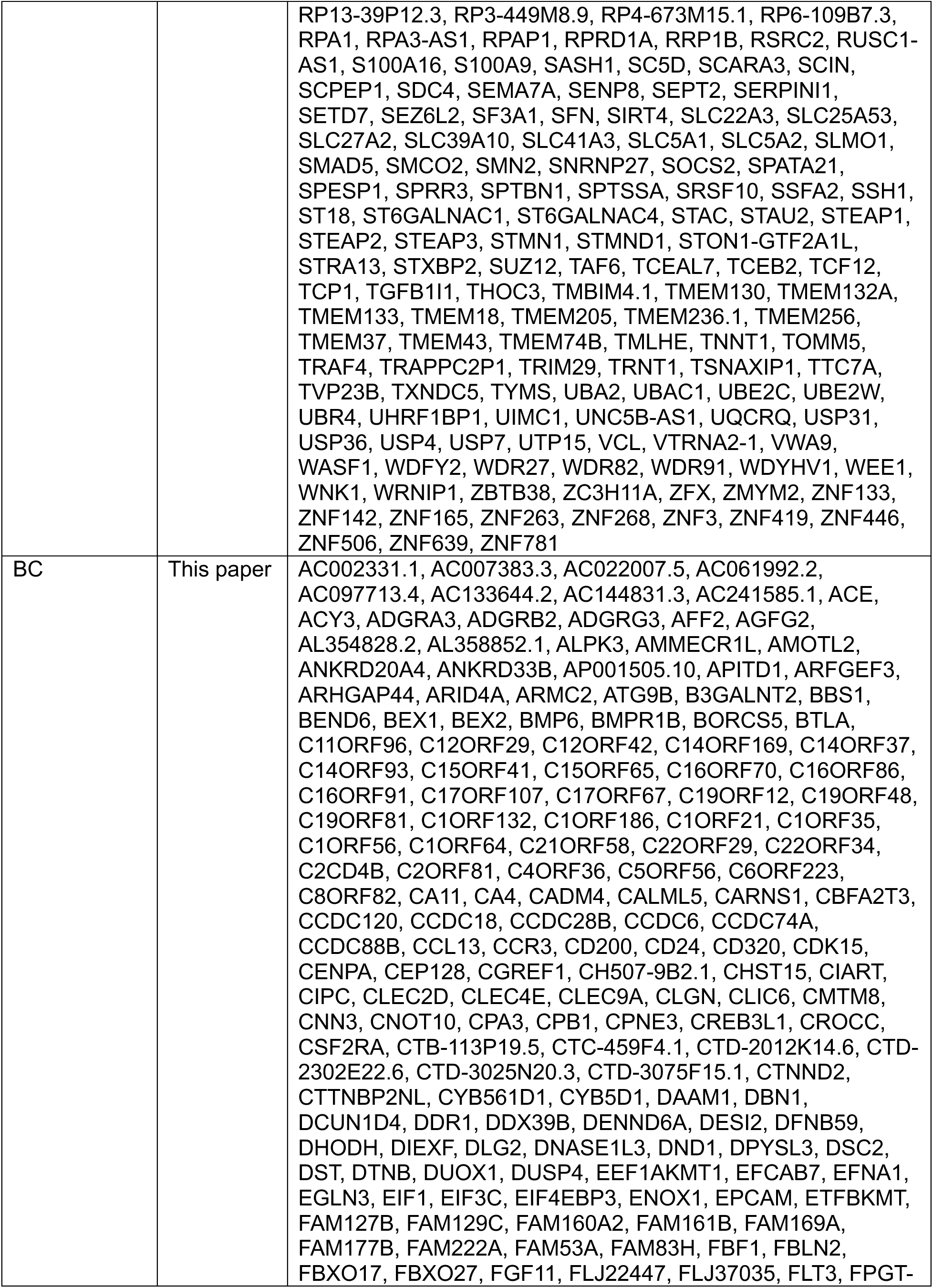
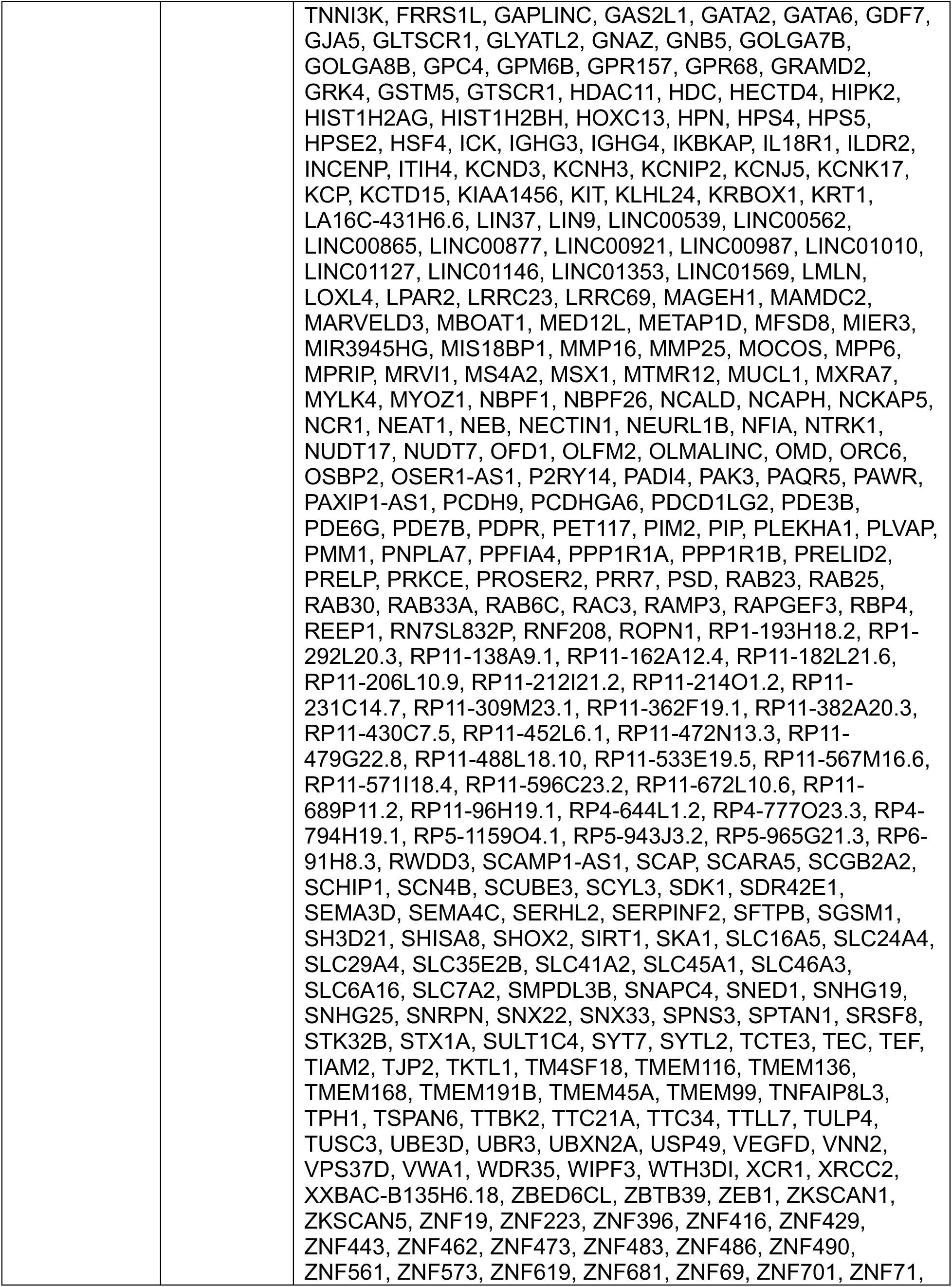
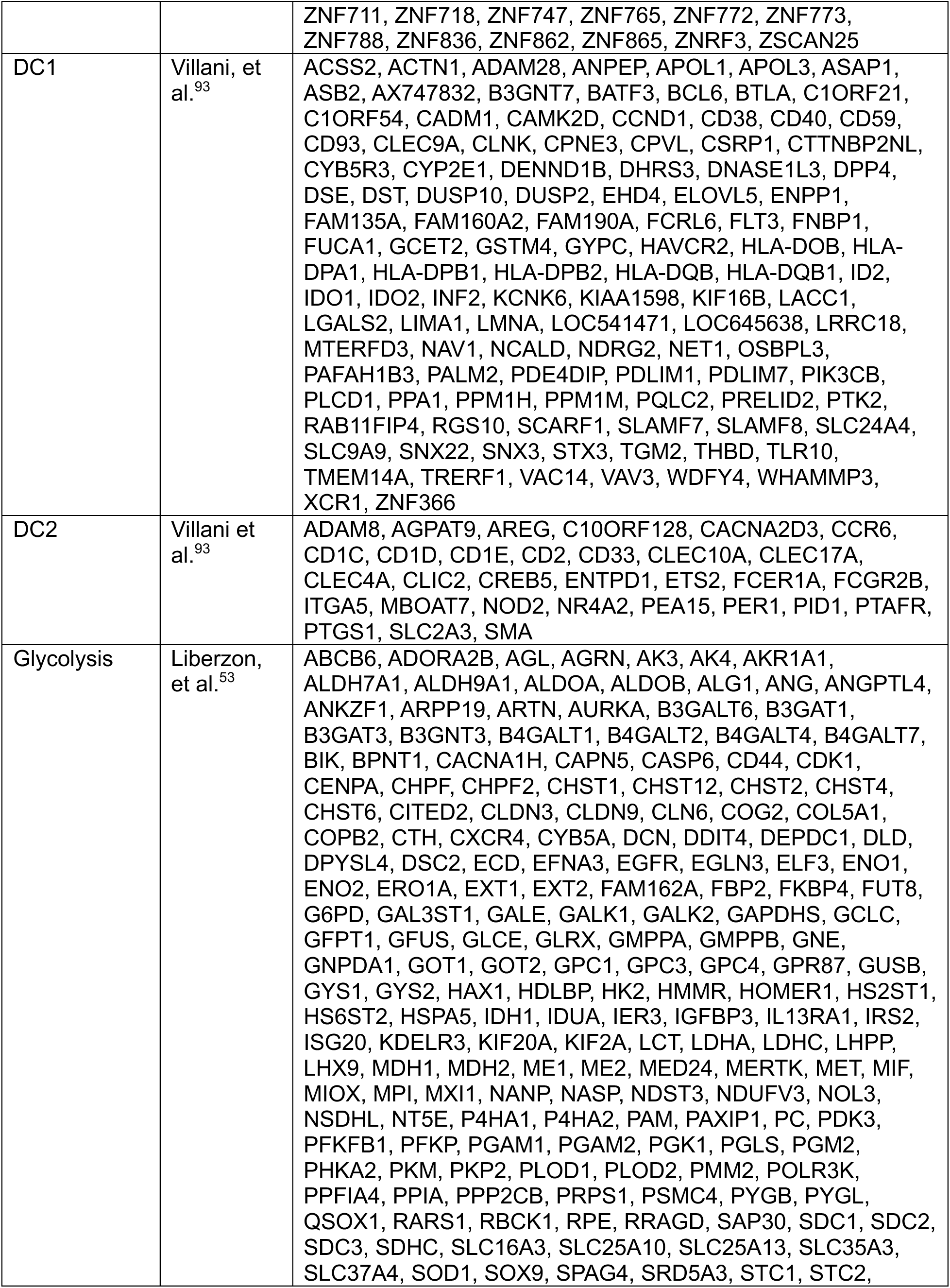
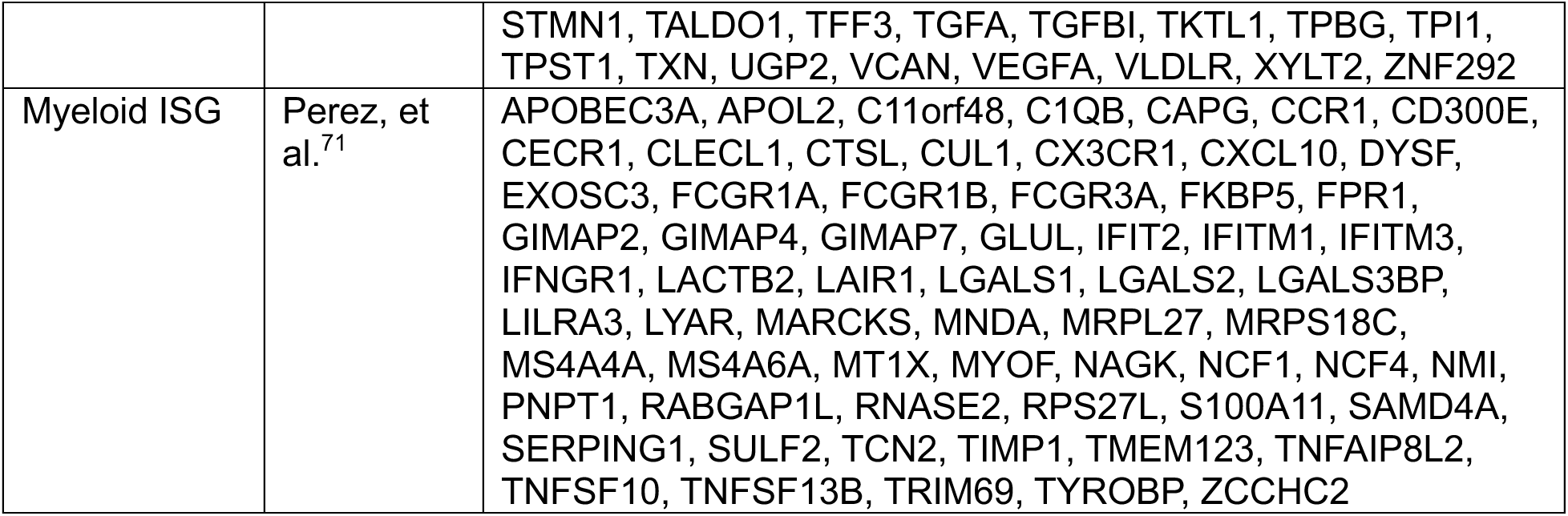
Human gene signatures used in this paper. SLE: systemic lupus erythematosus; PDAC: pancreatic ductal adenocarcinoma; BC: breast cancer; SAVI: STING-associated vasculopathy with onset in infancy; COVID: coronavirus disease 2019.

### Human disease specific cDC signature generation

To derive signatures specific to cDCs from five different human disease datasets (SLE, PDAC, BC, SAVI, and COVID), the following procedure was applied. First, for each dataset, cDCs from patients with disease were scored for the human *Inflammatory* and *IFNAR-dependent* signatures. A Wilcoxon rank sum test was then conducted between cells which were positive for both signatures (DP) versus cells which were negative for both signatures (DN). Herein, genes whose MAGIC imputed expression had an area under the curve (AUC) statistic greater than 0.75 in the DP group versus the DN group were retained. Finally, each list of genes was further pruned by removing any genes shared between two or more diseases, and leaving only those genes unique to a single disease, and then retaining the top 500 genes (ranked by AUC).

### Differentiation state and pseudotime analysis

To gauge the relative differentiation state of cells, the CytoTRACE algorithm^34^ was used, where CytoTRACE scores were computed using the CellRank library^108^. Since CytoTRACE assigns high scores to cells in the early stages of differentiation, for clarity, the CytoTRACE score was transformed and inverted to highlight terminally differentiated cells using the value 1 - CytoTRACE^2^. Separately, to conduct pseudotime analysis, the leading 50 principal components and Leiden clusters (computed as described above) were passed to the slingshot library^35^, with the starting cluster (t = 0) set to the cluster with the highest *Mki67* expression. To visualize these results, pseudotemporal trajectories were drawn atop a UMAP embedding with waypoints set at the cluster medoids.

### Integrated principal component analysis (PCA)

To gauge global transcriptomic similarities, normalized gene expression matrices from different datasets were concatenated together, restricting to genes that were shared by all datasets (after converting human genes to their murine orthologs as described above). Then, using the batchelor library^109^, the leading 50 principal components were computed from the top 2000 highly variable genes in the concatenated data, where each individual dataset was a batch in a batch-balanced PCA algorithm. Integration was then performed by adjusting these principal components using the Harmony algorithm^98^, where the theta and lambda parameters were set to 10 and 0.01, respectively. Lastly, the leading two harmonized principal components were used to visualize populations of interest, where each gated population from a dataset was represented by its respective medoid.

### Gene set enrichment analysis (GSEA)

To perform GSEA, cells within a region were compared to all other cells outside that region. Herein, the area under the curve (AUC) value derived from a Wilcoxon rank sum test (performed as described above) was used as inputs for pre-ranked GSEA using the fgsea library^96^ for all gene ontology biological process (GOBP) gene sets^92,110^ that were offspring of the tolerance induction (GO:0002507) or the immune effector response (GO:0002252) terms. GSEA plots, alongside normalized enrichment scores (NES) and adjusted p-values (FDR) were generated using the fgsea library.

### Gene ontology (GO) analysis

To perform GO analysis, the topGO library^111^ was used. Herein, individual signatures were passed to the topGO::runTest function, using the weight01 algorithm and the Fisher test, calculating enrichments for all biological process gene sets. For visualization of results, gene sets that had at least two significant genes and had a p-value less than 0.05 were displayed using lollipop plots. For murine signatures, the gene sets shown were restricted to those that were offspring of the immune system process (GO:0002376) term.

### Graphics generation

Biaxial plots and histograms were produced using the ggplot2 library^97^. Euler and Venn diagrams were generated using the eulerr library^95^. Heatmaps were produced using the ComplexHeatmap library^112^. Flow cytometry plots were prepared using FlowJo 10 (FlowJo, LLC). Bar charts and violin plots were made in GraphPad Prism 10 (GraphPad Software). The graphical abstract and various experimental schematics were generated using BioRender.

### Quantification and statistical analysis

Sex- and age-matched mice of specified genotypes were randomly assigned into individual experimental groups. Data are presented as mean ± standard deviation (SD). Group sizes were determined based on the results of preliminary experiments, with no statistical method to predetermine the sample size. When comparing only two different groups for differences, an unpaired two-tailed Welch’s *t*-test was employed. Otherwise, for pairwise comparisons involving three or more groups, Dunnett’s T3 multiple comparisons test was used. Differences were considered to be statistically significant when p < 0.05, where p*-*values were calculated in GraphPad Prism 10 (GraphPad Software).

---

**Supplemental Figure 1.**
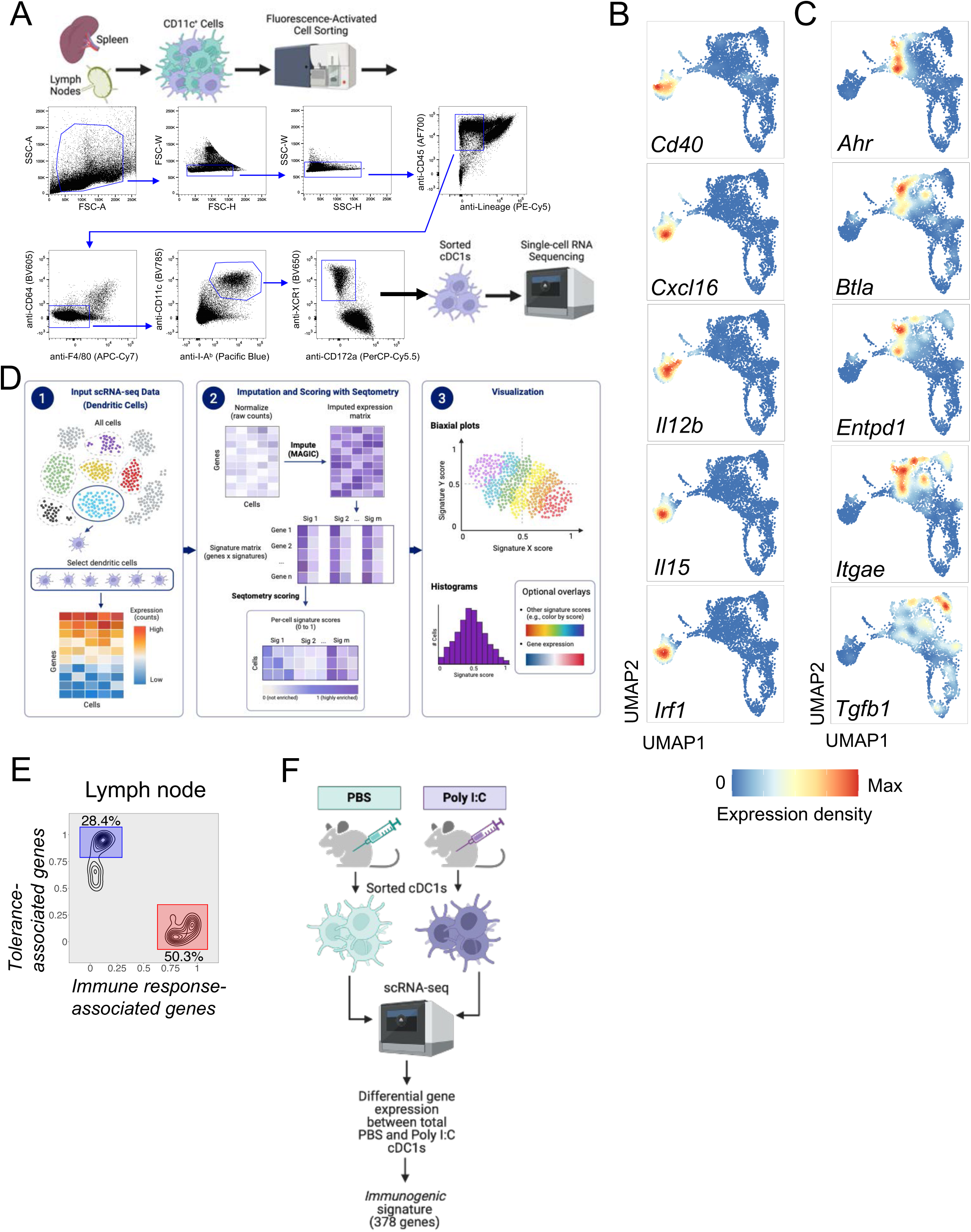
**(A)** Experimental strategy for sorting of cDC1s from spleens and peripheral lymph nodes (LNs) of WT mice for scRNA-seq. Plots show staining intensities of the indicated antibodies, and gates indicate cells selected for downstream analysis. cDC1s are identified as singlet, CD45^+^, Lineage (CD3ε, CD19, B220, NK1.1)^neg^, F4/80^neg^, CD64^neg^, I-A^b+^, CD11c^+^, XCR1^+^, CD172a^lo^. **(B)** UMAP embeddings of splenic cDC1s from mice treated with PBS overlaid with expression densities of *Immune response–associated* genes as indicated. **(C)** UMAP embeddings of splenic cDC1s from mice treated with PBS overlaid with expression densities of *Tolerance–associated* genes as indicated. **(D)** Schematic showing workflow for Seqtometry analysis. **(E)** Plot showing distributions of scores for indicated signatures among cDC1s from peripheral lymph nodes (LNs). Regions indicate *Tolerance-associated*^pos^ (blue) and *Immune response–associated*^pos^ (red) populations used for subsequent analysis. Numbers next to regions indicate corresponding percentages. **(F)** Experimental schematic for derivation of the *Immunogenic* signature.

**Supplemental Figure 2.**
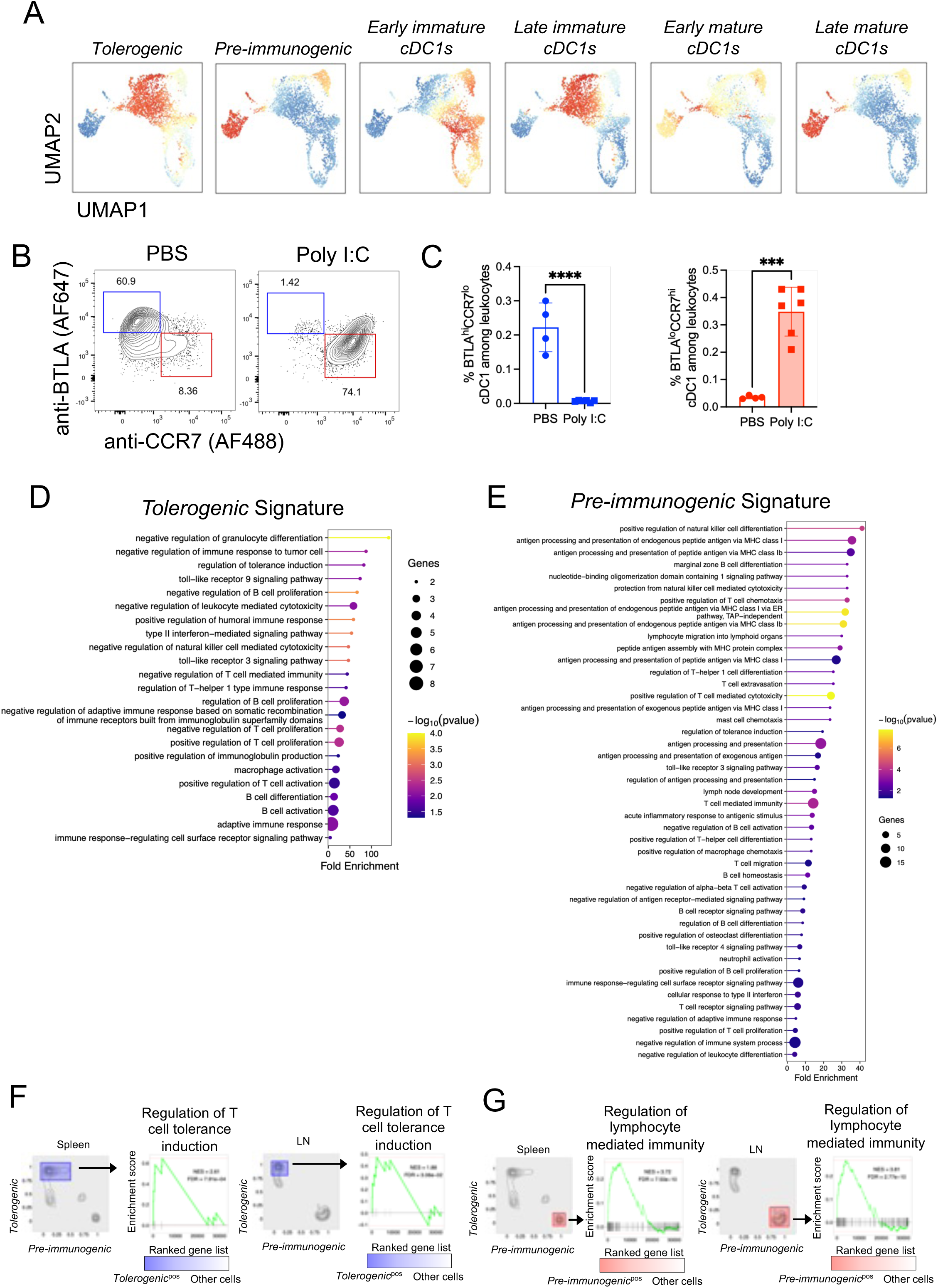
**(A)** UMAP embeddings of splenic cDC1s from mice treated with PBS overlaid with color overlay indicating the indicated signatures, including those from Bosteels, et al.^17^ **(B)** Plots (representative of three independent experiments) showing the staining intensity of the indicated fluorochrome-conjugated antibodies among all splenic cDC1s from wild-type mice treated with PBS or poly I:C 12 hours before analysis by flow cytometry. cDC1s were gated as in Supplemental Figure 1A. Regions and corresponding percentages indicate BTLA^hi^CCR7^lo^ (blue) and BTLA^lo^CCR7^hi^ (red) populations. **(C)** Graphs show percentages of BTLA^hi^CCR7^lo^ cDC1 (left, blue) or BTLA^lo^CCR7^hi^ (right, red) among all splenic cDC1s in the indicated treatment groups, gated as in Figure S2B (n = 4-6 mice per group, pooled from three experiments). Error bars represent SD. ** P< 0.01 and **** P< 0.0001, determined by unpaired two-tailed *t*-test. **(D)** Gene ontology (GO) analysis of the *Tolerogenic* signature. **(E)** Gene ontology (GO) analysis of the *Pre-immunogenic* signature **(F and G)** Plots showing distributions of scores for *Pre-immunogenic* and *Tolerogenic* signatures among cDC1s from spleens and peripheral LNs as indicated. Regions indicate *Tolerogenic*^pos^ (blue) populations (F) and *Pre-immunogenic*^pos^ (red) populations (G) used for GSEA analysis. Corresponding GSEA plots show enrichment of indicated biological processes comparing *Tolerogenic*^pos^ populations (F) and *Pre-immunogenic*^pos^ populations (G) and to all other cDC1s for spleens and LNs. NES: normalized enrichment score. FDR: false discovery rate.

**Supplemental Figure 3.**
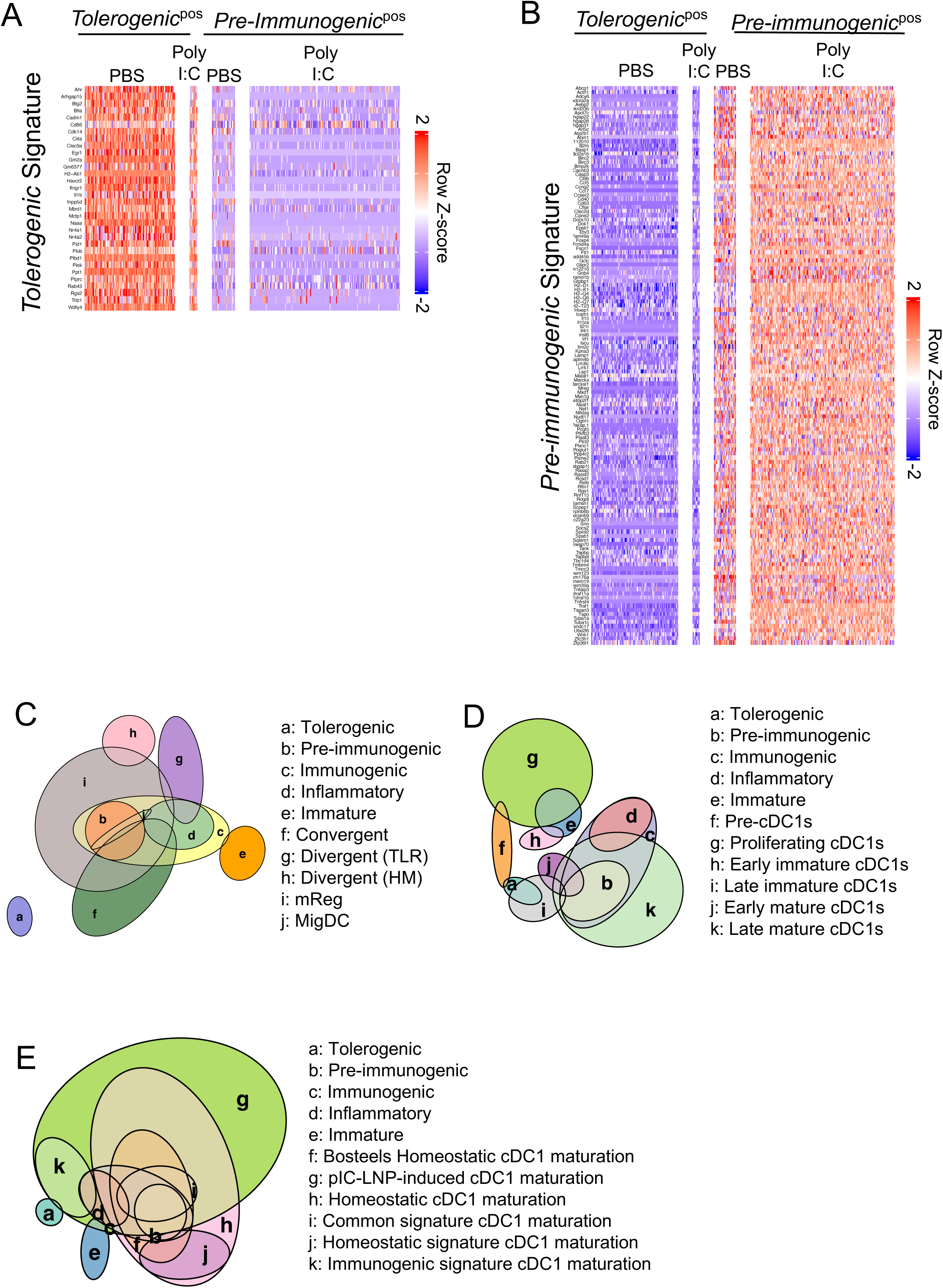
**(A)** Heatmap of genes of the *Tolerogenic* signature for cDC1s from spleens of WT mice treated with PBS or poly I:C, with populations gated as in Figure 1I. **(B)** Heatmap of genes of the *Pre-immunogenic* signature for cDC1s from spleens of WT mice treated with PBS or poly I:C, with populations gated as in Figure 1I. **(C-E)** Euler diagrams showing relative overlaps of *Tolerogenic*, *Pre-immunogenic*, and *Immunogenic* signatures. The *Immature* signature was derived by comparing all cells with a low differentiation score (1-CytoTRACE^2^ < 0.2) to all other cells (as shown in Figure 1B). *Convergent*, *Divergent (TLR*), and *Divergent (HM)* signatures were obtained from^22^. *mReg* (mature DCs enriched in immunoregulatory molecules) gene set was obtained from^23^. *migDC* (migratory DC) gene set was obtained from^91^. TLR: Toll-like receptor, HM: homeostatic maturation. *Proliferating cDC1s*, *Early immature cDC1s*, *Late immature cDC1s*, *Early mature cDC1s*, *Late mature cDC1s* signatures were obtained from Bosteels, et al.^17^ *Homeostatic cDC1 maturation*, *pIC-LNP-induced cDC1 maturation*, *Homeostatic cDC1 maturation*, *Common signature cDC1 maturation*, *Homeostatic signature cDC1 maturation*, *Immunogenic signature cDC1 maturation* were obtained from Rennen, et al.^36^

**Supplemental Figure 4.**
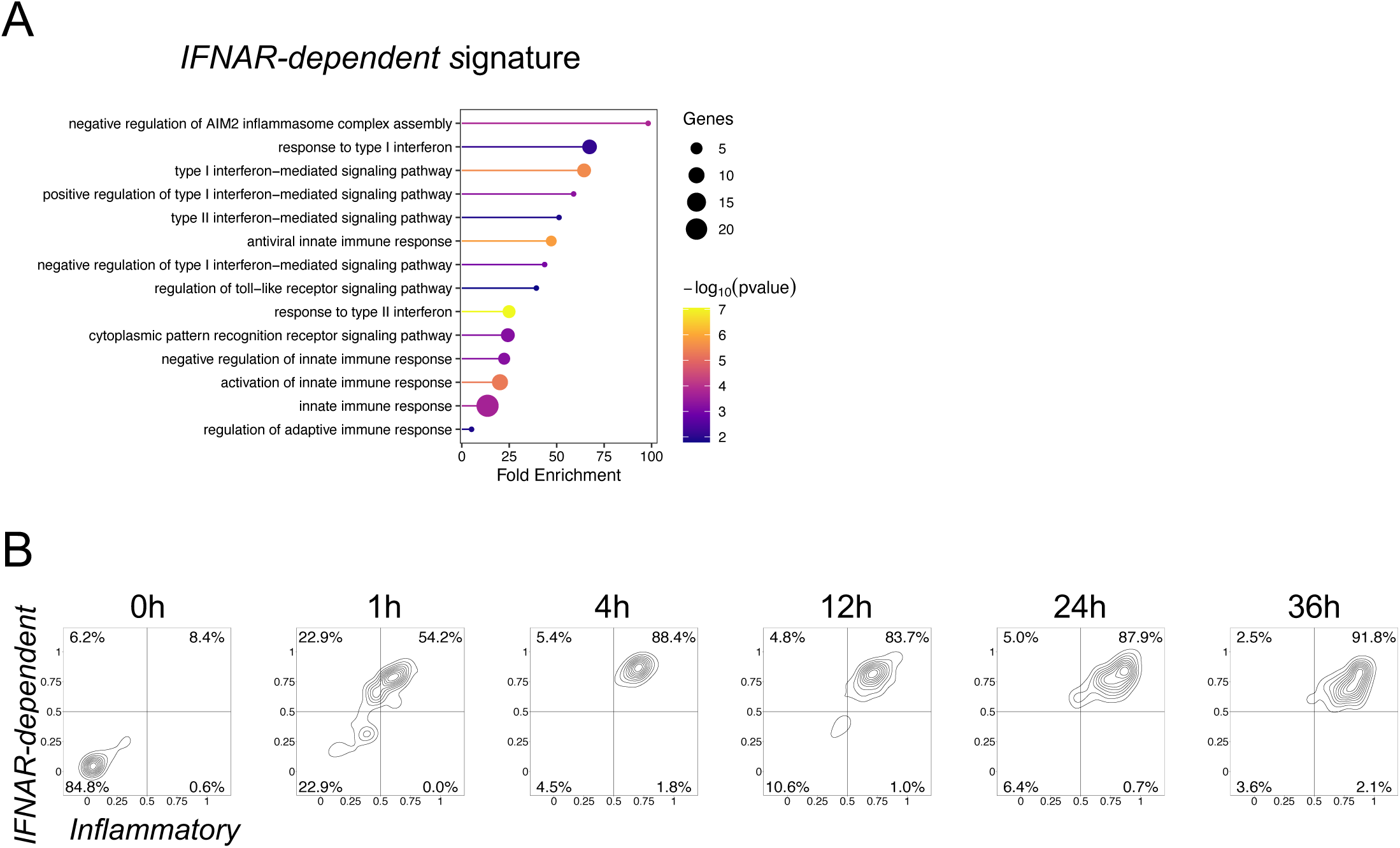
**(A)** Gene ontology (GO) analysis of the *IFNAR-dependent* signature. **(B)** Data obtained from de Cevins, et al.^64^ Plots showing distributions of scores of indicated signatures among cDCs at specific time points from i*n vitro* cultures with IFN-β.

**Supplemental Figure 5.**
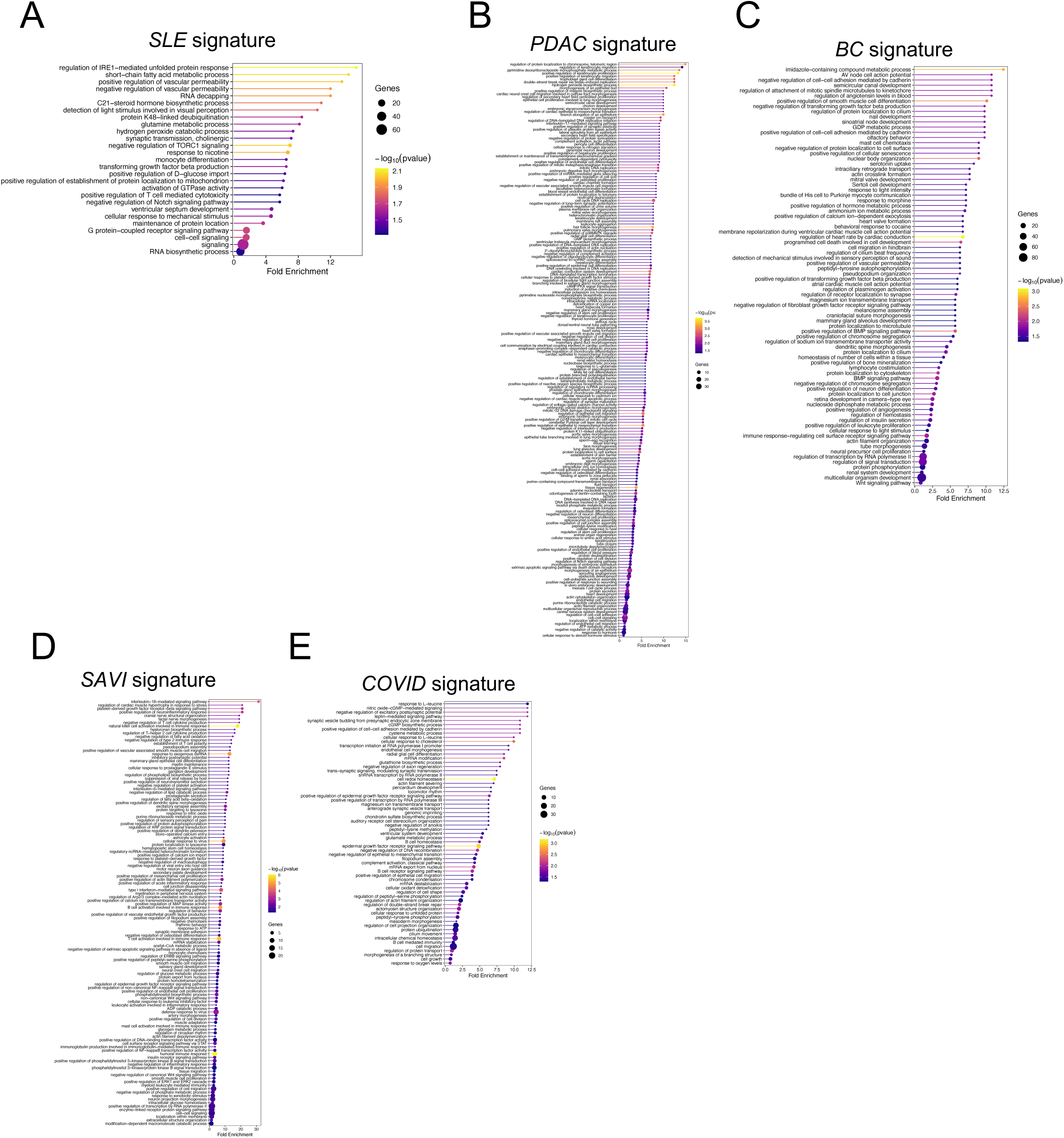
**(A–E)** Data for individual diseases were obtained from van der Wijst, et al.,^76^ de Cevins, et al.,^64^ Perez, et al.,^71^ Peng, et al.,^77^ and Azizi, et al.^78^ HC: healthy control; SLE: systemic lupus erythematosus; PDAC: pancreatic ductal adenocarcinoma; BC: breast cancer; SAVI: STING-associated vasculopathy with onset in infancy; COVID: coronavirus disease 2019. GO analysis of the indicated disease-specific signatures.

